# Semaphorin 4A maintains functional diversity of the hematopoietic stem cell pool

**DOI:** 10.1101/2024.11.12.622506

**Authors:** Dorsa Toghani, Sanika Gupte, Sharon Zeng, Elmir Mahammadov, Edie I. Crosse, Negar Seyedhassantehrani, Christian Burns, David Gravano, Stefan Radtke, Hans-Peter Kiem, Sonia Rodriguez, Nadia Carlesso, Amogh Pradeep, Alexis Georgiades, Fabienne Lucas, Nicola K. Wilson, Sarah J. Kinston, Berthold Göttgens, Le Zong, Isabel Beerman, Bongsoo Park, Derek Janssens, Daniel Jones, Ali Toghani, Claus Nerlov, Eric Pietras, Marion Mesnieres, Christa Maes, Atsushi Kumanogoh, Thomas Worzfeld, Jin-Gyu Cheong, Steven Z Josefowicz, Peter Kharchenko, David T. Scadden, Antonio Scialdone, Joel A Spencer, Lev Silberstein

## Abstract

Somatic stem cell pools are comprised of diverse, highly specialized subsets whose individual contribution is critical for the overall regenerative function. In the bone marrow, myeloid-biased HSC (myHSC) are indispensable for replenishment of myeloid cells and platelets during inflammatory response but at the same time, become irreversibly damaged during inflammation and aging. Here, we identify an extrinsic factor, Semaphorin 4A (Sema4A), which non cell-autonomously confers myHSC resilience to inflammatory stress. We show that the absence of Sema4A, myHSC inflammatory hyper-responsiveness in young mice drives excessive myHSC expansion, myeloid bias and profound loss of regenerative function with age. Mechanistically, Sema4A is mainly produced by neutrophils, signals via a cell surface receptor Plexin D1 and safeguards myHSC epigenetic state. Our study shows that by selectively protecting a distinct stem cell subset, an extrinsic factor preserves functional diversity of somatic stem cell pool throughout organismal lifespan.

## INTRODUCTION

Cellular diversity is a fundamental feature of complex organisms because the presence of specialized cell types – i.e., those that are more adept to performing a specific function – is beneficial for adaptability to environmental challenge and organismal survival.^1^ Until recently, cellular diversity was considered a sole property of terminally differentiated cells. However, recent studies suggest that in multiple tissues (brain, skeletal muscle, cornea, bone marrow), the most primitive cells – somatic stem cells – are also highly heterogenous and contain specialized subsets which are specifically “designed” to drive tissue regeneration in specific physiological contexts.^1–5^ While each of these subsets makes a unique contribution to tissue regenerative function, little is known about the signals which selectively maintain their abundance and functional competence.

In the bone marrow, myeloid-biased HSC (myHSC) are a specialized HSC subset which is tasked with replenishment of mature myeloid cells and platelets during inflammatory response.^6–13^ MyHSC, which have been found in both mice and humans ^14, 15^, are exquisitely responsive to acute and chronic inflammatory stimuli and primed for generation of myeloid/platelet progeny via myeloid-bypass hematopoiesis. In addition, myHSC are emerging as effectors of trained immunity, raising a possibility that they serve as a long-lasting reservoir of inflammatory memory.^16^ Thus, myHSC are critical for host defense and organismal response to inflammation.

Inflammatory stress causes irreversible loss of HSC function.^17^ Hence, in order to perform their specialized role as “first-line” inflammation responders, myHSC must possess specific mechanisms which mediate their resilience to inflammatory stress. Failure of such mechanisms is exemplified by aging, in which the loss of the ability to withstand chronic inflammatory signals leads excessive myHSC expansion and myeloid bias, which in turn account for immune dysfunction and increased risk of myeloid malignancy with age. ^18^ ^19^ Although the existence of regulatory signals which selectively regulate myHSC response to stress has been suggested by prior studies,^11, 20, 21^ only few have been identified to date and their functional significance, particularly in the context of inflammation and aging, has not been defined.

In the current study, we identify as a extrinsic factor, Semaphorin 4A (Sema4A),^22–24^ as a novel inhibitory signal which selectively attenuates myHSC activation during inflammation and preserves function while having little/no measurable effect on the other HSC subset - lymphoid-biased HSC (lyHSC). In the absence of Sema4A, myHSC display enhanced responsiveness to tonic and induced inflammation and excessively expand with age, giving rise to exaggerated myeloid bias, attrition of myHSC regenerative function and an overall accelerated aging-like hematopoietic phenotype. Sema4A acts in a non cell-autonomous manner via binding to a cell surface receptor, PlxnD1^25^, and originates predominantly from mature neutrophils. Collectively, these findings define Sema4A/PlxnD1 as a previously unknown myHSC-protective regulatory axis. More generally, they demonstrate that functional diversity of a somatic stem cell pool is maintained by a cell-extrinsic signal, which is responsible for preserving a specialized stem cell subset in a specific physiological context.

## RESULTS

### The absence of Sema4A results in premature hematopoietic aging-like phenotype

We have previously performed an *in vivo* screen for novel cell-extrinsic regulators of HSC function. Briefly, we compared single cell transcriptional signatures of stromal cells located in close proximity to transplanted HSC and those further away, and prioritized extrinsic factors with higher expression in the former as candidates for functional validation.^26, 27^. Several of these molecules were found to act as extrinsic regulators of HSC self-renewal.^28, 29^ Among those not yet characterized, we decided to focus on Sema4A – a membrane-bound and secreted protein with an established role in neurogenesis, angiogenesis and immune response ^25, 30, 31^ but no known function in hematopoiesis.

In the course of phenotypic analysis of Sema4AKO mice, we unexpectedly noticed that by mid-age (44 weeks), these animals developed progressive neutrophilia and thrombocytosis, as well as anemia when they became fully aged (74-80 week old). (Fig 1A-C). Histological examination of the bone marrow in aged Sema4AKO mice revealed a relative reduction of erythroid cells due to an expansion of myelopoieisis (myeloid hyperplasia), leading to increased M:E ratio (Fig 1E, F). Myeloid hyperplasia also manifested as focal crowding of hematopoietic elements into the sub-epiphyseal fat space (Fig 1D, Extended Data Fig 1A), similar to the bone marrow findings in patients with myeloproliferative disorders ^32^. These features represent hallmarks of hematopoietic aging ^33^ but were exaggerated in aged Sema4AKO mice, suggesting the absence of Sema4A resulted in premature aging-like phenotype. In keeping with this notion, immunophenotypic analysis of the bone marrow using fluorescence-activated cell sorting (FACS) showed that aged Sema4A mice displayed significantly increased absolute numbers of HSC, myeloid and megakaryocyte progenitors but decreased erythroid progenitors. In addition, we observed expansion of mature myeloid cells but contraction of the B-cell compartment (Fig 1G-J). These changes were due to a difference in frequencies of respected cellular subsets between aged WT/Sema4AKO mice, since the bone marrow cellularity was comparable (Extended Data Fig 1B).

**Fig 1.**
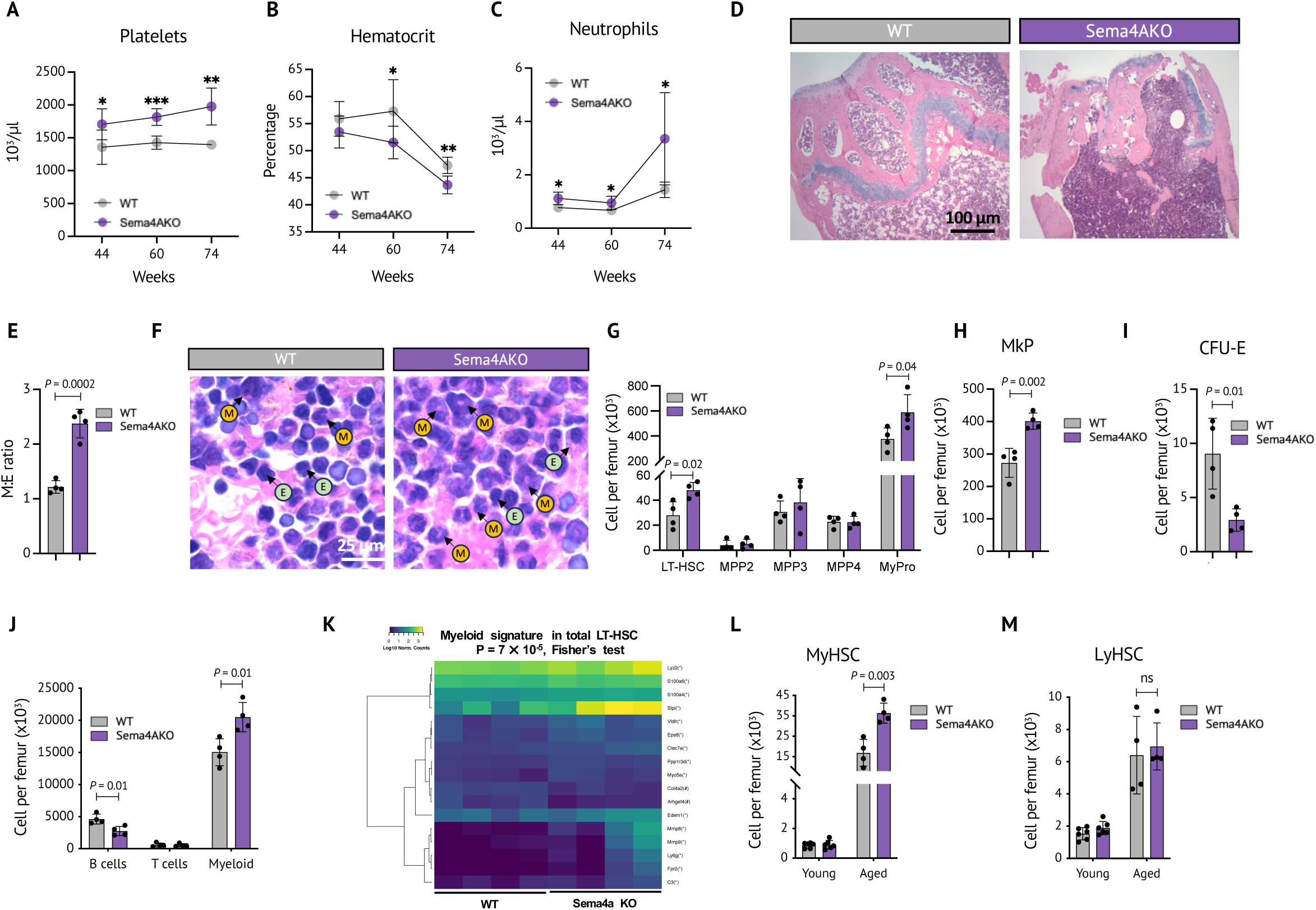
The absence of Sema4A leads to excessive myeloid expansion and premature hematopoietic aging-like phenotype. **A-C.** Platelet count **(A)**, hematocrit level **(B)**, and neutrophil count **(C)** in the peripheral blood of aged WT/Sema4AKO mice (n=5 mice per group). **D.** Representative images of sub-epiphyseal area of H&E stained femurs from aged WT/Sema4AKO mice. **E.** Myeloid to erythroid ratio in the bone marrow of aged WT/Sema4AKO mice (n=4 mice per group). **F.** Representative images of H&E staining of femurs from aged WT/Sema4AKO mice. Arrows with letters M and E indicate myeloid and erythroid cells respectively. **G-J.** Absolute number of hematopoietic stem and progenitor subsets **(G),** megakaryocyte progenitors (MkP) **(H)**, erythroid progenitors (colony-forming unit – erythroid, CFU-E) **(I)** and mature cells **(J)** in the bone marrow of aged WT/Sema4AKO mice (n=4-5 mice per group). **K.** Projection of previously published myeloid signature onto the RNA-Seq profiles of HSC from aged WT/Sema4A (n=4 animals per group). **L, M.** Absolute numbers of myHSC **(L)** and LyHSC **(M)** in the bone marrow of aged WT/Sema4AKO mice (n=5 mice per group). P values are shown. For the peripheral blood counts, *P<0.05, **P<0.01, ***P<0.001. Statistical significance was assessed by two-tailed *t*-test. Mean +/- (SEM) are shown.

To understand the molecular basis for the enhanced myelopoiesis in aged Sema4AKO mice, we performed bulk RNA sequencing (RNA-Seq) analysis of phenotypic HSC (Lin^-^Kit^+^Sca^+^CD48^-^ Flk2^-^ CD150^+^) from aged WT and Sema4AKO mice. Interestingly, these experiments showed that aged Sema4AKO HSC transcriptome was significantly enriched for a transcriptional signature associated with myeloid differentiation ^34^, suggesting that enhanced myelopoiesis in aged Sema4AKO animals originates at the HSC level (Fig 1K). This finding could be explained by either enhanced myeloid differentiation of the entire aged Sema4AKO HSC compartment or a compositional change, such as a higher relative proportion of myeloid-biased HSC within aged Sema4AKO HSC pool ^11^.

To distinguish between these scenarios, we next sought to quantify myHSC in aged WT/Sema4AKO mice using their previously established immunophenotypic definition as Lin^-^ Kit^+^Sca^+^CD48^-^Flk2^-^CD150^high^ (see Extended Data Fig 1 C,D for the gating strategy). In order to verify that this definition remains valid in the absence of Sema4A, we crossed Sema4AKO mice with vWF-Tomato reporter strain, since Lin^-^Kit^+^Sca^+^CD48^-^Flk2^-^ vWF^high^ HSC (i.e. isolated independently of CD150 expression level) have been shown in functional studies to display enhanced platelet/myeloid differentiation, both in young and aged animals^35^. We found that in young and aged WT mice, Lin^-^Kit^+^Sca^+^CD48^-^Flk2^-^**vWF^high^** HSC were highly enriched for Lin^-^ Kit^+^Sca^+^CD48^-^Flk2^-^**CD150^high^**HSC, in keeping with the published data ^35^, and that the level of vWF expression and the degree of the enrichment (∼80%) were unchanged in young and aged Sema4AKO mice (Extended Data Fig 1 E-H). Collectively, these data demonstrate that in the absence of Sema4A, CD150^high^ serves as a reliable immunophenotypic marker for myHSC.

Our subsequent analysis of the aged WT/Sema4AKO cohort showed that in WT mice, myHSC significantly expanded with age whereas the number of lyHSC (Lin^-^Kit^+^Sca^+^CD48^-^Flk2 CD150^low^) only modestly increased (Extended Data Fig 1G), consistent with prior studies^10^.

Strikingly, in aged Sema4AKO mice, expansion of immunophenotypic myHSC was markedly exaggerated; in contrast, the number of lyHSC was comparable between the groups (Fig 1L, M, Extended Data Fig 1 I,J). Taken together, these results suggest that Sema4A selectively regulates myHSC behavior during aging, and that excessive myHSC expansion in aged Sema4AKO mice may account for the premature aging-like hematopoietic phenotype.

### The absence of Sema4A during aging leads to profound loss of myHSC regenerative capacity but does not affect lyHSC

We next asked if in addition to selectively controlling myHSC *in situ* pool size during aging, Sema4A also regulates their regenerative function. To this end, we isolated CD150^high^ myHSC and CD150^low^ lyHSC from aged WT/Sema4AKO mice (both carrying CD45.2 congenic marker) and transplanted them into lethally irradiated Bl6 SJL recipient mice (CD45.1) using bone marrow cells from young CD45 1/2 mice as competitors. As a quality control, we first checked if aged WT CD150^high^ HSC exhibit post-transplant myeloid bias and indeed found that the myeloid/lymphoid ratio in the recipients of WT myHSC was significantly higher than in WT lyHSC (Extended Data Fig 2A, B).

Next, we examined post-transplant reconstitution potential of myHSC and lyHSC in aged WT/Sema4AKO mice using a competitive transplantation assay. To this end, equal numbers (2000 cells per mouse) of aged myHSC or lyHSC were transferred into lethally irradiated recipients, together with young WT bone marrow competitor cells. Strikingly, quantification of donor-derived cells in peripheral blood showed that while aged Sema4AKO lyHSC retained their competitive fitness, it was almost completely lost in aged Sema4AKO myHSC, despite a high number of these cells transplanted (Fig 2A-F, Extended Data Fig2C). We note that although the level of donor-derived cells in the recipients of aged Sema4AKO myHSC was very low, the reconstitution was long-term and multilineage, indicating that immunophenotypic aged Sema4AKO myHSC contained *bone fide* HSC (Fig 2C). In keeping with the findings in peripheral blood, bone marrow analysis at week 24 showed that in the recipients of aged Sema4AKO myHSC, donor-derived cells were almost undetectable, whereas in the recipients of aged Sema4AKO lyHSC their percentage was comparable to the controls (Fig 2 G,H).

**Fig 2.**
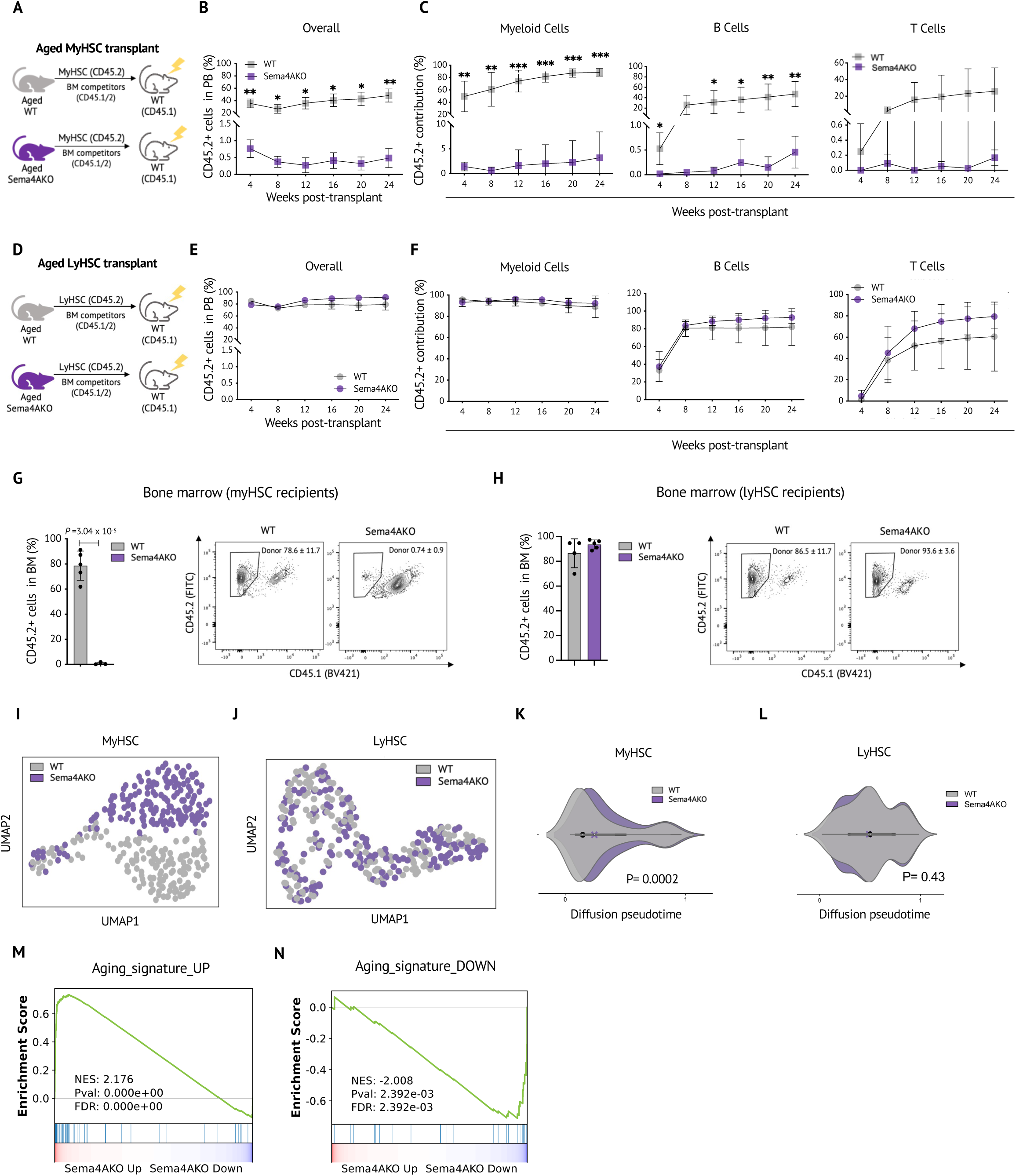
The absence of Sema4A leads to functional attrition of myHSC during aging. **A-C.** Experimental schema **(A)**, overall percentage of donor-derived peripheral blood cells **(B)** and lineage contribution by donor-derived cells **(C)** after competitive transplantation of WT/Sema4AKO myHSC (CD45.2) into WT recipients (CD45.1) (n=5 animals per group). **D-F.** Experimental schema **(D)**, overall percentage of donor-derived peripheral blood cells **(E)** and and lineage contribution by donor-derived cells **(F)** after competitive transplantation of WT/Sema4AKO lyHSC (CD45.2) into WT recipients (CD45.1) (n=5 animals per group). **G-H.** Overall percentage of donor-derived cells in the bone marrow after competitive transplantation of WT/Sema4AKO myHSC (CD45.2) **(G)** and WT/Sema4AKO lyHSC (CD45.2) **(H)** into WT recipients (CD45.1), quantification and representative plot for the overall bone marrow chimerism are shown (n=5 animals per group). **I, J.** UMAP representation of 162 myHSC **(I)** and 165 lyHSC **(J)** from aged WT and Sema4AKO mice (n=2 mice per genotype). **K, L.** Distribution of diffusion pseudotime values of myHSC **(K)** and lyHSC **(L)** from aged WT and Sema4AKO mice. **M,N** Enrichment of up- and down-regulated genes in the aged Sema4AKO myHSC signature for the upregulated **(M)** and downregulated **(N)** genes in the HSC “aging signature”^52^ as assessed by GSEA. P values are shown. For the transplant experiments, *P<0.05, **P<0.05. Statistical significance was assessed by two-tailed *t*-test, except for the diffusion pseudotime analysis (P) where Wilcoxon rank sum test was used. Mean +/- SEM are shown.

Collectively, the transplant data are consistent with selective and profound loss of regenerative capacity by aged Sema4AKO myHSC. In addition, we note that a homing defect - a prominent feature of physiological aging^36^ which is accelerated Sema4AKO myHSC - may have also contributed to their engraftment failure.

Aiming to understand the molecular underpinnings of these observations, we performed single cell RNA-Seq analysis of myHSC and lyHSC from aged WT and Sema4AKO mice. Because we expected a high degree of transcriptional similarity between the two HSC subsets, we performed deeper, full-length scRNA sequencing using the SMART-Seq2 platform ^37^.

Transcriptome-wide comparison revealed that consistent with the myHSC-selective effect of Sema4A, as revealed by phenotypic and functional analyses, aged myHSC from WT and Sema4AKO mice formed distinct, minimally overlapping clusters, while the transcriptomes of aged WT and Sema4AKO lyHSC were virtually indistinguishable (Fig 2I, J, Extended Data Fig 2D). At the level of individual genes, aged Sema4AKO myHSC displayed higher expression of transcripts encoding for AP-1 family transcription factors (*Jun*, *JunD, Fos, FosB)*, which are known to be associated with inflammatory response and myeloid/megakaryocytic differentiation. ^38–43^ In contrast, several genes known to restrict HSC activation and inhibit differentiation (*Relb, Stat1, CD74, Irgm1*) ^44–47^ were downregulated (Extended Data Fig 2E). Overall, these data suggested that *in situ*-expanded aged Sema4aKO myHSC are hypersensitive to inflammatory stress and primed for myeloid differentiation, which may explain their dramatically reduced regenerative capacity.

To further explore these findings, we performed diffusion pseudotime (DPT) analysis ^48^, which can quantify the differentiation state of each cell going from naïve (corresponding to HSC) to less primitive (multipotent progenitors, MPP) state. For this analysis, we generated 10x Genomics single cell RNA-Seq profiles of lin^-^c-Kit^+^ HSPC from aged WT animals. In this dataset, we used previously described markers to map the clusters corresponding to HSC (*Ly6a, Procr, Hlf*) and MPP (*Cd34, Cebpa, Ctsg*) ^49^ and utilized the transcriptomes of cells within these clusters to estimate a differentiation trajectory, in which higher DPT values correspond to more mature cells (see Methods for details). Analysis of known self-renewing/differentiation marker genes ^50^ revealed downregulation of vWF, *Mpl*, *Fdg5*, C*tnnal1*, *Procr*, and upregulation of *Ctsg* and *Cbpa* as cells progressed from HSC to MPP, thus validating our analysis pipeline (Extended Data Fig 2F). Next, we used these differentiation trajectories to test the difference in the distribution of DPT values for aged WT/Sema4AKO myHSC and lyHSC, which we estimated with the Wilcoxon rank-sum test. In agreement with the published data^12^, we observed that the DPT values for WT aged myHSC were significantly lower than for WT aged lyHSC, suggesting that myHSC are positioned at the top of the HSC hierarchy (Extended Data Fig 2G). Applying DPT analysis to the transcriptomes of WT and Sema4AKO aged myHSC revealed the DPT values for the latter were significantly higher and closer to that of WT lyHSC, suggesting that the absence of Sema4A leads to myHSC shift to a more differentiated cellular state (Fig 2K). In contrast, no difference in DPT values were observed between WT and Sema4AKO aged lyHSC (Fig 2L).

Given that the pro-inflammatory signature and accelerated differentiation – the hallmarks of HSC aging^51^- are more prominently displayed in the transcriptome of aged Sema4AKO myHSC, we wondered whether these cells would be molecularly “more aged” overall. We therefore projected a published “aging signature” ^52^ onto the “pseudo-bulk” single cell RNA-Seq signature of myHSC from aged WT/Sema4AKO mice. As shown in Fig 2 M, N, both up- and down- regulated transcripts in the “aging signature” displayed significant enrichment for their respective counterparts in the aged Sema4AKO myHSC signature (P<0.01, FDR <0.01 in both directions), consistent with premature aging of Sema4AKO myHSC.

Taken together, these data provide functional and molecular evidence that during aging, Sema4A is required for myHSC resilience to inflammatory stress, prevention of differentiation and maintenance of stemness; however, Sema4A is largely dispensable for lyHSC in this physiological context.

### The absence of Sema4A in young animals leads to selective myHSC hyperactivation at the steady state

Having established that Sema4A functions as a critical hematopoietic regulator during aging, we wondered if the effect of Sema4A absence on hematopoiesis is already apparent in young animals. Interestingly, we discovered that neutrophilia and thrombocytosis in the peripheral blood, which had been present in aged Sema4AKO mice, were already detectable in young Sema4AKO mice, albeit considerably less prominent (Fig 3A). Similarly, in young Sema4AKO bone marrow, we observed a modest increase in the number myeloid/platelet-biased multipotent progenitor MPP2 and mature myeloid cells. However, selective myHSC expansion – the key feature of aged Sema4AKO phenotype – was absent in young Sema4AKO mice (Fig 3B-D).

**Fig 3.**
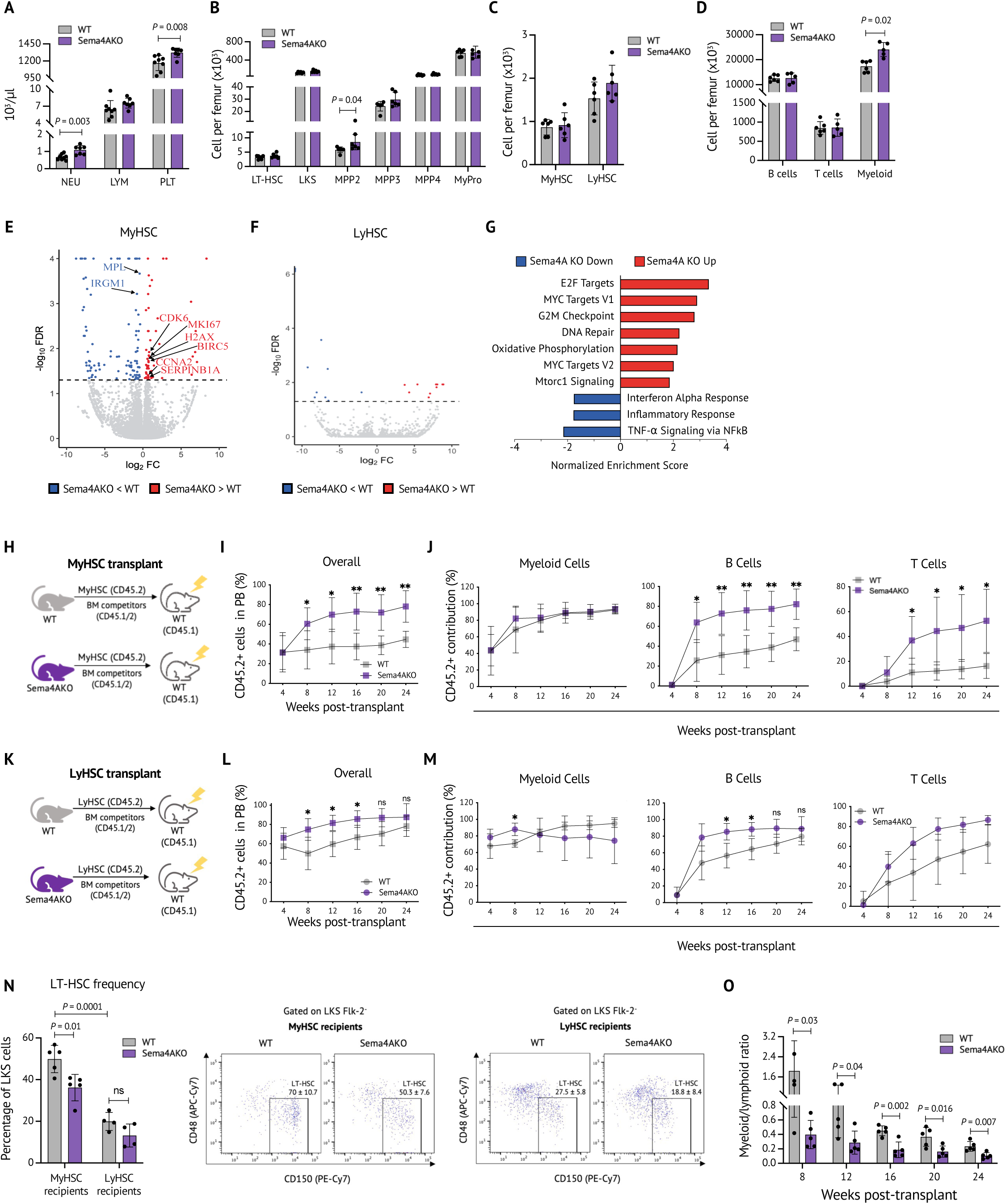
The absence of Sema4A in young animals leads to selective myHSC hyperactivation at the steady state. **A.** Baseline peripheral blood counts of WT and Sema4AKO mice (n=7 mice per group). **B-D.** Absolute number of primitive hematopoietic cells **(B)**, myHSC and lyHSC **(C)** and mature myeloid cells **(D)** in WT/Sema4AKO mice (n=6 mice per group). **E, F.** Volcano plots showing DEGs in myHSC **(E)** and lyHSC **(F)** from WT and Sema4KO mice. The *x* and *y* axes indicate the expression fold change (FC) (log_2_) and the false discovery rate (FDR) (−log_10_) for each gene versus controls, respectively. Legends highlight upregulated (red) or downregulated (blue) transcripts, as well as genes not passing cutoff criteria for FC (black) and FDR (gray). Selected representative genes are shown (n=4 biological replicates per genotype). **G.** Gene set enrichment analysis (GSEA) showing top overrepresented canonical pathways which are upregulated (red) or downregulated (blue) in Sema4AKO myHSC as compared to WT myHSC. The pathways with FDR <0.05 are shown (n=4 biological replicates per genotype). **H-J.** Experimental schema **(H)**, overall percentage of donor-derived peripheral blood cells **(I)** and lineage contribution by donor-derived cells **(J)** in WT mice (CD45.1) which were competitively transplanted with WT/Sema4AKO myHSC. (CD45.2) (n=3-5 mice per group). **K-M.** Experimental schema **(K)**, overall percentage of donor-derived peripheral blood cells **(L)** and lineage contribution by donor-derived cells **(M)** in WT mice (CD45.1) which were competitively transplanted with WT/Sema4AKO lyHSC (CD45.2). (n=5 mice per group). **N.** Frequency of LT-HSC (as percentage of Lin^-^Kit^+^Sca1^+^ cells) in the bone marrow of WT mice (CD45.1) after competitive transplantation of WT/Sema4AKO myHSC (myHSC recipients, CD45.2) and WT/Sema4AKO lyHSC (lyHSC recipients, CD45.2), quantification and representative plots are shown (n=4-5 mice). **O.** Myeloid/lymphoid ratio of peripheral blood donor-derived cells in WT mice which were competitively transplanted with WT/Sema4AKO myHSC. P values are shown. For the transplant experiments, *P<0.05, **P<0.05. Statistical significance was assessed by two-tailed *t*-test, mean +/- SEM are shown.

Nevertheless, RNA-Seq analysis of sorted CD150^high^ myHSC and CD150^low^ lyHSC showed that the absence of Sema4A induced evident transcriptional changes that were largely restricted to myHSC. In particular, we found that 154 genes were differentially expressed between WT/Sema4A myHSC. In contrast, only 20 genes were differentially expressed between WT/Sema4A lyHSC, none of which have been previously implicated in regulation of HSC function (Fig 3E, F, Extended Data Table 1). Examining Sema4AKO myHSC signature in more detail, we observed that expression of several genes which are known to promote HSC proliferation and loss of stemness (*Mki67*, *H2AX, CCNA2*, *Cdk6*) ^50, 53–56^ was increased in Sema4AKO myHSC while expression of *Mpl* and *Irgm1*, which are involved in maintenance of stem cell state, was reduced. Gene set enrichment analysis revealed upregulation of several pathways involved in cellular proliferation (“E2F targets”, “MYC targets”) and metabolic activation (“Oxidative phosphorylation,” “mTORC1 signaling”) (Fig 3G). Paradoxically, inflammatory pathways were downregulated, consistent with a tolerizing effect of excessive inflammatory stimulation. ^57^

Collectively, the results of phenotypic and transcriptional analysis indicate that the absence of Sema4A leads to enhanced proliferative and metabolic activation of myHSC (but not lyHSC) at the steady state, likely due to increased myHSC responsiveness to tonic inflammatory signals from gut microbiota, which are consistently present in conventionally housed animals.^58^ In keeping with these data, cell cycle analysis and BrdU incorporation studies revealed that Sema4AKO myHSC proliferation was increased, whereas Sema4A lyHSC proliferation was less uniformly affected and not statistically different between the groups (Extended Data Fig 3A-D).

### The absence of Sema4A in young animals results in loss of myHSC myeloid bias

Having established that the absence of Sema4A causes distinct phenotypic and molecular abnormalities in myHSC, we next tested their functional impact in a competitive transplantation assay. We used the same experimental design as for the aged WT/Sema4A cohort, except that a smaller number (200 cells per animal) of myHSC/lyHSC were transplanted. As an initial validation step, we asked whether young WT myHSC generated a myeloid-biased graft, as would be evidenced by a higher peripheral blood myeloid/lymphoid ratio of myHSC-derived progeny as compared to that of lyHSC-generated progeny. ^10^ This was indeed the case, thus validating our myHSC/lyHSC isolation strategy in young animals (Extended Data Fig 3E).

Assessing post-transplant behavior of young Sema4AKO myHSC, we found that, consistent with their metabolic and transcriptional hyperactivation in donor animals (Fig 3G), these cells displayed a markedly higher level of multi-lineage post-transplant chimerism compared to young WT myHSC, with the difference progressively increasing over time (Fig 3H-J). In contrast, although transplanted young Sema4AKO lyHSC produced slightly higher chimerism at earlier time points during post-transplant reconstitution, no difference to controls was observed by 24 weeks (Fig 3K-M). Bone marrow analysis at week 24 showed that in the recipients of Sema4AKO myHSC, the frequency of donor-derived long-term HSC (LT-HSC) was lower compared to the recipients of WT myHSC; however, the number of donor-derived LT-HSC in the lyHSC-transplanted cohort was not significantly different (Fig 3N). These data suggest that the absence of Sema4A leads to excessive myHSC differentiation, resulting in higher post-transplant output, but has only a minor effect on lyHSC behavior.

Previous studies have reported that – as we also observed – WT myHSC normally produce a lower level of donor chimerism compared to lyHSC, which is associated with their predominantly myeloid differentiation and suppression of the lymphoid potential. ^59, 60^ We therefore wondered if the aberrantly high level of donor chimerism generated by Sema4AKO myHSC, as we found, would be linked to the loss of the myeloid potential and enhanced differentiation towards the lymphoid lineage. To address this possibility, we calculated myeloid/lymphoid ratio of the peripheral blood progeny for each time point of the young WT/Sema4AKO myHSC transplant, which we expected to diminish if a relative proportion of Sema4AKO myHSC-derived lymphoid cells was to increase. Indeed, we observed that the myeloid/lymphoid ratio of the Sema4AKO myHSC-derived peripheral blood progeny was significantly lower than that of WT myHSC-derived progeny at several time points post-transplant. Notably, by post-transplant week 24, the myeloid/lymphoid ratio in the Sema4AKO myHSC cohort became almost identical to that of the WT lyHSC cohort (∼0.1) (Fig 3O, compare to Extended Data Fig 3E), indicating that Sema4AKO myHSC were no longer “myeloid-biased.”

To further explore this finding, we examined the primitive hematopoietic compartment in the post-transplant marrow of these animals. As expected, we found that in the WT myHSC cohort, the frequency of lymphoid-biased Flk2^med/high^ multipotent progenitors MPP4 ^61, 62^ and their level of Flk2 expression were lower compared to WT lyHSC cohort, consistent with the predominantly myeloid potential of WT myHSC (Extended Data Fig 3F, G). In contrast, in the Sema4AKO myHSC cohort, both MPP4 frequency and Flk2 expression level were increased and comparable to those observed in WT lyHSC cohort, suggesting that post-transplant Sema4AKO myHSC differentiation program has shifted towards the lymphoid lineage. Taken together, these observations support the result of peripheral blood post-transplant analysis and suggest that in the absence of Sema4A, immunophenotypic myHSC acquire increased lymphoid differentiation potential and lose myeloid bias – the key feature of their functional identity.

### Sema4AKO myHSC are hypersensitive to acute inflammatory stress

The experiments presented so far have examined the role of Sema4A under physiological conditions, i.e., steady-state and aging. However, given the key role of myHSC in host defense against viral and bacterial pathogens, we sought to test how the absence of Sema4A influences myHSC behavior in infectious disease-relevant models, such as response to acute and chronic innate immune agonists. Given that myHSC express higher levels of TLR4^20^ and preferentially respond to lipopolysaccharide (LPS) ^63^, we used LPS treatment as an myHSC-directed challenge.

We first employed an acute LPS treatment model, in which the animals received a single LPS dose (4 mg/kg) (Fig 4A), and performed a time course analysis to define the peak of myHSC regenerative response. These studies revealed that, consistent with the published data^64^, the greatest myHSC expansion occurred at 72 hours post-LPS injection, with myHSC comprising ∼80% of the total HSC pool, as compared to ∼20% at baseline (Extended Data Fig 4A,B). In addition, we observed an increase in the number of myeloid-biased multipotent progenitors MPP2 and MPP3 (Extended Data Fig 4C). Hence, the 72-hour time point was chosen for the subsequent experiments.

**Fig 4.**
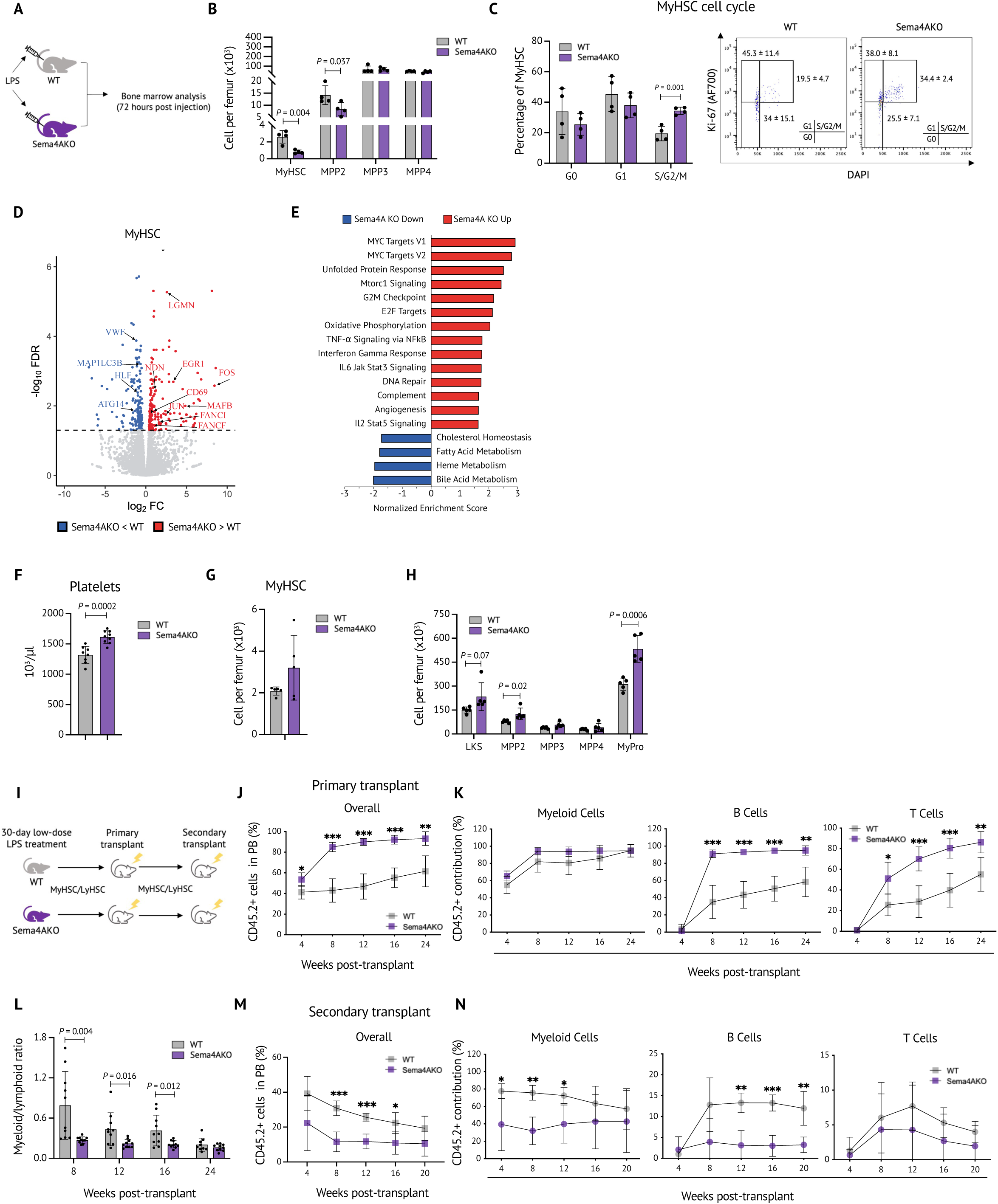
Sema4AKO myHSC are hypersensitive to acute innate immune activation. **A-C.** Experimental schema **(A),** absolute number of myHSC and progenitor subsets **(B)** and myHSC cell cycle analysis **(C)** in WT/Sema4AKO mice 72 hours post-LPS injection. **D.** Volcano plot representation of the RNA-Seq data from WT/Sema4AKO myHSC 72 hours post-LPS injection (n=4). **E**. Gene set enrichment analysis of myHSC from WT/Sema4AKO mice on day 3 post-LPS injection (n=4). **F-H.** Peripheral blood platelet count **(F)**, absolute number of myHSC **(G)** and primitive hematopoietic cells in WT/Sema4AKO mice **(H)** 30 days after treatment with low-dose LPS (n=5 mice per group). **I-K.** Experimental schema **(I)**, overall percentage of donor-derived peripheral blood cells **(J)** and lineage contribution by donor-derived cells **(K)** in primary recipient WT mice (CD45.1) which were competitively transplanted with myHSC from low-dose LPS-treated WT/Sema4A (CD45.2) mice (n=5 mice per group). **L.** Myeloid/lymphoid ratio of donor derived cells in WT mice (CD45.1) which were competitively transplanted with myHSC from low-dose LPS-treated WT/Sema4A (CD45.2) mice (n=5 mice per group). **M,N.** Overall percentage of donor-derived peripheral blood cells **(M)** and lineage contribution by donor-derived cells **(N)** in secondary recipient WT mice (CD45.1) which were competitively transplanted with myHSC from primary recipient mice (CD45.2) as shown in **(I)** (n=5 mice per group). P values are shown. For the transplant experiments, *P<0.05, **P<0.05. Statistical significance was assessed by two-tailed *t*-test. Mean +/- SEM are shown.

Given that acute LPS-induced inflammation causes changes in the HSC cell surface marker profile ^64^, we next tested if immunophenotypic definition of myHSC as CD150^high^ is still valid in this experimental setting. Because actively cycling HSC engraft poorly ^65^, addressing this question in functional studies, such as transplantation assay, is not feasible. We therefore asked if gene expression profile of myHSC which were immunophenotypically defined as CD150^high^ in LPS-injected animals would be enriched for a myHSC transcriptional signature which was obtained agnostic of CD150 expression level under similar inflammatory conditions. To this end, we generated RNA-Seq profiles of WT CD150^high^ myHSC and CD150^low^ lyHSC 72 hours post-LPS treatment and compared them to define a transcriptional profile of the CD150^high^ fraction.

We then overlapped this profile with a CD150-independent, functionally validated myHSC signature which had been previously published ^19^ and found a significant enrichment (p-value 2.9e-20 by hypergeometric test) (Extended Data Figure 4D). Thus, the CD150^high^ HSC fraction in the acute LPS-induced inflammation model is enriched for transcriptionally defined myHSC.

Next, we proceeded to investigate how Sema4A impacts myHSC response to acute LPS challenge. Immunophenotypic analysis of LPS-treated WT/Sema4AKO mice at 72 hours revealed that the absence of Sema4A markedly impaired regeneration of myHSC and multipotent progenitors, as the frequency and the absolute number of myHSC and MPP2 were significantly lower in Sema4AKO mice compared to WT controls (Fig 4B, Extended Data Fig 4E). Importantly, cell cycle analysis revealed that in Sema4AKO mice, the frequency of myHSC which were actively proliferating was significantly higher than in WT mice (Fig 4C). Given that increased HSC proliferative activity during acute inflammatory stress is known to cause a shift from self-renewing to asymmetric/differentiating divisions,^66^ our data suggest that in the absence of Sema4A, increased myHSC cycling resulted in depletion of myHSC through excessive differentiation.

To further test this hypothesis, we performed RNA-Seq of myHSC from WT and Sema4AKO mice 72 hours post-LPS injection. Our analysis showed that similar to the changes observed in aged Sema4AKO myHSC, several key molecules which “sense” inflammation (*EGR1* and AP-1 family members *Jun*, *Fos*, *MafB*) and mark myeloid differentiation (*CD69* and *LGMN)* were upregulated in “inflamed” Sema4AKO myHSC (Fig 4D). ^38–41^ Notably, we also observed a higher expression of *Ndn*, a known *p53* target gene and Fanconi anemia genes *FancI* and *FancF,* consistent with inflammation-induced activation of DNA repair pathways. ^67^ Interestingly, *HLF* and several genes in the autophagy pathway (A*tg14* and *MAP1LC3B),* which are known to promote HSC-self renewal, ^68, 69^ were downregulated, in keeping with a shift towards differentiation. Accordingly, gene set enrichment analysis (GSEA) revealed that compared to WT myHSC, Sema4AKO myHSC displayed significant upregulation of pathways involved in inflammatory response, cell cycle and DNA repair (Fig 4E). Collectively, our data demonstrates that the absence of Sema4A during acute inflammation results in increased myHSC proliferative stress and excessive activation of the myeloid differentiation program, which likely account for failed myHSC regeneration.

### Lack of Sema4A results in loss of myHSC self-renewal following chronic LPS exposure

Given that the direct impact of Sema4A on myHSC function in acute LPS-induced inflammation model is difficult to assess due to defective engraftment of actively proliferating cells, ^65^ we turned to chronic LPS-induced inflammation model. To this end, we implanted WT and Sema4AKO animals with intra-abdominal LPS-releasing pumps ^70^ and performed blood and bone marrow analysis at the end of the 30-day continuous infusion period.

Overall, we found that phenotypic and functional abnormalities in chronic LPS-treated Sema4AKO mice were greater than those present in these animals at the steady state. In particular, peripheral blood thrombocytosis was more pronounced, and changes in the HSPC compartment were more extensive, including expansion of MPP2 and kit+ myeloid-biased progenitors (Fig 4F-H). Similarly, upon transplantation, chronic LPS treatment caused a more prominent increase in Sema4AKO myHSC-derived multilineage chimerism, which rapidly (by week 8) reached the ∼90% mark (Fig 4I-K). Consistent with our prior observations in the straight Sema4AKO model, Sema4AKO myHSC from LPS-treated mice had compromised myeloid reconstitution potential, as evidenced by a lower myeloid/lymphoid ratio (Fig 4L). To assess the effect of chronic LPS treatment on Sema4AKO myHSC self-renewal, we performed secondary transplantation experiments. These revealed that in contrast to exaggerated output in primary transplant recipients, Sema4AKO myHSC generated much lower level of post-transplant chimerism in secondary recipient mice, indicating a severe self-renewal defect (Fig 4M, N).

Taken together, our results reveal that the absence of Sema4A markedly compromises myHSC resilience to chronic inflammatory stress leading to functional exhaustion, thus closely resembling the aged Sema4AKO myHSC phenotype.

### PlxnD1 is a functional receptor for Sema4A on myHSC

Since Sema4A is an extrinsic factor, we next sought to identify its cell surface receptor on myHSC. Sema4A is known to signal via several cell surface receptors across different cell types^30^. Analysis of published LT-HSC gene expression data ^71^ demonstrated that Plexin B2 (PlxnB2), Plexin D1 (PlxnD1) and Nrp1, a co-receptor for PlxnD1, displayed the most robust expression (Fig 5A). Although PlxnB2 was expressed at a higher level than PlxnD1, it was thought to be an unlikely candidate receptor for Sema4A for the following reasons. First, several features of the hematopoietic phenotype of animals deficient in PlxnB2 or another PlxnB2 ligand, Angiogenin, were the opposite of the Sema4AKO phenotype. ^29, 72^ For example, PlxnB2/AngKO mice displayed lower HSC post-transplant reconstitution and increased lymphoid bias with age, whereas HSC post-transplant reconstitution was increased in Sema4AKO mice, and their hematopoiesis was heavily myeloid-biased. Second, the level of PlxnB2 expression did not differ between MyHSC and lyHSC and the percentage of PlxnB2^+^ myHSC was slightly lower than PlxnB2^+^ lyHSC, which would be difficult to reconcile with the myHSC-selective effect of Sema4A (Extended Data Fig 5A, B). In contrast, both PlxnD1 and Nrp1 showed a higher level of expression in myHSC compared to lyHSC at the steady state, and their expression increased further during acute LPS-induced inflammation – a physiological context where Sema4A plays an important myHSC-regulatory role (Fig 5B, C, Extended Data Fig 5C, D). Notably, analysis of published data from patients with sepsis showed that the frequency of circulating PlxnD1+ CD34^+^ HSPC and myeloid progenitors (but not lymphoid progenitors) was markedly increased (Extended Data Fig 5E). ^73^. This indicates that in both mice and humans, PlxnD1 is upregulated during acute inflammatory response and thus may be functionally relevant Hence, we prioritized PlxnD1 for further validation.

**Fig 5.**
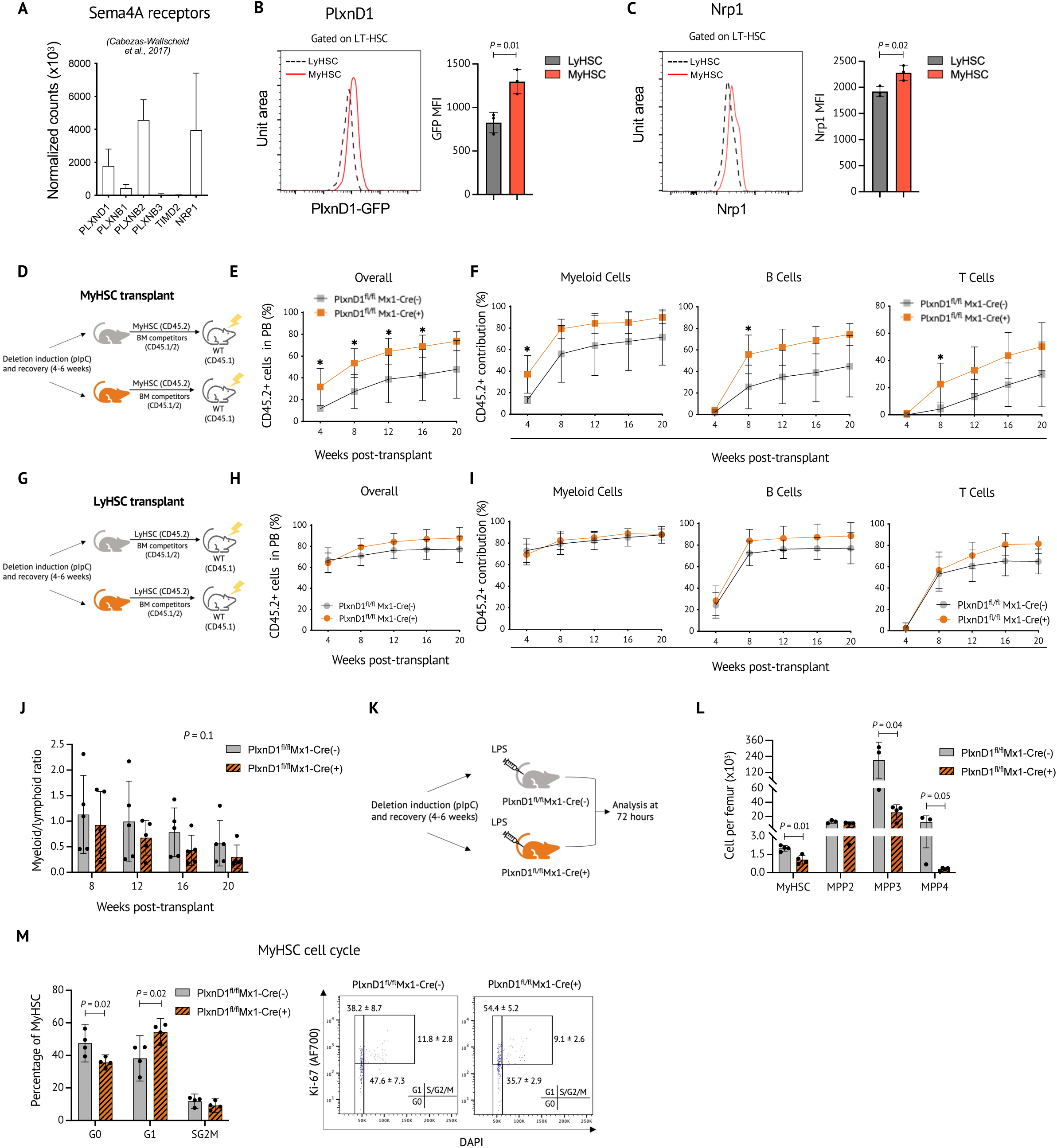
PlxnD1 is a functional receptor for Sema4A on myHSC. **A.** Expression of mRNA encoding for known Sema4A receptors in HSC (taken from Cabezas-Wallscheid, *Cell,* 2017)^67^. **B, C.** Expression of PlxnD1-GFP reporter **(B)** and Nrp1 protein **(C)** in myHSC and lyHSC in WT mice, as assessed by flow cytometry (n=3 biological replicates). **D-F.** Experimental schema **(D)**, overall percentage of donor-derived peripheral blood cells **(E)** and lineage contribution by donor-derived cells **(F)** in WT mice (CD45.1) transplanted with myHSC from untreated PlxnD1fl/fl Mx1Cre(+) and PlxnD1fl/fl Cre(-) mice (CD45.2) (n=5 mice per group). **G-I.** Experimental schema **(G)**, overall percentage of donor-derived peripheral blood cells **(H)** and lineage contribution by donor-derived cells **(I)** in WT mice (CD45.1) transplanted with lyHSC from untreated PlxnD1fl/fl Mx1Cre(+) and PlxnD1fl/fl Cre(-) mice (CD45.2) (n=5 mice per group). **(J)** Myeloid/lymphoid ratio of donor-derived cells in WT mice (CD45.1) transplanted with myHSC from PlxnD1fl/fl Mx1Cre(+) and PlxnD1fl/fl Cre(-) mice (CD45.2) (n=5 mice per group). Overall statistical significance assessed by ANOVA. **K-M**. Experimental schema **(K),** absolute number of myHSC and progenitor subsets **(L)** and myHSC cell cycle analysis **(M)** in PlxnD1fl/fl Mx1Cre(+) and PlxnD1fl/fl Cre(-) mice 72 hours post-LPS injection. P values are shown. For the transplant experiments, *P<0.05, **P<0.05. Statistical significance was assessed by two-tailed *t*-test unless otherwise stated. Mean +/- SEM are shown.

Global deletion of PlxnD1 in mice is embryonic lethal due to structural cardiac and vascular defects, thus precluding functional analysis of adult HSC from these animals. ^74^ We therefore conditionally deleted PlxnD1 by crossing PlxnD1 “floxed” mice with the Mx1-Cre strain. ^75^ We confirmed a complete excision of PlxnD1 by PCR and Q-PCR analysis of sorted Lin^-^Kit^+^Sca^+^ cells after PolyIC induction (Extended Data Fig 5F, G). Analysis of peripheral blood and the bone marrow of PlxnD1^fl/fl^ Mx1-Cre(+) and PlxnD1^fl/fl^ Mx1-Cre(-) mice at the steady state showed no abnormalities (Extended Data Fig 5H-L). However, competitive transplantation experiments revealed that similar to the behavior of myHSC/lyHSC from Sema4AKO mice, myHSC from PlxnD1^fl/fl^ Mx1-Cre(+) donors generated a higher level of multi-lineage chimerism compared to PlxnD1^fl/fl^ Mx1-Cre(-) controls, whereas PlxnD1^fl/fl^ Mx1-Cre(+) and PlxnD1^fl/fl^ Mx1-Cre(-) lyHSC displayed equivalent engraftment levels (Fig 5D-I). Assessment of graft lineage composition revealed that especially at later time points, PlxnD1-deficient myHSC displayed a lower myeloid/lymphoid ratio (similar to Sema4AmyHSC) although the difference did not reach statistical significance (Fig 5J, p-value 0.1 by ANOVA). Post-transplant bone marrow analysis at week 20 demonstrated that PlxnD1 deletion in myHSC (unlike Sema4A deletion) had no measurable impact on the frequency of myHSC-derived HSC and MPP4 (Extended Data Fig 5M, N). However, the level of MPP4 Flk2 expression in the PlxnD1-deleted cohort was higher and consistent with a shift towards lymphoid differentiation, as seen in the Sema4AKO model (Extended Data Fig 5O). Importantly, PlxnD1 deletion was inconsequential for lyHSC differentiation (Extended Data 5M-O). Cumulatively, these results show that although the impact of PlxnD1 deletion on myHSC was somewhat less pronounced than that of Sema4A, it recapitulates the key aspect of the Sema4AKO phenotype – the myHSC-selective effect.

Given a newly discovered role of Sema4A in regulating myHSC sensitivity to acute LPS-induced inflammatory stress, we asked if PlnxD1 deletion also resembles that of Sema4A in this experimental model. These experiments demonstrated that 72 hours post-LPS injection, PlxnD1-deficient myHSC excessively proliferated, became depleted in number and generated fewer downstream multipotent progenitors, as seen in the Sema4AKO phenotype (Fig 5K-M, Extended Data Fig 5P). These results demonstrate that like the absence of Sema4A, the absence of PlxnD1 enhances myHSC sensitivity to acute inflammatory stress and compromises their regenerative function.

Taken together, our results provide several lines of experimental evidence for a newly defined function of PlxnD1 as a Sema4A receptor on myHSC. However, we cannot rule out that other cell surface molecules and in particular Nrp1, a co-receptor for PlxnD1, are involved in Sema4A signaling in myHSC.

### The effect of Sema4A is non cell-autonomous

In the next set of experiments, we asked whether Sema4A acts as a cell-autonomous, i.e., myHSC-derived, or a cell non-autonomous regulator. Examination of Sema4A expression in the bone marrow at baseline (as we all as during acute inflammation and aging) showed that in myHSC, Sema4A mRNA and protein levels were low/undetectable; in contrast, this molecule was abundantly expressed in granulocytes and monocytes (Extended Data Fig 6A,B), which served as predominant sources of Sema4A in the bone marrow (Fig 6A, Extended Data Fig 6C). Such expression pattern suggested that Sema4A likely acts as a non cell-autonomous, myHSC-extrinsic regulator.

**Fig 6.**
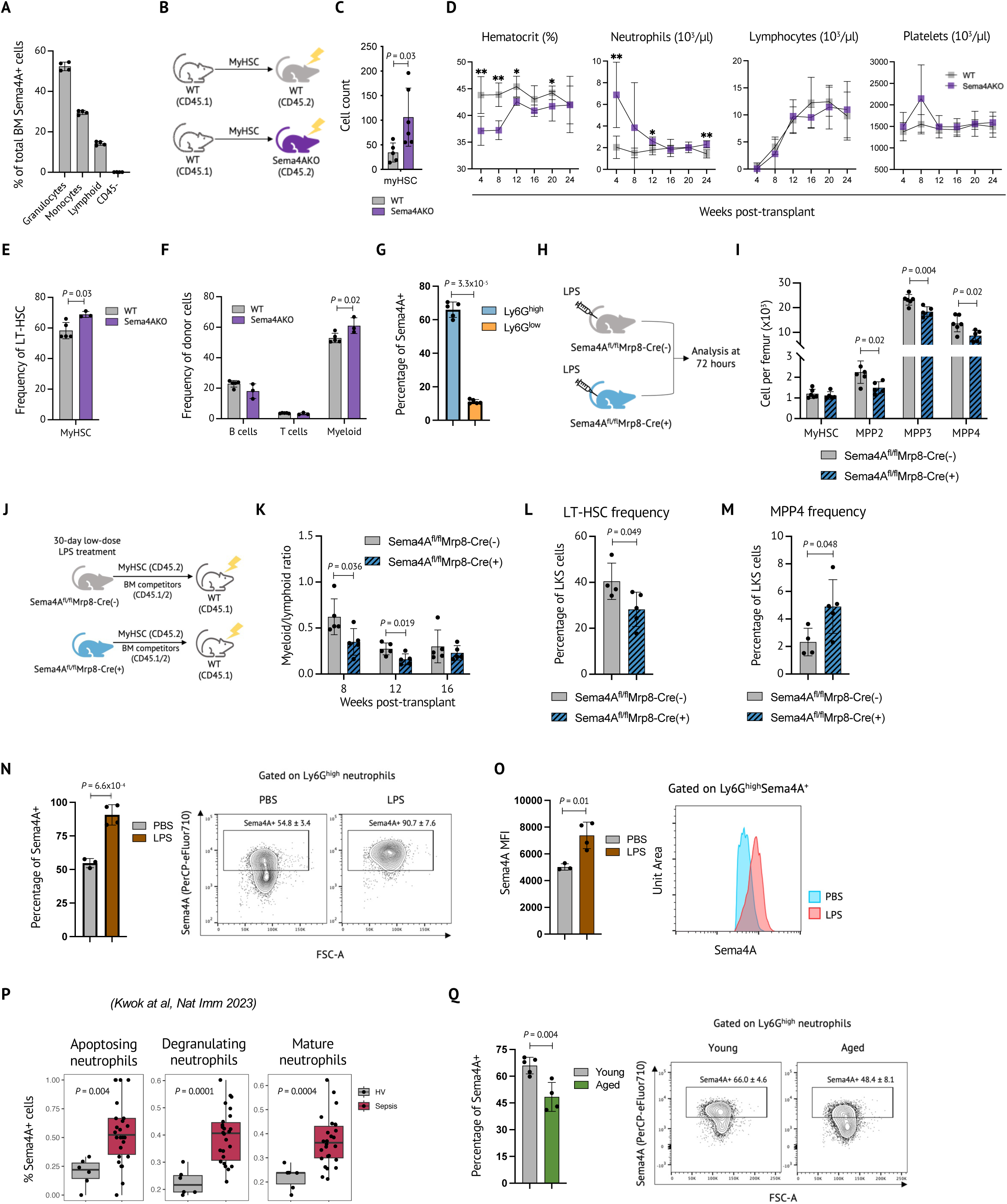
Neutrophils serve as a physiologically important source of Sema4A. **A.** Relative contribution of distinct cellular subsets to Sema4A production in the bone marrow at the steady state (n=3-4 mice per group)**. B-C.** Experimental schema for non-competitive transplantation of WT myHSC into lethally irradiated WT/Sema4AKO recipients **(B),** the number of transplanted myHSC progeny in calvarial bone marrow, as assessed by intra-vital microscopy at the 24-hour time point **(C)** (n= 4 mice per group)**. D-F.** Longitudinal blood counts **(D),** donor-derived LT-HSC frequency **(E)** and mature cell frequency **(F)** in the bone marrow of a separate WT/Sema4AKO cohort which was non-competitively transplanted with WT myHSC (n= 3-5 mice per group) **(G).** Sema4A expression in Ly6G^high^ vs Ly6G^low^ neutrophils (n=5 mice per group). **H,I.** Experimental schema **(H)** and absolute number of myHSC and primitive hematopoietic cells **(I)** in Sema4A^fl/fl^ Mrp8-Cre(+) and Sema4A^fl/fl^ Mrp8-Cre(-) 72 hours after LPS injection (n=4-7 mice per group). **J-M.** Experimental schema **(J)**, myeloid/lymphoid ratio of peripheral blood donor-derived cells **(K),** donor-derived LT-HSC frequency **(L)** and donor-derived MPP4 frequency **(M)** in WT mice (CD45.1) which were competitively transplanted with myHSC from low-dose LPS-treated Sema4A^fl/fl^ Mrp8-Cre(+) and Sema4A^fl/fl^ Mrp8-Cre(-) (CD45.2) mice (n=5 mice per group). **N, O.** Sema4A expression in bone marrow WT Ly6G^high^ neutrophils 24 hours post-LPS injection, as quantified by frequency of Sema4A^+^ cells **(N)** and Sema4A mean fluorescence intensity (MFI) of Sema4A+ Ly6G^high^ neutrophils **(O)** (n=4 mice per group) Bar graph and representative flow cytometry plots are shown. **P.** Frequency of Sema4A^+^ cells in peripheral blood neutrophil subsets from patients with sepsis and healthy volunteers (HV), as assessed by single cell RNA-Seq in Kwok et al.^69^ **Q.** Frequency of Sema4A^+^ Ly6G^high^ neutrophils in the bone marrow of young and aged WT mice (n=4-5 mice per group). P values are shown. For the transplant experiments, *P<0.05, **P<0.05. Statistical significance was assessed by two-tailed *t*-test. Mean +/- SEM are shown.

To investigate the effect of global microenvironmental deletion of Sema4A on myHSC, we first performed intra-vital microscopy of transplanted WT myHSC in lethally irradiated WT/Sema4AKO animals. We transferred an equal number of WT myHSC (which had been fluorescently labeled ex vivo) and imaged these cells in the calvarial bone marrow of live mice 24 hours later (Fig 6B). Notably, we observed that the number of transplanted cells in the Sema4AKO-deficient hosts was significantly higher, consistent with their faster proliferation (Fig 6C, Extended Data Fig 6D), i.e. similar to what we have observed in non-transplanted Sema4AKO animals (see Figure 4C, Extended Data Fig 3A). Interestingly, flow cytometry showed that this time point post-irradiation, neutrophils became the main source of Sema4A in the bone marrow, contributing to almost 90%, and that neutrophil expression of Sema4A on per-cell basis also increased (Extended Data Fig 6E).

Next, we examined long-term reconstitution kinetics of WT myHSC in WT/Sema4A-deficient bone marrow microenvironment using the same experimental setting. We found that Sema4AKO recipients developed anemia, neutrophilia in the peripheral blood, as well as myHSC expansion and myeloid bias in the bone marrow (Fig 6D-F), i.e. the features which were also present in Sema4AKO mice, particularly during aging (see Fig 1A-C, Fig 3 A, D).

Collectively, these transplantation experiments show that pan-cellular deletion of Sema4A in the host bone marrow microenvironment is sufficient to recapitulate the key aspects of the Sema4AKO phenotype.

### Neutrophils serve as a physiologically important source of Sema4A

Having established the Sema4A acts in a non cell-autonomous fashion, we next focused on granulocytes as a predominant cellular source of Sema4A. We observed that amongst the granulocytes, Sema4A^+^ expression was particularly high in the mature (Ly6G^high^) ^76^ neutrophil fraction (Fig 6G, Extended Data Fig 6F). To directly test an impact of neutrophil-derived Sema4A on myHSC, we crossed Sema4A “floxed” mice with Mrp8-Cre strain.^77^ We observed Cre-mediated excision in over 90% of neutrophils (Extended Data Fig 6G), consistent with the published data.^78^ At baseline, Sema4A^fl/fl^ Mrp8-Cre(+) mice had no phenotypic hematopoietic abnormalities or functional myHSC defects (Extended Data Fig 6H-J and data not shown). We therefore examined the phenotype of these mice under acute and chronic inflammatory conditions. Following acute LPS challenge, Sema4A^fl/fl^ Mrp8-Cre(+) mice displayed exaggerated response, as evidenced by reduced frequency and the absolute number of MPPs (but not myHSC), (Fig 6H,I, Extended Data Fig 6 K,L), mirroring major findings in straight Sema4AKO and PlxnD1KO strains. Moreover, following chronic LPS treatment, myHSC from Sema4A^fl/fl^ Mrp8-Cre(+) mice displayed similar post-transplant behavior to that of straight Sema4AKO myHSC, i.e. lymphoid-biased reconstitution in the peripheral blood, as well as defective regeneration of HSC but enhanced production of lymphoid-biased MPP4 in the bone marrow (Fig 6J,M, Extended Data Fig 6M,N).

Collectively, these findings demonstrate that neutrophils serve as physiologically important, non-redundant cellular source of Sema4A in the bone marrow niche. Given that neutrophils are a direct progeny of myHSC, our data suggest the existence of a feedback loop whereby neutrophil safeguard their own production by maintaining inflammation resilience and “lineage fidelity” of myHSC.

### Neutrophil expression of Sema4A is dynamically regulated during inflammation and aging

Finally, we asked whether changes in neutrophil Sema4A expression also occur during inflammation and aging, which would link our findings in the genetic knockout models to a potential role of Sema4A in normal physiological settings. Interestingly, we discovered that in mice, both the frequency of Sema4A^+^ neutrophils within the Ly6G^high^ fraction and their level of Sema4A expression markedly increased following acute treatment with LPS (Fig 6N,O).

Similarly, in humans with sepsis, the frequency of Sema4A^+^ neutrophils across several neutrophil subsets, including mature neutrophils - but not in lymphoid cells - was also markedly elevated (Fig 6P, Extended Data Fig 6O).

Similarly, in humans with sepsis, the frequency of Sema4A^+^ neutrophils across several neutrophil subsets, including mature neutrophils - but not in lymphoid cells - was also markedly elevated (Fig 6P, Extended Data Fig 6P). Given our findings that a total absence of Sema4A leads to premature hematopoietic aging-like phenotype, this observation raises a possibility that age-related decrease in neutrophil-derived Sema4A may contribute to the onset of physiological hematopoietic aging.

### Sema4A maintains the epigenetic identity of myHSC

The data from multiple models employed in our study reveal that Sema4AKO myHSC display inferior self-renewal and lymphoid-skewed post-transplant differentiation, as compared to myHSC from controls. Intriguingly, the same features characterize WT lyHSC when compared to WT myHSC^12, 79^. A recent study by Meng et al^80^ has shown that in case of WT lyHSC/WT myHSC, the functional differences between the two HSC subsets are closely linked to their distinct epigenetic signatures. We therefore hypothesized that a similar, epigenetically-driven mechanism may explain the functional differences between WT/Sema4AKO myHSC that we observed.

To test this hypothesis, we performed Assay for Transposase-Accessible Chromatin with high-throughput sequencing (ATAC-Seq) profiling of these cells from aged WT/Sema4AKO mice.

Assessment of the chromatin accessibility using principal component analysis (PCA) showed that PC1 distinguishes the genome-wide ATAC-Seq profiles of aged Sema4AKO myHSC from those of aged WT myHSC (Fig 7A). Consistent with this, we detected 242 ATAC-seq peaks, or open chromatin regions (OCRs) with an increase in signal and 144 OCRs with a decrease in signal in the aged myHSC ATAC-Seq profiles compared to those of aged WT myHSC (Fig 7B). Thus, Sema4AKO myHSC appear to be epigenetically distinct from WT myHSCs.

**Fig 7.**
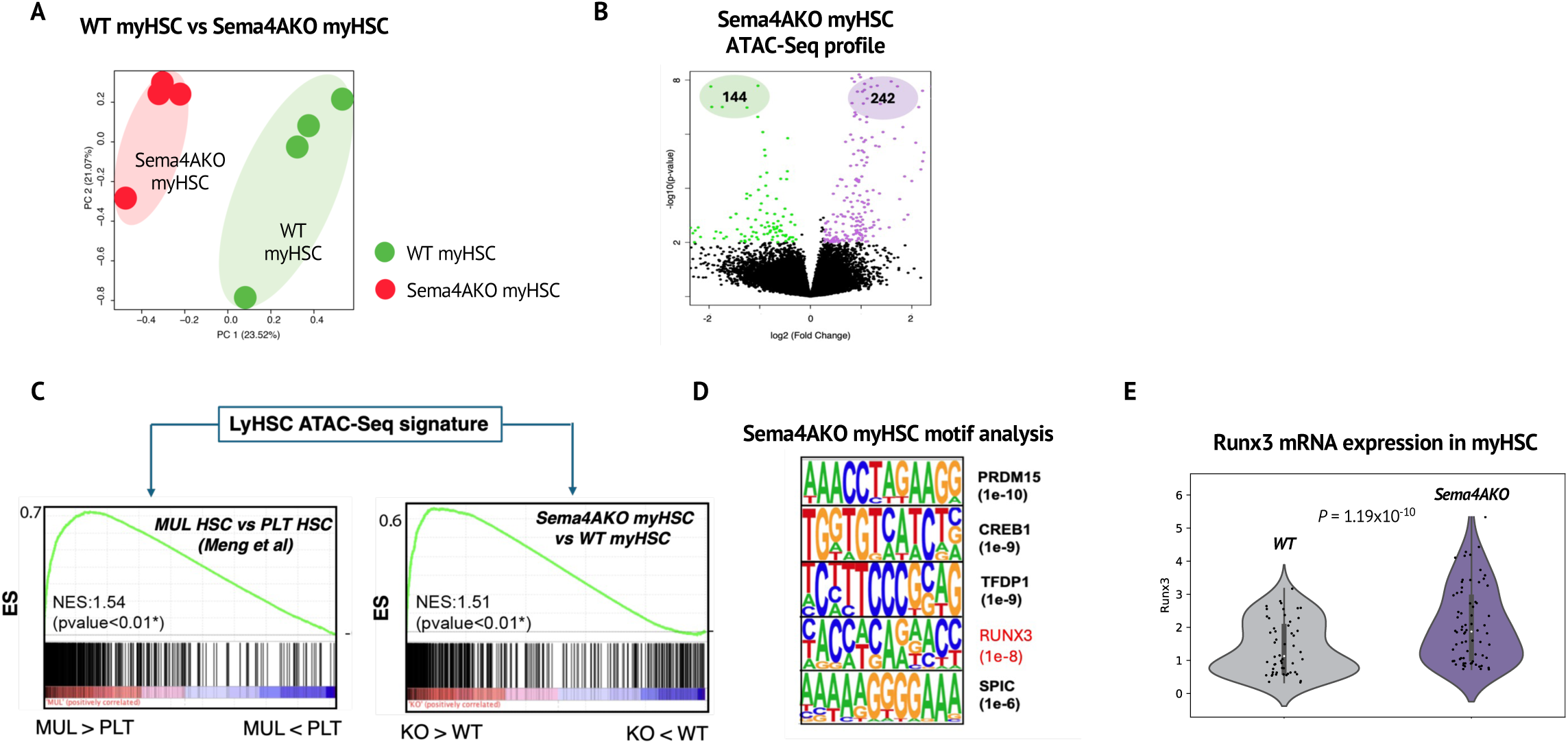
Sema4A maintains the epigenetic identity of myHSC. **A.** PCA analysis of ATAC-Seq signatures of myHSC from aged WT/Sema4AKO mice. **B.** Volcano plot of open chromatin regions that show increased accessibility (n=242) and decreased accessibility (n=144) in aged Sema4AKO myHSC. **C.** Gene Set Enrichment Analysis GSEA) of the lyHSC differentially accessible gene signature over the lyHSC gene signature from Meng et al (MUL vs PLT; left panel) and the Sema4A myHSC accessibility genes (right panel). Note that in the Meng et al dataset, MUL and PLT denotes lyHSC and myHSC comparable HSC subsets, respectively. **D.** Enriched denovo motifs predicted from myHSC/Sema4AKO enriched open chromatin. **E.** Gene expression level of Runx3 in aged WT/Sema4AKO myHSC (from RNA-seq data)

As mentioned above, Sema4AKO myHSC display an lyHSC-like in vivo behavior. To determine whether the changes we observed in Sema4AKO myHSC makes them epigenetically more similar to lyHSCs, we first generated a WT lyHSC ATAC-Seq signature. Specifically, we compiled a list of differentially regulated genes with upstream regulator elements (URE) within 1-50 kb from the transcription start site that showed increased accessibility in aged WT lyHSC (which we also profiled) as compared to aged WT myHSC (Extended Data Table 1). As expected, our lyHSC ATAC-Seq signature was strongly enriched in genes that were previously identified by Meng et. al. when WT lyHSC (referred to as MUL HSC) and WT myHSC (referred to as PLT HSC) were compared by ATAC-Seq profiling (Fig 7C; left panel). Furthermore, we observed a similar enrichment of this lyHSC ATAC-seq gene signature in genes that showed an increased accessibility of the URE in Sema4AKO myHSC as compared to WT myHSC (Fig 7C; right panel). These results indicate that functional distinctions between WT myHSC/WT lyHSC and WT myHSC/Sema4AKO myHSC with respect to self-renewal and lineage bias are likely driven by a shared epigenetic mechanism. However, the global difference between Sema4AKO myHSC and WT myHSC may be less pronounced than the difference between WT myHSC and WT lyHSC where we found a larger number of differentially regulated OCRs (2,654 and 2,225, respectively, data not shown).

Next, we performed motif enrichment analysis to identify DNA-binding transcription factors that may be alerted and contribute to changes in the epigenetic programming of the myHSC pool in response to the loss of Sema4AKO. Among our top hits in the accessible regions in the Sema4AKO myHSCs were motifs that matched the preferred binding sites of the transcription factors PRDM15, CREB1, TFDP1, RUNX3 and SPIC (Fig 7D). In addition, we find that Runx3 transcription is significantly upregulated in myHSC in the Sema4AKO condition based on our single-cell RNA-seq analysis (Fig 7E). Interestingly, the study of WT lyHSC/myHSC by Meng et. Al. found binding sequences for RUNX3 - a chromatin pioneer factor which acts as a master regulator of HSC lymphoid bias – were highly enriched in the accessible chromatin of WT lyHSC, and that *Runx3* mRNA level in lyHSC is higher. Thus, WT lyHSC and Sema4AKO myHSC share not only the global features of WT lyHSC epigenetic signature but also specific molecules which regulate their chromatin state.

Taken together, our data highlight the critical role of Sema4A in maintaining the epigenetic identity of myHSC. In the absence of Sema4A, the epigenetic distinctions between myHSC and lyHSC become blurred, and myHSC start to functionally resemble lyHSC, leading to the loss of diversity within the total HSC pool

## DISCUSSION

In this work, we define Sema4A as a novel extrinsic factor which selectively protects a specialized stem cell subset – myeloid-biased HSC – from inflammation-induced injury. Although prior studies have suggested that myHSC may rely on dedicated signals to preserve their function ^11, 20, 21^, those that are important in the context of inflammatory stress – to which myHSC are most vulnerable – have not, to our knowledge, been previously identified. We show that the protective role of Sema4A is functionally relevant across several animal models of inflammation, such as tonic inflammatory stimulation, LPS-induced emergency myelopoiesis and chronic low-dose LPS exposure but is most critical during aging when the lack of Sema4A leads to marked *in situ* amplification of phenotypic HSC but an almost complete loss of their competitive fitness upon transplantation.

Our gene expression data show that both “inflamed” and aged Sema4AKO myHSC signature is dominated by genes which are known to be upregulated in response to inflammatory stress, such as *EGR1*, *Jun* and *Fos*.^42^ This indicates that excessive expansion of aged Sema4AKO myHSC and their subsequent regenerative failure resulted from increased sensitivity to inflammatory cues, which are abundant during “inflammaging” ^51^. In light of recent findings that inflammatory signals drive hematopoietic aging ^17^, our results demonstrate that when resistance to these signals is compromised – as occurs in the absence of Sema4A – myHSC aging, as well as overall hematopoietic aging, are premature. Thus, we propose that by protecting myHSC from inflammatory stress and safeguarding their integrity throughout the animal’s lifespan, Sema4A maintains the overall functional diversity of the HSC pool and ensures hematopoietic longevity.

The striking functional deficit of aged Sema4AKO myHSC that we observed is likely a consequence of myHSC hyperactivation at young age, which is evident from our transcriptomic and functional studies. Interestingly, our experiments in young WT/Sema4AKO animals showed that in addition to regulating “generic” myHSC properties, such as self-renewal and multi-lineage reconstitution, Sema4A was also responsible for maintaining two key functional characteristics of myHSC, namely lower post-transplant output and predominantly myeloid-biased differentiation. ^60^ We showed that upon transplantation, Sema4AKO myHSC lose functional identity, since their level of peripheral donor chimerism and lineage output become comparable to that of WT lyHSC. In this regard, it is noteworthy that the “lyHSC drift” of young Sema4AKO myHSC correlated with transcriptional evidence for upregulation of oxidative phosphorylation – a metabolic hallmark of lyHSC, which rely on intact mitochondrial function for maintenance of the lymphoid differentiation potential. ^59^ Thus, we hypothesize that by preventing myHSC transition to oxidative phosphorylation, Sema4A may control a metabolic node which is required to maintain the functional myHSC phenotype. Notably, a similar “metabolic stabilizer” function for Sema4A has been previously described in another long-lived cell population – regulatory T cells – where Sema4A prevented overactivation of Akt-mTOR signaling, a key regulator oxidative phosphorylation and mitochondrial function.^31^ Interestingly, excessive mitochondrial metabolism via upregulation of CD38 was recently reported to impair HSC function during aging ^81^, suggesting that myHSC metabolic hyperactivation in young animals may have contributed to the aged Sema4AKO myHSC phenotype.

Unexpectedly, we discovered that the effect of Sema4A is non cell-autonomous and that neutrophils (in particular, their mature Ly6C^high^ fraction) serve as a major cellular source of Sema4A in the bone marrow. Our experiments using Mrp8-Cre Sema4A^fl/fl^ clearly demonstrated that neutrophil-derived Sema4A regulates myHSC, although the phenotype was not as strong as that of global Sema4A deletion, suggesting that other cell types also contribute. Neutrophil-derived Sema4A likely acts locally, since prior studies have shown that Gr1^+^ myeloid cells are evenly distributed throughout the marrow, positioned in close proximity (<50 μm) to all c-Kit*^+^*hematopoietic stem/progenitor cells and serve as a source of paracrine HSC-regulatory signals.^82, 83^

We found that Sema4A expression in Ly6C^high^ neutrophils becomes markedly upregulated following acute inflammatory insult, likely via an NF-kB-mediated mechanism since Sema4A promoter contains an NF-KB binding site. ^84^ Given our data that the lack of neutrophil-derived Sema4A impairs myHSC function, this observation suggests that an increase in neutrophil-derived Sema4A expression serves as a myHSC-protective mechanism. We propose that this mechanism operates via a negative feedback loop whereby neutrophils – a direct progeny of myHSC – secrete Sema4A to prevent inflammation-induced myHSC damage. However, it appears that during aging, this mechanism fails since Sema4A production by Ly6G^high^ neutrophils decreases due to significant loss of Sema4A+ expressing cells within this population. Given that the total absence of Sema4A resulted in premature aging-like phenotype and that neutrophils serve as a major source of Sema4A in the bone marrow, our data suggest that age-related decrease in neutrophil-derived Sema4A may contribute to physiological myHSC aging. This observation supports the emerging notion that dysregulated production of paracrine factors by the aging microenvironment – for example, decreased secretion of IGF1 in the bone marrow niche ^85^ or increased production of FGF1 in the skeletal muscle stem cell niche ^86^ – is sufficient to drive progressive decline of stem cell function. Hence, pharmacological restoration of niche-derived signals may offer an opportunity for anti-aging therapeutic intervention. ^87^

We identify PlxnD1 as a functional receptor for Sema4A on myHSC. While this molecule was previously known as a regulator of endothelial and mature immune cells, ^74^ ^88^ its role in hematopoiesis had not, to our knowledge, been previously defined. Our data demonstrates that PlxnD1 serves as an inhibitory receptor on myHSC and attenuates the intensity of inflammatory response upon binding Sema4A. Similar to Sema4A, PlxnD1 expression is upregulated during acute inflammation and increases in myHSC in mice and HSC in humans, thus further enhancing the strength of protective Sema4A signaling. Given that in some experimental settings, deletion of PlxnD1 resulted in less pronounced changes than deletion of Sema4A, it is likely that PlxnD1 is not the only molecule that transmits Sema4A signals. We hypothesize that PlxnD1 co-receptor Nrp1 ^89^ also contributes, since it shows higher expression on myHSC and is further upregulated in response to acute inflammation.

Finally, we note that while our study highlights the importance of HSC-directed “inflammation resistance” signals, such as Sema4A, in mouse inflammatory hematopoiesis, our work may have direct translational relevance to human health. Recent studies have shown that in humans, HSC inflammatory memory (which is epigenetically mediated) underlies hematopoietic and immune defects in aging, clonal hematopoiesis and inflammatory disorders^90^. Thus, reducing HSC inflammation responsiveness via dampening the acquisition of such a memory would be clinically beneficial. Given that Sema4A serves as an “inflammation resistance” signal for myHSC and operates at the chromatin level, as we have discovered, recombinant Sema4A protein or small molecule agonists of Sema4A/PlxnD1 may serve as future therapeutic agents to fulfil this role.

## LIMITATIONS OF THE STUDY

Our work established that Sema4A/PlxnD1 preferentially regulates myHSC, which were defined based on a higher CD150 expression, as per several prior reports. ^10, 60^ However, CD150 expression is a continuum and thus only provides a relatively imprecise myHSC definition.

Future experiments using unbiased strategies, such as *in vivo* clonal tracking, could help better resolve an HSC subset specifically regulated by Sema4A/PlxnD1 and identify the underlying molecular mechanism. We also acknowledge that our current set of observations is limited to a single innate immune agonist – LPS. Thus, we cannot definitively conclude if Sema4A/PlxnD1 are “generic” inhibitors of innate immune activation or whether their effect is restricted to TLR4-triggered inflammatory response.

## Supporting information

Reagent List

## ACKNOWLEDGEMENTS

The authors would like to acknowledge the Core Facilities of Fred Hutchinson Cancer Center, E Regiarto, M Baum and K Krum for technical help, and Dr A Wilkinson and Dr S Mckinney-Freeman for helpful discussions and critical reading of the manuscript.

## CONTRIBUTIONS

LS conceived the project, performed experiments and wrote the manuscript. DT, SG, SZ, NS, CB, DG, SR, SR, NC, AP, AG, FL, NKW, SJK, LZ, AT, MM designed and performed experiments. HPK, NC, BG, IB, BP, DJ, DJ, CN, EP, CM, AK, TW, SZJ, PK, DTS, AS and JAS performed data analysis and contributed to manuscript writing.

## FIGURE LEGENDS

**Extended Data Fig 1. (related to Fig 1).**
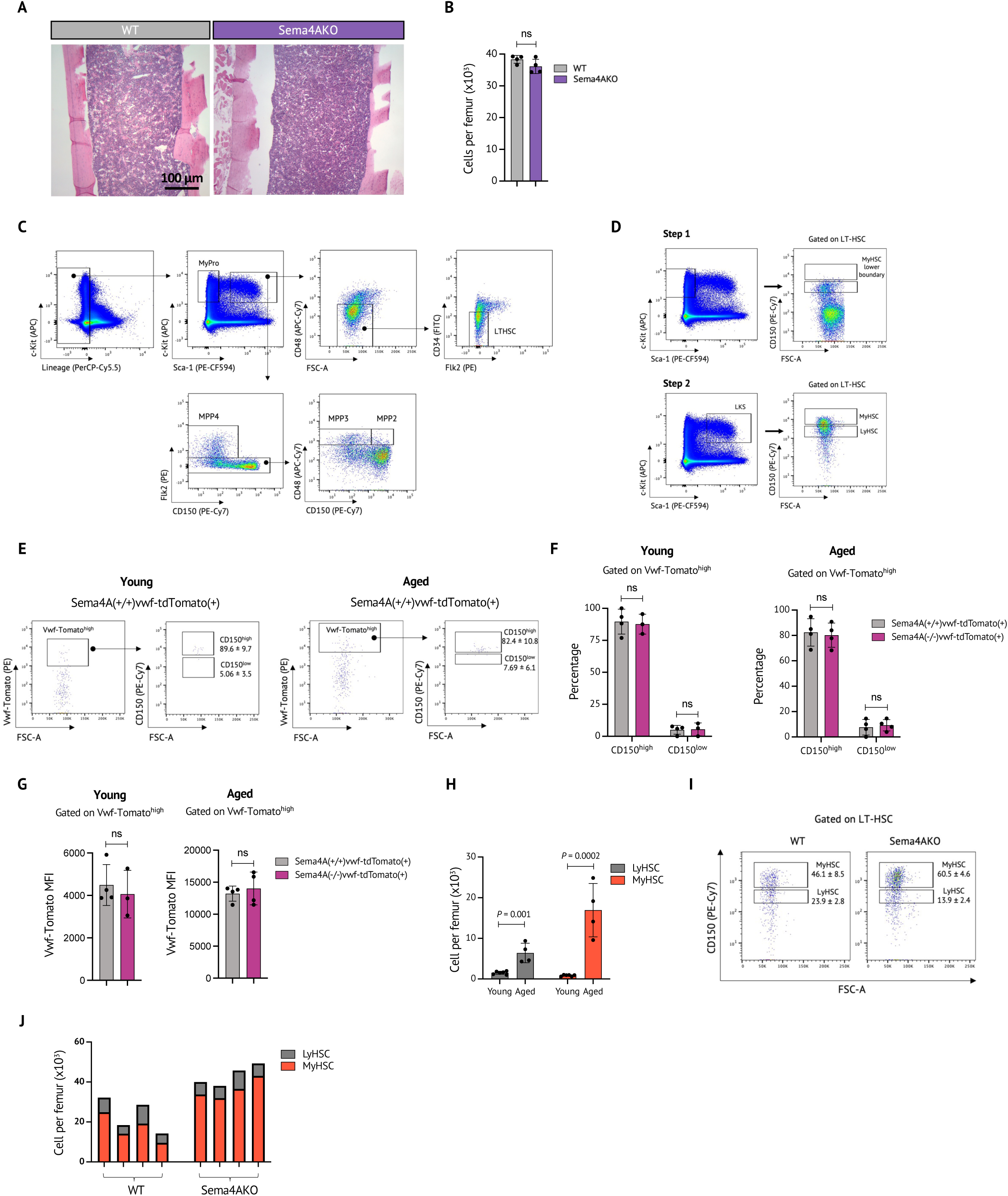
**A.** Representative images of H&E staining of femurs from aged WT/Sema4AKO mice. **B.** Bone marrow cellularity of aged WT/Sema4A mice (n=4-5 mice per group). **C.** Gating strategy for flow cytometric analysis of primitive bone marrow subsets (WT aged mice are shown). LT-HSC denotes long-term HSC. **D.** Gating strategy for flow cytometric identification of myHSC and lyHSC based on intensity of CD150 expression. **E-F.** Gating strategy **(E),** CD150^high/low^ enrichment analysis **(F)** and vWF Tomato fluorescence intensity **(G)** of vWF-Tomato^high^ LT-HSC in young and aged WT/Sema4AKO vWF-Tomato+ mice (n=3-4 per group). **H.** Absolute numbers of myHSC and lyHSC in young and aged WT mice (n=4-6 mice per group). **I.** Representative plots of myHSC quantification in aged WT/Sema4KO mice. **J.** Absolute number of myHSC and lyHSC in individual aged WT/Sema4AKO mice. P values are shown. Statistical significance was assessed by two-tailed *t*-test. Mean +/- (SEM) are shown.

**Extended data Fig 2. (related to Fig 2).**
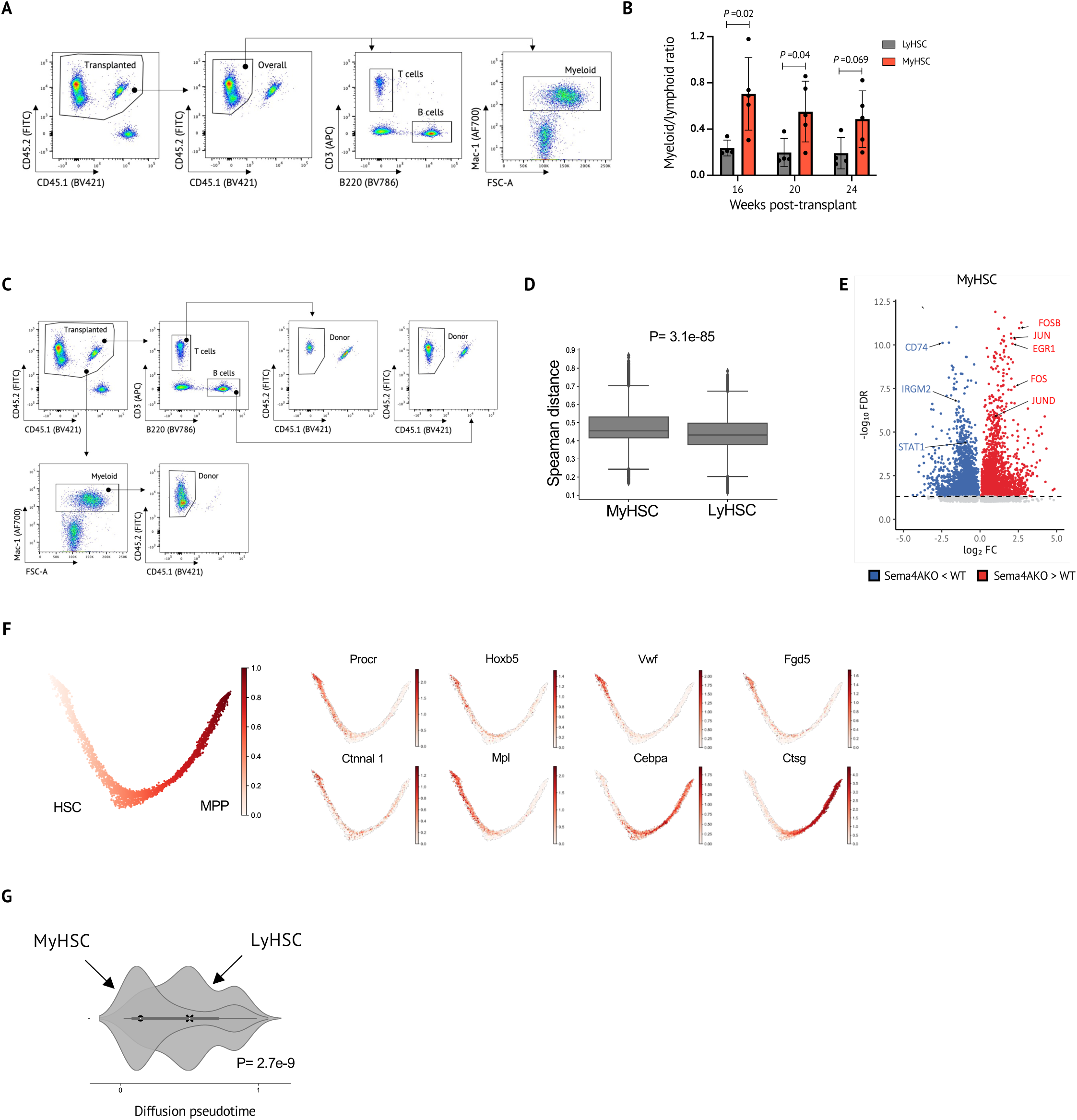
**A.** Gating strategy for quantification of overall reconstitution by donor-derived HSC and analysis of lineage composition of donor-derived graft. **B.** Myeloid/lymphoid ratio of peripheral blood donor-derived cells (CD45.2) in WT mice (CD45.1) which were competitively transplanted with aged WT myHSC or aged WT lyHSC (CD45.2)(n=5 animals per group). **C.** Gating strategy for quantification of lineage contribution by donor-derived cells. **D.** Distribution of pairwise Spearman’s correlation distances between aged WT and Sema4AKO myHSC (left) and lyHSC (right). **E.** Volcano plots showing DEGs in myHSC from aged WT and Sema4KO mice. The *x* and *y* axes indicate the expression fold change (FC) (log_2_) and the false discovery rate (FDR) (−log_10_) for each gene versus controls, respectively. Legends highlight upregulated (red) or downregulated (blue) transcripts, as well as genes not passing cutoff criteria for FC (black) and FDR (gray). Selected representative genes are shown (n=2 biological replicates per genotype). **F.** Expression of HSC/MPP marker genes along the DPT trajectory. Cells are colored based on their normalized expression levels for each the gene indicated at the top. **G.** Distribution of diffusion pseudotime values of myHSC and lyHSC from aged WT mice. P values are shown. Statistical significance was assessed by two-tailed *t*-test (*P<0.05, **P<0.05), except for the diffusion pseudotime analysis where Wilcoxon rank sum test was used. Mean +/- SEM are shown.

**Extended Data Fig 3. (related to Fig 3).**
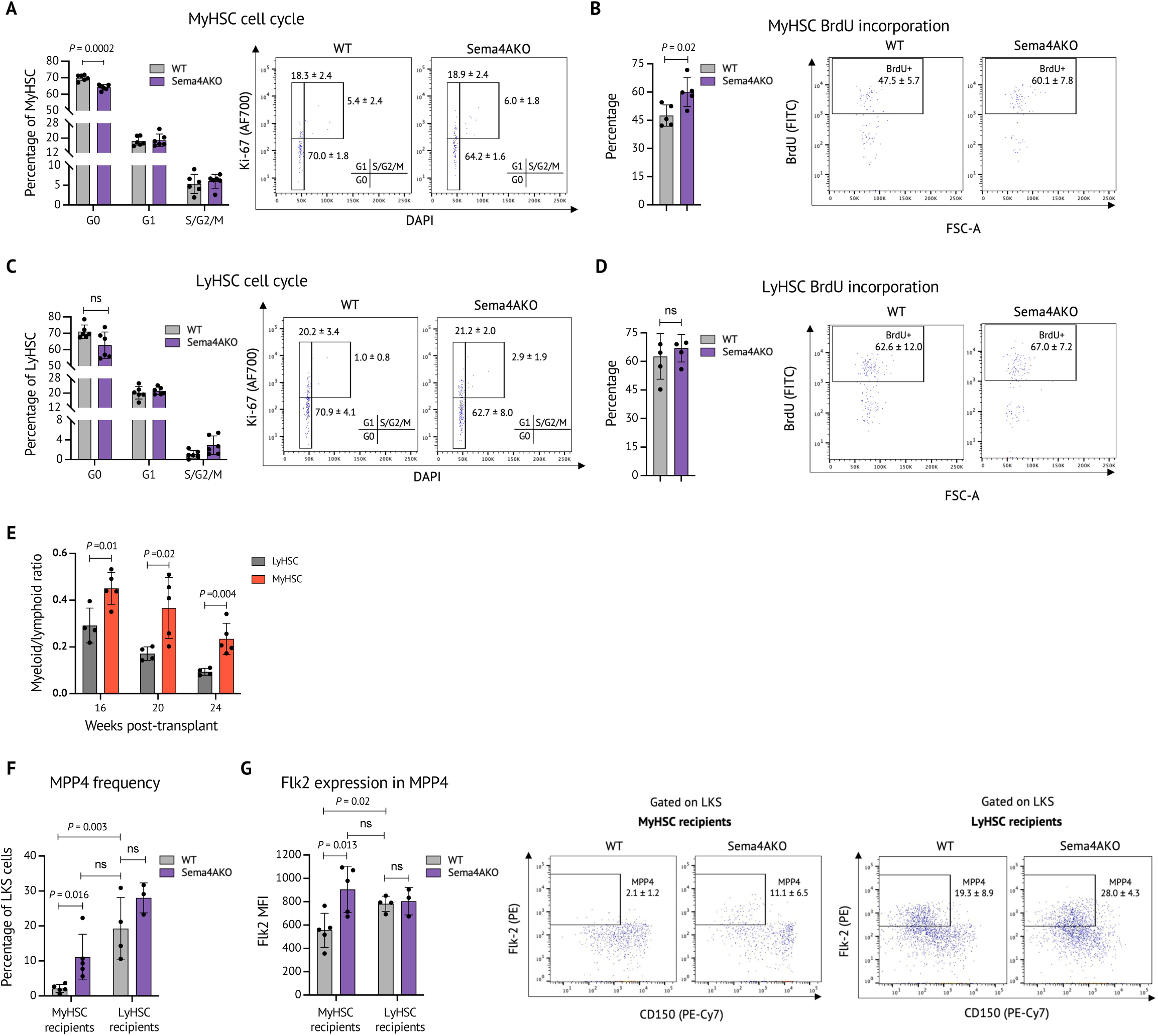
**A, B.** MyHSC cell cycle analysis **(A)** and myHSC short-term (4-day) BrdU incorporation **(B)** in WT/Sema4AKO mice. **C, D.** LyHSC cell cycle analysis **(C)** and myHSC short-term (4-day) BrdU incorporation **(D)** in WT/Sema4AKO mice (n=4-6 mice per group). **E.** Myeloid/lymphoid ratio of peripheral blood donor-derived cells in WT mice (CD45.1) which were competitively transplanted with WT myHSC or LyHSC (CD45.2). **F, G.** Frequency **(F)** and mean fluorescent intensity of Flk2 expression **(G)** of MPP4 in the bone marrow of WT mice (CD45.1) after competitive transplantation of WT/Sema4AKO myHSC (myHSC recipients) (CD45.2) and WT/Sema4AKO lyHSC (lyHSC recipients) (CD45.2), quantification and representative plots are shown for MPP4 frequency (n=4-5 mice per group). P values are shown. Statistical significance was assessed by two-tailed *t*-test.

**Extended Data Fig 4. (related to Fig 4).**
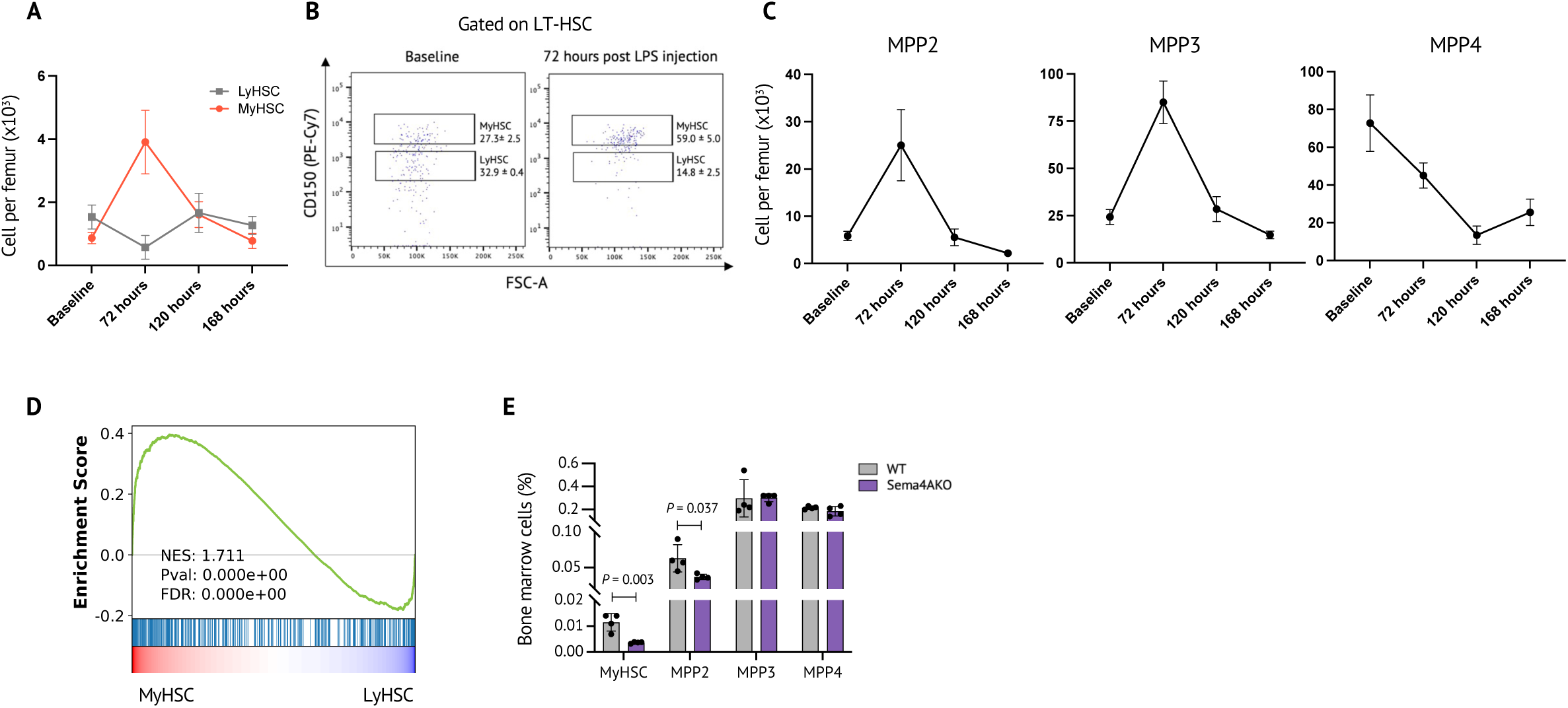
**A.** Absolute number of WT myHSC/lyHSC at baseline and post-LPS injection at indicated time points (n=4-7 mice per group). **B.** Representative flow cytometry plots of myHSC at baseline and 72 hours after LPS injection. **C.** Absolute number of WT MPPs at baseline and post-LPS injection at indicated time points (n=4-7 mice per group). **D.** Enrichment analysis of transcriptional profile of WT CD150^high^ HSC isolated from LPS-injected WT mice for the previously published myHSC signature. **E.** Frequency of myHSC and MPPs in WT/Sema4AKO mice 72 hours post-LPS injection. P values are shown. *P<0.05, **P<0.05. Statistical significance was assessed by two-tailed *t*-test. Mean +/- SEM are shown.

**Extended Data Fig 5. (related to Fig 5).**
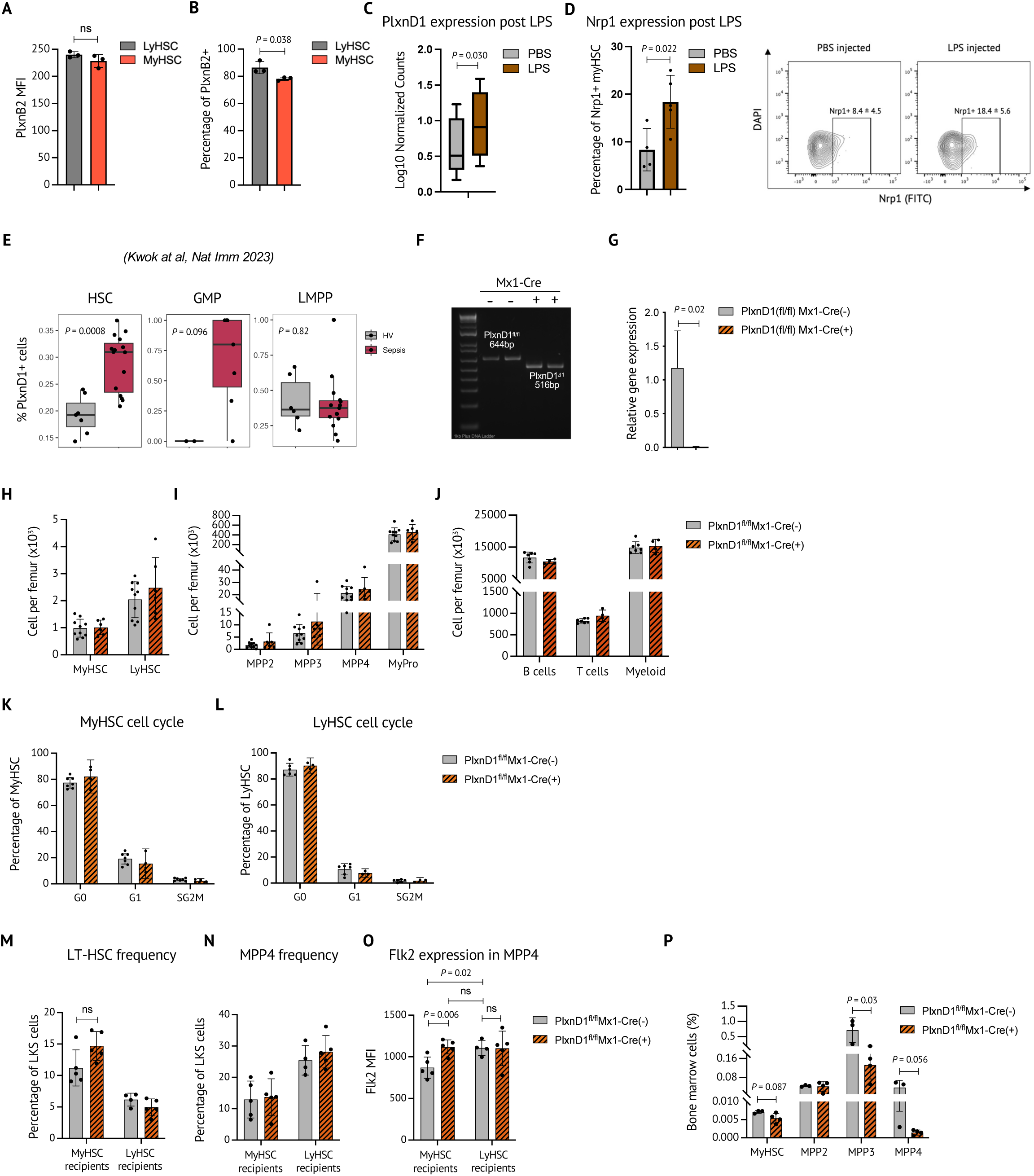
**A.** Expression of PlxnB2 in myHSC and lyHSC in WT mice (n=3 biological replicates). **B.** Frequency of PlxnB2-expressing myHSC and lyHSC in WT mice (n=3 biological replicates). **C, D.** Expression of PlxnD1 mRNA in WT myHSC **(C)** and percentage of Nrp1-expressing WT myHSC **(D)** 72 hours post-LPS injection, as assessed by RNA-Seq and flow cytometry, respectively. **E.** Frequency of PlxnD1^+^ CD34^+^ hematopoietic stem/progenitor cells in patients with sepsis, as assessed by scRNA Seq in Kwok et al.^69^ **F, G.** PlxnD1 excision validation in Lin^-^Kit^+^Sca1^+^ cells by genomic DNA PCR **(F)** and Q-PCR **(G)**. **H-L.** Absolute number of myHSC and lyHSC **(H)**, other primitive hematopoietic cells **(I)**, mature cells **(J),** myHSC cell cycle analysis **(K),** lyHSC cell cycle analysis **(L)** in PlxnD1fl/fl Mx1Cre(+) and PlxnD1fl/fl Cre(-) mice at baseline (n=3-10 mice per group). **M.** Frequency of LT-HSC in the bone marrow of WT mice (CD45.1) after competitive transplantation of PlxnD1fl/fl Mx1Cre(+) and PlxnD1fl/fl Cre(-) myHSC (myHSC recipients) (CD45.2) and PlxnD1fl/fl Mx1Cre(+) and PlxnD1fl/fl Cre(-) lyHSC (lyHSC recipients) (CD45.2) (n=4-5 mice per group). **N, O.** Frequency **(N)** and mean fluorescent intensity of Flk2 expression **(O)** of MPP4 in the bone marrow of WT mice (CD45.1) after competitive transplantation of PlxnD1fl/fl Mx1Cre(+) and PlxnD1fl/fl Cre(-) myHSC (myHSC recipients) and PlxnD1fl/fl Mx1Cre(+) and PlxnD1fl/fl Cre(-) lyHSC (lyHSC recipients), quantification and representative plots are shown for MPP4 frequency (n=4-5 mice per group). **P.** Frequency of myHSC and MPPs in PlxnD1fl/fl Mx1Cre(+) and PlxnD1fl/fl Cre(-) mice 72 hours post-LPS injection. P values are shown. Statistical significance was assessed by two-tailed *t*-test, except for the diffusion pseudotime analysis (P) where Wilcoxon rank sum test was used. Mean +/- SEM are shown.

**Extended Data Fig 6. (related to Fig 6).**
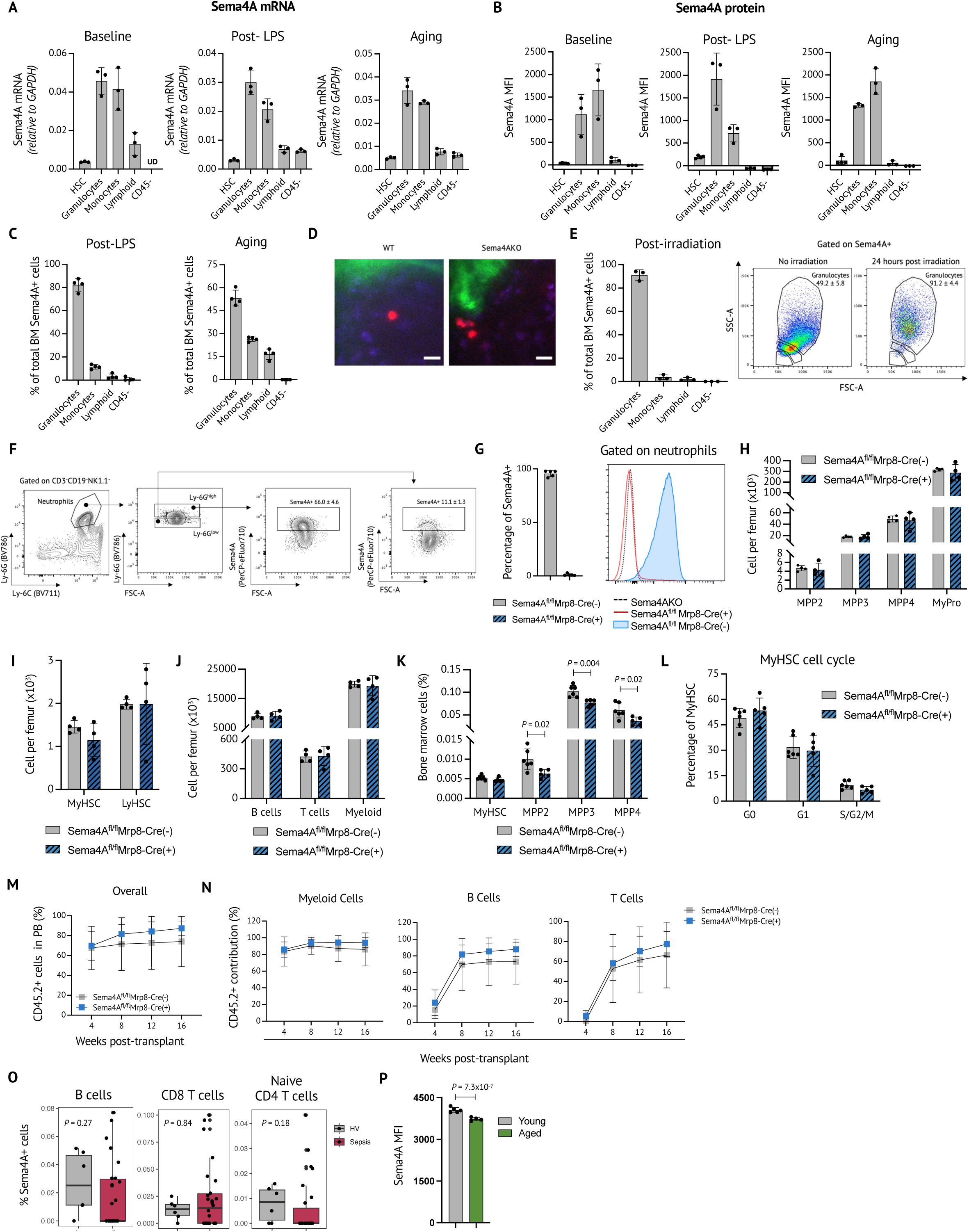
**A, B.** Sema4A mRNA **(A)** and protein expression **(B)** at baseline, 24 hours after LPS injection and upon aging (n=3 mice per group per condition). **C.** Relative contribution of distinct cellular subsets to Sema4A production in the bone marrow 24 hours after LPS injection and upon aging (n=3-4 mice per group)**. D.** Representative calviarial intra-vital microscopy images of WT myHSC transplanted into lethally irradiated WT/Sema4AKO recipients. Red – progeny of transplanted myHSC, green – bone, blue – collagen. Scale bar – 10 microne. **E.** Relative contribution of distinct cellular subsets to Sema4A production in the bone marrow 24 hours after 950 cGy irradiation, as estimated by flow cytometry with Sema4A antibody (n=3 mice per group). Bar graph and representative flow cytometry plots are shown. **F.** Gating strategy for assessing Sema4A expression in Ly6G^high^ vs Ly6G^low^ neutrophils. **G.** Quantification and representative histogram of Sema4A deletion from neutrophils in Sema4A^fl/fl^ Mrp8-Cre(+) mice (n=5 mice per group). Sema4A antibody-stained neutrophils from Sema4AKO mouse were used as a negative control. **H-J.** Absolute numer of primitive hematopoietic cells **(H)**, myHSC and lyHSC **(I)** and mature cells **(J)** at baseline in Sema4A^fl/fl^ Mrp8-Cre(+) and Sema4A^fl/fl^ Mrp8-Cre(-) mice (n=5 mice per group). **K.** Frequency of myHSC and MPPs in Sema4A^fl/fl^ Mrp8-Cre(+) and Sema4A^fl/fl^ Mrp8-Cre(-) mice 72 hours after LPS injection (n=4-7 mice per group). **L.** Cell cycle analysis of myHSC from Sema4A^fl/fl^ Mrp8-Cre(+) and Sema4A^fl/fl^ Mrp8-Cre(-) mice 72 hours after LPS injection (n=4-7 mice per group). **M, N.** Overall percentage of donor-derived peripheral blood cells **(M)** and lineage contribution by donor-derived cells **(N)** in WT mice (CD45.1) which were competitively transplanted with myHSC from low-dose LPS-treated Sema4A^fl/fl^ Mrp8-Cre(+) and Sema4A^fl/fl^ Mrp8-Cre(-) (CD45.2) mice (n=5 mice per group). **O.** Frequency of Sema4A^+^ cells in peripheral blood lymphoid subsets from patients with sepsis and healthy volunteers (HV), as assessed by single cell RNA-Seq in Kwok et al.^69^ **P.** Sema4A mean fluorescent intensity of Sema4A^+^Ly6G^high^ neutrophils in the bone marrow of young and aged WT mice (n=4-5 mice per group). P values are shown. Statistical significance was assessed by two-tailed *t*-test. Mean +/- SEM are shown.

## References

1. Scaramozza A, Park D, Kollu S, Beerman I, Sun X, Rossi DJ, Lin CP, Scadden DT, Crist C, Brack AS. Lineage Tracing Reveals a Subset of Reserve Muscle Stem Cells Capable of Clonal Expansion under Stress. Cell Stem Cell. 2019;24(6):944–57 e5. Epub 2019/04/23. doi: 10.1016/j.stem.2019.03.020. PubMed PMID: 31006621; PMCID: PMC6597014.

2. Ibrayeva A, Bay M, Pu E, Jorg DJ, Peng L, Jun H, Zhang N, Aaron D, Lin C, Resler G, Hidalgo A, Jang MH, Simons BD, Bonaguidi MA. Early stem cell aging in the mature brain. Cell Stem Cell. 2021;28(5):955–66 e7. Epub 20210412. doi: 10.1016/j.stem.2021.03.018. PubMed PMID: 33848469; PMCID: PMC10069280.

3. Altshuler A, Amitai-Lange A, Tarazi N, Dey S, Strinkovsky L, Hadad-Porat S, Bhattacharya S, Nasser W, Imeri J, Ben-David G, Abboud-Jarrous G, Tiosano B, Berkowitz E, Karin N, Savir Y, Shalom-Feuerstein R. Discrete limbal epithelial stem cell populations mediate corneal homeostasis and wound healing. Cell Stem Cell. 2021;28(7):1248–61 e8. Epub 2021/05/14. doi: 10.1016/j.stem.2021.04.003. PubMed PMID: 33984282; PMCID: PMC8254798.

4. Farrelly O, Suzuki-Horiuchi Y, Brewster M, Kuri P, Huang S, Rice G, Bae H, Xu J, Dentchev T, Lee V, Rompolas P. Two-photon live imaging of single corneal stem cells reveals compartmentalized organization of the limbal niche. Cell Stem Cell. 2021;28(7):1233–47 e4. Epub 2021/05/14. doi: 10.1016/j.stem.2021.02.022. PubMed PMID: 33984283.

5. Hsu YC, Pasolli HA, Fuchs E. Dynamics between stem cells, niche, and progeny in the hair follicle. Cell. 2011;144(1):92–105. Epub 2011/01/11. doi: 10.1016/j.cell.2010.11.049. PubMed PMID: 21215372; PMCID: PMC3050564.

6. Matatall KA, Shen CC, Challen GA, King KY. Type II interferon promotes differentiation of myeloid-biased hematopoietic stem cells. Stem Cells. 2014;32(11):3023–30. Epub 2014/08/01. doi: 10.1002/stem.1799. PubMed PMID: 25078851; PMCID: PMC4198460.

7. Yamamoto R, Wilkinson AC, Nakauchi H. Changing concepts in hematopoietic stem cells. Science. 2018;362(6417):895-6. doi: 10.1126/science.aat7873. PubMed PMID: 30467158.

8. Yamamoto R, Morita Y, Ooehara J, Hamanaka S, Onodera M, Rudolph KL, Ema H, Nakauchi H. Clonal analysis unveils self-renewing lineage-restricted progenitors generated directly from hematopoietic stem cells. Cell. 2013;154(5):1112–26. doi: 10.1016/j.cell.2013.08.007. PubMed PMID: 23993099.

9. Florez MA, Matatall KA, Jeong Y, Ortinau L, Shafer PW, Lynch AM, Jaksik R, Kimmel M, Park D, King KY. Interferon Gamma Mediates Hematopoietic Stem Cell Activation and Niche Relocalization through BST2. Cell Rep. 2020;33(12):108530. doi: 10.1016/j.celrep.2020.108530. PubMed PMID: 33357430; PMCID: PMC7816211.

10. Beerman I, Bhattacharya D, Zandi S, Sigvardsson M, Weissman IL, Bryder D, Rossi DJ. Functionally distinct hematopoietic stem cells modulate hematopoietic lineage potential during aging by a mechanism of clonal expansion. Proc Natl Acad Sci U S A. 2010;107(12):5465–70. Epub 20100318. doi: 10.1073/pnas.1000834107. PubMed PMID: 20304793; PMCID: PMC2851806.

11. Challen GA, Boles NC, Chambers SM, Goodell MA. Distinct hematopoietic stem cell subtypes are differentially regulated by TGF-beta1. Cell Stem Cell. 2010;6(3):265–78. doi: 10.1016/j.stem.2010.02.002. PubMed PMID: 20207229; PMCID: PMC2837284.

12. Sanjuan-Pla A, Macaulay IC, Jensen CT, Woll PS, Luis TC, Mead A, Moore S, Carella C, Matsuoka S, Bouriez Jones T, Chowdhury O, Stenson L, Lutteropp M, Green JC, Facchini R, Boukarabila H, Grover A, Gambardella A, Thongjuea S, Carrelha J, Tarrant P, Atkinson D, Clark SA, Nerlov C, Jacobsen SE. Platelet-biased stem cells reside at the apex of the haematopoietic stem-cell hierarchy. Nature. 2013;502(7470):232-6. Epub 2013/08/13. doi: 10.1038/nature12495. PubMed PMID: 23934107.

13. Carrelha J, Meng Y, Kettyle LM, Luis TC, Norfo R, Alcolea V, Boukarabila H, Grasso F, Gambardella A, Grover A, Hogstrand K, Lord AM, Sanjuan-Pla A, Woll PS, Nerlov C, Jacobsen SEW. Hierarchically related lineage-restricted fates of multipotent haematopoietic stem cells. Nature. 2018;554(7690):106-11. Epub 2018/01/04. doi: 10.1038/nature25455. PubMed PMID: 29298288.

14. Notta F, Zandi S, Takayama N, Dobson S, Gan OI, Wilson G, Kaufmann KB, McLeod J, Laurenti E, Dunant CF, McPherson JD, Stein LD, Dror Y, Dick JE. Distinct routes of lineage development reshape the human blood hierarchy across ontogeny. Science. 2016;351(6269):aab2116. Epub 20151105. doi: 10.1126/science.aab2116. PubMed PMID: 26541609; PMCID: PMC4816201.

15. Pang WW, Price EA, Sahoo D, Beerman I, Maloney WJ, Rossi DJ, Schrier SL, Weissman IL. Human bone marrow hematopoietic stem cells are increased in frequency and myeloid-biased with age. Proc Natl Acad Sci U S A. 2011;108(50):20012–7. Epub 2011/11/30. doi: 10.1073/pnas.1116110108. PubMed PMID: 22123971; PMCID: PMC3250139.

16. Mitroulis I, Ruppova K, Wang B, Chen LS, Grzybek M, Grinenko T, Eugster A, Troullinaki M, Palladini A, Kourtzelis I, Chatzigeorgiou A, Schlitzer A, Beyer M, Joosten LAB, Isermann B, Lesche M, Petzold A, Simons K, Henry I, Dahl A, Schultze JL, Wielockx B, Zamboni N, Mirtschink P, Coskun U, Hajishengallis G, Netea MG, Chavakis T. Modulation of Myelopoiesis Progenitors Is an Integral Component of Trained Immunity. Cell. 2018;172(1-2):147–61 e12. Epub 2018/01/13. doi: 10.1016/j.cell.2017.11.034. PubMed PMID: 29328910; PMCID: PMC5766828.

17. Bogeska R, Mikecin AM, Kaschutnig P, Fawaz M, Buchler-Schaff M, Le D, Ganuza M, Vollmer A, Paffenholz SV, Asada N, Rodriguez-Correa E, Frauhammer F, Buettner F, Ball M, Knoch J, Stable S, Walter D, Petri A, Carreno-Gonzalez MJ, Wagner V, Brors B, Haas S, Lipka DB, Essers MAG, Weru V, Holland-Letz T, Mallm JP, Rippe K, Kramer S, Schlesner M, McKinney Freeman S, Florian MC, King KY, Frenette PS, Rieger MA, Milsom MD. Inflammatory exposure drives long-lived impairment of hematopoietic stem cell self-renewal activity and accelerated aging. Cell Stem Cell. 2022;29(8):1273–84 e8. Epub 20220719. doi: 10.1016/j.stem.2022.06.012. PubMed PMID: 35858618; PMCID: PMC9357150.

18. Goodell MA, Rando TA. Stem cells and healthy aging. Science. 2015;350(6265):1199–204. doi: 10.1126/science.aab3388. PubMed PMID: 26785478.

19. Mann M, Mehta A, de Boer CG, Kowalczyk MS, Lee K, Haldeman P, Rogel N, Knecht AR, Farouq D, Regev A, Baltimore D. Heterogeneous Responses of Hematopoietic Stem Cells to Inflammatory Stimuli Are Altered with Age. Cell Rep. 2018;25(11):2992–3005 e5. Epub 2018/12/13. doi: 10.1016/j.celrep.2018.11.056. PubMed PMID: 30540934; PMCID: PMC6424521.

20. Chen X, Deng H, Churchill MJ, Luchsinger LL, Du X, Chu TH, Friedman RA, Middelhoff M, Ding H, Tailor YH, Wang ALE, Liu H, Niu Z, Wang H, Jiang Z, Renders S, Ho SH, Shah SV, Tishchenko P, Chang W, Swayne TC, Munteanu L, Califano A, Takahashi R, Nagar KK, Renz BW, Worthley DL, Westphalen CB, Hayakawa Y, Asfaha S, Borot F, Lin CS, Snoeck HW, Mukherjee S, Wang TC. Bone Marrow Myeloid Cells Regulate Myeloid-Biased Hematopoietic Stem Cells via a Histamine-Dependent Feedback Loop. Cell Stem Cell. 2017;21(6):747–60 e7. Epub 2017/12/05. doi: 10.1016/j.stem.2017.11.003. PubMed PMID: 29198940; PMCID: PMC5975960.

21. Pinho S, Marchand T, Yang E, Wei Q, Nerlov C, Frenette PS. Lineage-Biased Hematopoietic Stem Cells Are Regulated by Distinct Niches. Dev Cell. 2018;44(5):634–41 e4. Epub 20180215. doi: 10.1016/j.devcel.2018.01.016. PubMed PMID: 29456137; PMCID: PMC5886750.

22. Alto LT, Terman JR. Semaphorins and their Signaling Mechanisms. Methods Mol Biol. 2017;1493:1–25. Epub 2016/10/28. doi: 10.1007/978-1-4939-6448-2_1. PubMed PMID: 27787839; PMCID: PMC5538787.

23. Kumanogoh A, Marukawa S, Suzuki K, Takegahara N, Watanabe C, Ch’ng E, Ishida I, Fujimura H, Sakoda S, Yoshida K, Kikutani H. Class IV semaphorin Sema4A enhances T-cell activation and interacts with Tim-2. Nature. 2002;419(6907):629-33. doi: 10.1038/nature01037. PubMed PMID: 12374982.

24. Kumanogoh A, Shikina T, Suzuki K, Uematsu S, Yukawa K, Kashiwamura S, Tsutsui H, Yamamoto M, Takamatsu H, Ko-Mitamura EP, Takegahara N, Marukawa S, Ishida I, Morishita H, Prasad DV, Tamura M, Mizui M, Toyofuku T, Akira S, Takeda K, Okabe M, Kikutani H. Nonredundant roles of Sema4A in the immune system: defective T cell priming and Th1/Th2 regulation in Sema4A-deficient mice. Immunity. 2005;22(3):305–16. Epub 2005/03/23. doi: 10.1016/j.immuni.2005.01.014. PubMed PMID: 15780988.

25. Toyofuku T, Yabuki M, Kamei J, Kamei M, Makino N, Kumanogoh A, Hori M. Semaphorin-4A, an activator for T-cell-mediated immunity, suppresses angiogenesis via Plexin-D1. EMBO J. 2007;26(5):1373–84. Epub 2007/02/24. doi: 10.1038/sj.emboj.7601589. PubMed PMID: 17318185; PMCID: PMC1817636.

26. Silberstein L, Goncalves KA, Kharchenko PV, Turcotte R, Kfoury Y, Mercier F, Baryawno N, Severe N, Bachand J, Spencer JA, Papazian A, Lee D, Chitteti BR, Srour EF, Hoggatt J, Tate T, Lo Celso C, Ono N, Nutt S, Heino J, Sipila K, Shioda T, Osawa M, Lin CP, Hu GF, Scadden DT. Proximity-Based Differential Single-Cell Analysis of the Niche to Identify Stem/Progenitor Cell Regulators. Cell Stem Cell. 2016;19(4):530–43. Epub 2016/08/16. doi: 10.1016/j.stem.2016.07.004. PubMed PMID: 27524439; PMCID: PMC5402355.

27. Kharchenko PV, Silberstein L, Scadden DT. Bayesian approach to single-cell differential expression analysis. Nat Methods. 2014;11(7):740–2. Epub 20140518. doi: 10.1038/nmeth.2967. PubMed PMID: 24836921; PMCID: PMC4112276.

28. Feng CG, Weksberg DC, Taylor GA, Sher A, Goodell MA. The p47 GTPase Lrg-47 (Irgm1) links host defense and hematopoietic stem cell proliferation. Cell Stem Cell. 2008;2(1):83–9. doi: 10.1016/j.stem.2007.10.007. PubMed PMID: 18371424; PMCID: PMC2278017.

29. Goncalves KA, Silberstein L, Li S, Severe N, Hu MG, Yang H, Scadden DT, Hu GF. Angiogenin Promotes Hematopoietic Regeneration by Dichotomously Regulating Quiescence of Stem and Progenitor Cells. Cell. 2016;166(4):894–906. Epub 2016/08/16. doi: 10.1016/j.cell.2016.06.042. PubMed PMID: 27518564; PMCID: PMC4988404.

30. Takamatsu H, Kumanogoh A. Diverse roles for semaphorin-plexin signaling in the immune system. Trends Immunol. 2012;33(3):127–35. Epub 20120209. doi: 10.1016/j.it.2012.01.008. PubMed PMID: 22325954.

31. Delgoffe GM, Woo SR, Turnis ME, Gravano DM, Guy C, Overacre AE, Bettini ML, Vogel P, Finkelstein D, Bonnevier J, Workman CJ, Vignali DA. Stability and function of regulatory T cells is maintained by a neuropilin-1-semaphorin-4a axis. Nature. 2013;501(7466):252-6. Epub 20130804. doi: 10.1038/nature12428. PubMed PMID: 23913274; PMCID: PMC3867145.

32. WHO Classification of Tumours Haematolymphoid Tumours. 5 ed2024.

33. Ho YH, Del Toro R, Rivera-Torres J, Rak J, Korn C, Garcia-Garcia A, Macias D, Gonzalez-Gomez C, Del Monte A, Wittner M, Waller AK, Foster HR, Lopez-Otin C, Johnson RS, Nerlov C, Ghevaert C, Vainchenker W, Louache F, Andres V, Mendez-Ferrer S. Remodeling of Bone Marrow Hematopoietic Stem Cell Niches Promotes Myeloid Cell Expansion during Premature or Physiological Aging. Cell Stem Cell. 2019;25(3):407–18 e6. Epub 20190711. doi: 10.1016/j.stem.2019.06.007. PubMed PMID: 31303548; PMCID: PMC6739444.

34. Pronk CJ, Rossi DJ, Mansson R, Attema JL, Norddahl GL, Chan CK, Sigvardsson M, Weissman IL, Bryder D. Elucidation of the phenotypic, functional, and molecular topography of a myeloerythroid progenitor cell hierarchy. Cell Stem Cell. 2007;1(4):428–42. doi: 10.1016/j.stem.2007.07.005. PubMed PMID: 18371379.

35. Grover A, Sanjuan-Pla A, Thongjuea S, Carrelha J, Giustacchini A, Gambardella A, Macaulay I, Mancini E, Luis TC, Mead A, Jacobsen SE, Nerlov C. Single-cell RNA sequencing reveals molecular and functional platelet bias of aged haematopoietic stem cells. Nat Commun. 2016;7:11075. Epub 2016/03/25. doi: 10.1038/ncomms11075. PubMed PMID: 27009448; PMCID: PMC4820843.

36. Liang Y, Van Zant G, Szilvassy SJ. Effects of aging on the homing and engraftment of murine hematopoietic stem and progenitor cells. Blood. 2005;106(4):1479–87. Epub 20050412. doi: 10.1182/blood-2004-11-4282. PubMed PMID: 15827136; PMCID: PMC1895199.

37. Picelli S, Faridani OR, Bjorklund AK, Winberg G, Sagasser S, Sandberg R. Full-length RNA-seq from single cells using Smart-seq2. Nat Protoc. 2014;9(1):171–81. Epub 2014/01/05. doi: 10.1038/nprot.2014.006. PubMed PMID: 24385147.

38. Karakaslar EO, Katiyar N, Hasham M, Youn A, Sharma S, Chung CH, Marches R, Korstanje R, Banchereau J, Ucar D. Transcriptional activation of Jun and Fos members of the AP-1 complex is a conserved signature of immune aging that contributes to inflammaging. Aging Cell. 2023;22(4):e13792. Epub 20230224. doi: 10.1111/acel.13792. PubMed PMID: 36840360; PMCID: PMC10086525.

39. Trizzino M, Zucco A, Deliard S, Wang F, Barbieri E, Veglia F, Gabrilovich D, Gardini A. EGR1 is a gatekeeper of inflammatory enhancers in human macrophages. Sci Adv. 2021;7(3). Epub 20210113. doi: 10.1126/sciadv.aaz8836. PubMed PMID: 33523892; PMCID: PMC7806227.

40. Jia D, Chen S, Bai P, Luo C, Liu J, Sun A, Ge J. Cardiac Resident Macrophage-Derived Legumain Improves Cardiac Repair by Promoting Clearance and Degradation of Apoptotic Cardiomyocytes After Myocardial Infarction. Circulation. 2022;145(20):1542–56. Epub 20220418. doi: 10.1161/CIRCULATIONAHA.121.057549. PubMed PMID: 35430895.

41. Marzio R, Jirillo E, Ransijn A, Mauel J, Corradin SB. Expression and function of the early activation antigen CD69 in murine macrophages. J Leukoc Biol. 1997;62(3):349–55. doi: 10.1002/jlb.62.3.349. PubMed PMID: 9307073.

42. Konturek-Ciesla A, Olofzon R, Kharazi S, Bryder D. Implications of stress-induced gene expression for hematopoietic stem cell aging studies. Nature Aging. 2024;4(2):177–84. doi: 10.1038/s43587-023-00558-z.

43. Yoshida H, Lareau CA, Ramirez RN, Rose SA, Maier B, Wroblewska A, Desland F, Chudnovskiy A, Mortha A, Dominguez C, Tellier J, Kim E, Dwyer D, Shinton S, Nabekura T, Qi Y, Yu B, Robinette M, Kim KW, Wagers A, Rhoads A, Nutt SL, Brown BD, Mostafavi S, Buenrostro JD, Benoist C, Immunological Genome P. The cis-Regulatory Atlas of the Mouse Immune System. Cell. 2019;176(4):897–912 e20. Epub 20190124. doi: 10.1016/j.cell.2018.12.036. PubMed PMID: 30686579; PMCID: PMC6785993.

44. King KY, Baldridge MT, Weksberg DC, Chambers SM, Lukov GL, Wu S, Boles NC, Jung SY, Qin J, Liu D, Songyang Z, Eissa NT, Taylor GA, Goodell MA. Irgm1 protects hematopoietic stem cells by negative regulation of IFN signaling. Blood. 2011;118(6):1525–33. Epub 20110601. doi: 10.1182/blood-2011-01-328682. PubMed PMID: 21633090; PMCID: PMC3156044.

45. Weih F, Carrasco D, Durham SK, Barton DS, Rizzo CA, Ryseck RP, Lira SA, Bravo R. Multiorgan inflammation and hematopoietic abnormalities in mice with a targeted disruption of RelB, a member of the NF-kappa B/Rel family. Cell. 1995;80(2):331–40. doi: 10.1016/0092-8674(95)90416-6. PubMed PMID: 7834753.

46. Li J, Williams MJ, Park HJ, Bastos HP, Wang X, Prins D, Wilson NK, Johnson C, Sham K, Wantoch M, Watcham S, Kinston SJ, Pask DC, Hamilton TL, Sneade R, Waller AK, Ghevaert C, Vassiliou GS, Laurenti E, Kent DG, Gottgens B, Green AR. STAT1 is essential for HSC function and maintains MHCIIhi stem cells that resist myeloablation and neoplastic expansion. Blood. 2022;140(14):1592–606. doi: 10.1182/blood.2021014009. PubMed PMID: 35767701; PMCID: PMC7614316.

47. Becker-Herman S, Rozenberg M, Hillel-Karniel C, Gil-Yarom N, Kramer MP, Barak A, Sever L, David K, Radomir L, Lewinsky H, Levi M, Friedlander G, Bucala R, Peled A, Shachar I. CD74 is a regulator of hematopoietic stem cell maintenance. PLoS Biol. 2021;19(3):e3001121. Epub 2021/03/05. doi: 10.1371/journal.pbio.3001121. PubMed PMID: 33661886; PMCID: PMC7963458.

48. Haghverdi L, Buttner M, Wolf FA, Buettner F, Theis FJ. Diffusion pseudotime robustly reconstructs lineage branching. Nat Methods. 2016;13(10):845–8. Epub 2016/08/30. doi: 10.1038/nmeth.3971. PubMed PMID: 27571553.

49. Nestorowa S, Hamey FK, Pijuan Sala B, Diamanti E, Shepherd M, Laurenti E, Wilson NK, Kent DG, Gottgens B. A single-cell resolution map of mouse hematopoietic stem and progenitor cell differentiation. Blood. 2016;128(8):e20–31. Epub 2016/07/02. doi: 10.1182/blood-2016-05-716480. PubMed PMID: 27365425; PMCID: PMC5305050.

50. Wilson NK, Kent DG, Buettner F, Shehata M, Macaulay IC, Calero-Nieto FJ, Sanchez Castillo M, Oedekoven CA, Diamanti E, Schulte R, Ponting CP, Voet T, Caldas C, Stingl J, Green AR, Theis FJ, Gottgens B. Combined Single-Cell Functional and Gene Expression Analysis Resolves Heterogeneity within Stem Cell Populations. Cell Stem Cell. 2015;16(6):712–24. Epub 20150521. doi: 10.1016/j.stem.2015.04.004. PubMed PMID: 26004780; PMCID: PMC4460190.

51. Kovtonyuk LV, Fritsch K, Feng X, Manz MG, Takizawa H. Inflamm-Aging of Hematopoiesis, Hematopoietic Stem Cells, and the Bone Marrow Microenvironment. Front Immunol. 2016;7:502. Epub 2016/11/30. doi: 10.3389/fimmu.2016.00502. PubMed PMID: 27895645; PMCID: PMC5107568.

52. Flohr Svendsen A, Yang D, Kim K, Lazare S, Skinder N, Zwart E, Mura-Meszaros A, Ausema A, von Eyss B, de Haan G, Bystrykh L. A comprehensive transcriptome signature of murine hematopoietic stem cell aging. Blood. 2021;138(6):439–51. doi: 10.1182/blood.2020009729. PubMed PMID: 33876187.

53. Cabezas-Wallscheid N, Klimmeck D, Hansson J, Lipka DB, Reyes A, Wang Q, Weichenhan D, Lier A, von Paleske L, Renders S, Wunsche P, Zeisberger P, Brocks D, Gu L, Herrmann C, Haas S, Essers MAG, Brors B, Eils R, Huber W, Milsom MD, Plass C, Krijgsveld J, Trumpp A. Identification of regulatory networks in HSCs and their immediate progeny via integrated proteome, transcriptome, and DNA methylome analysis. Cell Stem Cell. 2014;15(4):507–22. Epub 2014/08/28. doi: 10.1016/j.stem.2014.07.005. PubMed PMID: 25158935.

54. Laurenti E, Frelin C, Xie S, Ferrari R, Dunant CF, Zandi S, Neumann A, Plumb I, Doulatov S, Chen J, April C, Fan JB, Iscove N, Dick JE. CDK6 levels regulate quiescence exit in human hematopoietic stem cells. Cell Stem Cell. 2015;16(3):302–13. Epub 2015/02/24. doi: 10.1016/j.stem.2015.01.017. PubMed PMID: 25704240; PMCID: PMC4359055.

55. Fernando RN, Eleuteri B, Abdelhady S, Nussenzweig A, Andang M, Ernfors P. Cell cycle restriction by histone H2AX limits proliferation of adult neural stem cells. Proc Natl Acad Sci U S A. 2011;108(14):5837–42. Epub 20110321. doi: 10.1073/pnas.1014993108. PubMed PMID: 21436033; PMCID: PMC3078396.

56. Qian H, Buza-Vidas N, Hyland CD, Jensen CT, Antonchuk J, Mansson R, Thoren LA, Ekblom M, Alexander WS, Jacobsen SE. Critical role of thrombopoietin in maintaining adult quiescent hematopoietic stem cells. Cell Stem Cell. 2007;1(6):671–84. Epub 20071120. doi: 10.1016/j.stem.2007.10.008. PubMed PMID: 18371408.

57. de Laval B, Maurizio J, Kandalla PK, Brisou G, Simonnet L, Huber C, Gimenez G, Matcovitch-Natan O, Reinhardt S, David E, Mildner A, Leutz A, Nadel B, Bordi C, Amit I, Sarrazin S, Sieweke MH. C/EBPbeta-Dependent Epigenetic Memory Induces Trained Immunity in Hematopoietic Stem Cells. Cell Stem Cell. 2020;26(5):657–74 e8. Epub 20200312. doi: 10.1016/j.stem.2020.01.017. PubMed PMID: 32169166.

58. Collins A, Mitchell CA, Passegue E. Inflammatory signaling regulates hematopoietic stem and progenitor cell development and homeostasis. J Exp Med. 2021;218(7). Epub 20210615. doi: 10.1084/jem.20201545. PubMed PMID: 34129018; PMCID: PMC8210624.

59. Luchsinger LL, de Almeida MJ, Corrigan DJ, Mumau M, Snoeck HW. Mitofusin 2 maintains haematopoietic stem cells with extensive lymphoid potential. Nature. 2016;529(7587):528-31. Epub 20160120. doi: 10.1038/nature16500. PubMed PMID: 26789249; PMCID: PMC5106870.

60. Morita Y, Ema H, Nakauchi H. Heterogeneity and hierarchy within the most primitive hematopoietic stem cell compartment. J Exp Med. 2010;207(6):1173–82. Epub 2010/04/28. doi: 10.1084/jem.20091318. PubMed PMID: 20421392; PMCID: PMC2882827.

61. Pietras EM, Reynaud D, Kang YA, Carlin D, Calero-Nieto FJ, Leavitt AD, Stuart JM, Gottgens B, Passegue E. Functionally Distinct Subsets of Lineage-Biased Multipotent Progenitors Control Blood Production in Normal and Regenerative Conditions. Cell Stem Cell. 2015;17(1):35–46. Epub 20150618. doi: 10.1016/j.stem.2015.05.003. PubMed PMID: 26095048; PMCID: PMC4542150.

62. Adolfsson J, Mansson R, Buza-Vidas N, Hultquist A, Liuba K, Jensen CT, Bryder D, Yang L, Borge OJ, Thoren LA, Anderson K, Sitnicka E, Sasaki Y, Sigvardsson M, Jacobsen SE. Identification of Flt3+ lympho-myeloid stem cells lacking erythro-megakaryocytic potential a revised road map for adult blood lineage commitment. Cell. 2005;121(2):295–306. doi: 10.1016/j.cell.2005.02.013. PubMed PMID: 15851035.

63. Esplin BL, Shimazu T, Welner RS, Garrett KP, Nie L, Zhang Q, Humphrey MB, Yang Q, Borghesi LA, Kincade PW. Chronic exposure to a TLR ligand injures hematopoietic stem cells. J Immunol. 2011;186(9):5367–75. Epub 2011/03/29. doi: 10.4049/jimmunol.1003438. PubMed PMID: 21441445; PMCID: PMC3086167.

64. Demel UM, Lutz R, Sujer S, Demerdash Y, Sood S, Grunschlager F, Kuck A, Werner P, Blaszkiewicz S, Uckelmann HJ, Haas S, Essers MAG. A complex proinflammatory cascade mediates the activation of HSCs upon LPS exposure in vivo. Blood Adv. 2022;6(11):3513–28. doi: 10.1182/bloodadvances.2021006088. PubMed PMID: 35413096; PMCID: PMC9198917.

65. Passegué E, Wagers AJ, Giuriato S, Anderson WC, Weissman IL. Global analysis of proliferation and cell cycle gene expression in the regulation of hematopoietic stem and progenitor cell fates. J Exp Med. 2005;202(11):1599–611. doi: 10.1084/jem.20050967. PubMed PMID: 16330818; PMCID: PMC2213324.

66. Haltalli MLR, Watcham S, Wilson NK, Eilers K, Lipien A, Ang H, Birch F, Anton SG, Pirillo C, Ruivo N, Vainieri ML, Pospori C, Sinden RE, Luis TC, Langhorne J, Duffy KR, Göttgens B, Blagborough AM, Lo Celso C. Manipulating niche composition limits damage to haematopoietic stem cells during Plasmodium infection. Nat Cell Biol. 2020;22(12):1399–410. Epub 20201123. doi: 10.1038/s41556-020-00601-w. PubMed PMID: 33230302; PMCID: PMC7611033.

67. Walter D, Lier A, Geiselhart A, Thalheimer FB, Huntscha S, Sobotta MC, Moehrle B, Brocks D, Bayindir I, Kaschutnig P, Muedder K, Klein C, Jauch A, Schroeder T, Geiger H, Dick TP, Holland-Letz T, Schmezer P, Lane SW, Rieger MA, Essers MA, Williams DA, Trumpp A, Milsom MD. Exit from dormancy provokes DNA-damage-induced attrition in haematopoietic stem cells. Nature. 2015;520(7548):549-52. Epub 2015/02/25. doi: 10.1038/nature14131. PubMed PMID: 25707806.

68. Komorowska K, Doyle A, Wahlestedt M, Subramaniam A, Debnath S, Chen J, Soneji S, Van Handel B, Mikkola HKA, Miharada K, Bryder D, Larsson J, Magnusson M. Hepatic Leukemia Factor Maintains Quiescence of Hematopoietic Stem Cells and Protects the Stem Cell Pool during Regeneration. Cell Rep. 2017;21(12):3514–23. doi: 10.1016/j.celrep.2017.11.084. PubMed PMID: 29262330.

69. Warr MR, Binnewies M, Flach J, Reynaud D, Garg T, Malhotra R, Debnath J, Passegue E. FOXO3A directs a protective autophagy program in haematopoietic stem cells. Nature. 2013;494(7437):323-7. Epub 20130206. doi: 10.1038/nature11895. PubMed PMID: 23389440; PMCID: PMC3579002.

70. Bott KN, Yumol JL, Comelli EM, Klentrou P, Peters SJ, Ward WE. Trabecular and cortical bone are unaltered in response to chronic lipopolysaccharide exposure via osmotic pumps in male and female CD-1 mice. PLoS One. 2021;16(2):e0243933. Epub 20210205. doi: 10.1371/journal.pone.0243933. PubMed PMID: 33544708; PMCID: PMC7864436.

71. Cabezas-Wallscheid N, Buettner F, Sommerkamp P, Klimmeck D, Ladel L, Thalheimer FB, Pastor-Flores D, Roma LP, Renders S, Zeisberger P, Przybylla A, Schonberger K, Scognamiglio R, Altamura S, Florian CM, Fawaz M, Vonficht D, Tesio M, Collier P, Pavlinic D, Geiger H, Schroeder T, Benes V, Dick TP, Rieger MA, Stegle O, Trumpp A. Vitamin A-Retinoic Acid Signaling Regulates Hematopoietic Stem Cell Dormancy. Cell. 2017;169(5):807–23 e19. Epub 2017/05/10. doi: 10.1016/j.cell.2017.04.018. PubMed PMID: 28479188.

72. Yu W, Goncalves KA, Li S, Kishikawa H, Sun G, Yang H, Vanli N, Wu Y, Jiang Y, Hu MG, Friedel RH, Hu GF. Plexin-B2 Mediates Physiologic and Pathologic Functions of Angiogenin. Cell. 2017;171(4):849–64 e25. Epub 2017/11/04. doi: 10.1016/j.cell.2017.10.005. PubMed PMID: 29100074; PMCID: PMC5847377.

73. Kwok AJ, Allcock A, Ferreira RC, Cano-Gamez E, Smee M, Burnham KL, Zurke YX, Emergency Medicine Research O, McKechnie S, Mentzer AJ, Monaco C, Udalova IA, Hinds CJ, Todd JA, Davenport EE, Knight JC. Neutrophils and emergency granulopoiesis drive immune suppression and an extreme response endotype during sepsis. Nat Immunol. 2023;24(5):767–79. Epub 20230424. doi: 10.1038/s41590-023-01490-5. PubMed PMID: 37095375.

74. Zhang Y, Singh MK, Degenhardt KR, Lu MM, Bennett J, Yoshida Y, Epstein JA. Tie2Cre-mediated inactivation of plexinD1 results in congenital heart, vascular and skeletal defects. Dev Biol. 2009;325(1):82–93. Epub 2008/11/11. doi: 10.1016/j.ydbio.2008.09.031. PubMed PMID: 18992737; PMCID: PMC2650856.

75. Ding JB, Oh WJ, Sabatini BL, Gu C. Semaphorin 3E-Plexin-D1 signaling controls pathway-specific synapse formation in the striatum. Nat Neurosci. 2011;15(2):215–23. Epub 20111218. doi: 10.1038/nn.3003. PubMed PMID: 22179111; PMCID: PMC3267860.

76. Kim MH, Yang D, Kim M, Kim SY, Kim D, Kang SJ. A late-lineage murine neutrophil precursor population exhibits dynamic changes during demand-adapted granulopoiesis. Sci Rep. 2017;7:39804. Epub 20170106. doi: 10.1038/srep39804. PubMed PMID: 28059162; PMCID: PMC5216372.

77. Passegue E, Wagner EF, Weissman IL. JunB deficiency leads to a myeloproliferative disorder arising from hematopoietic stem cells. Cell. 2004;119(3):431–43. doi: 10.1016/j.cell.2004.10.010. PubMed PMID: 15507213.

78. Abram CL, Roberge GL, Hu Y, Lowell CA. Comparative analysis of the efficiency and specificity of myeloid-Cre deleting strains using ROSA-EYFP reporter mice. J Immunol Methods. 2014;408:89–100. Epub 20140522. doi: 10.1016/j.jim.2014.05.009. PubMed PMID: 24857755; PMCID: PMC4105345.

79. Dykstra B, Kent D, Bowie M, McCaffrey L, Hamilton M, Lyons K, Lee SJ, Brinkman R, Eaves C. Long-term propagation of distinct hematopoietic differentiation programs in vivo. Cell Stem Cell. 2007;1(2):218–29. doi: 10.1016/j.stem.2007.05.015. PubMed PMID: 18371352.

80. Meng Y, Carrelha J, Drissen R, Ren X, Zhang B, Gambardella A, Valletta S, Thongjuea S, Jacobsen SE, Nerlov C. Epigenetic programming defines haematopoietic stem cell fate restriction. Nat Cell Biol. 2023;25(6):812–22. Epub 20230501. doi: 10.1038/s41556-023-01137-5. PubMed PMID: 37127714.

81. Song Z, Park SH, Mu WC, Feng Y, Wang CL, Wang Y, Barthez M, Maruichi A, Guo J, Yang F, Lin AW, Heydari K, Chini CCS, Chini EN, Jang C, Chen D. An NAD(+)-dependent metabolic checkpoint regulates hematopoietic stem cell activation and aging. Nat Aging. 2024. Epub 20240723. doi: 10.1038/s43587-024-00670-8. PubMed PMID: 39044033.

82. Kwak HJ, Liu P, Bajrami B, Xu Y, Park SY, Nombela-Arrieta C, Mondal S, Sun Y, Zhu H, Chai L, Silberstein LE, Cheng T, Luo HR. Myeloid cell-derived reactive oxygen species externally regulate the proliferation of myeloid progenitors in emergency granulopoiesis. Immunity. 2015;42(1):159–71. Epub 20150108. doi: 10.1016/j.immuni.2014.12.017. PubMed PMID: 25579427; PMCID: PMC4303526.

83. Zhu H, Kwak HJ, Liu P, Bajrami B, Xu Y, Park SY, Nombela-Arrieta C, Mondal S, Kambara H, Yu H, Chai L, Silberstein LE, Cheng T, Luo HR. Reactive Oxygen Species-Producing Myeloid Cells Act as a Bone Marrow Niche for Sterile Inflammation-Induced Reactive Granulopoiesis. J Immunol. 2017;198(7):2854–64. Epub 20170224. doi: 10.4049/jimmunol.1602006. PubMed PMID: 28235862; PMCID: PMC5360524.

84. Wang L, Song G, Zheng Y, Tan W, Pan J, Zhao Y, Chang X. Expression of Semaphorin 4A and its potential role in rheumatoid arthritis. Arthritis Res Ther. 2015;17(1):227. Epub 20150825. doi: 10.1186/s13075-015-0734-y. PubMed PMID: 26303122; PMCID: PMC4549119.

85. Young K, Eudy E, Bell R, Loberg MA, Stearns T, Sharma D, Velten L, Haas S, Filippi MD, Trowbridge JJ. Decline in IGF1 in the bone marrow microenvironment initiates hematopoietic stem cell aging. Cell Stem Cell. 2021;28(8):1473–82 e7. Epub 20210412. doi: 10.1016/j.stem.2021.03.017. PubMed PMID: 33848471; PMCID: PMC8349778.

86. Chakkalakal JV, Jones KM, Basson MA, Brack AS. The aged niche disrupts muscle stem cell quiescence. Nature. 2012;490(7420):355-60. Epub 20120926. doi: 10.1038/nature11438. PubMed PMID: 23023126; PMCID: PMC3605795.

87. Arthur L, Esaulova E, Mogilenko DA, Tsurinov P, Burdess S, Laha A, Presti R, Goetz B, Watson MA, Goss CW, Gurnett CA, Mudd PA, Beers C, O’Halloran JA, Artyomov MN. Cellular and plasma proteomic determinants of COVID-19 and non-COVID-19 pulmonary diseases relative to healthy aging. Nat Aging. 2021;1(6):535–49. Epub 20210511. doi: 10.1038/s43587-021-00067-x. PubMed PMID: 37117829.

88. Choi YI, Duke-Cohan JS, Ahmed WB, Handley MA, Mann F, Epstein JA, Clayton LK, Reinherz EL. PlexinD1 glycoprotein controls migration of positively selected thymocytes into the medulla. Immunity. 2008;29(6):888–98. Epub 20081127. doi: 10.1016/j.immuni.2008.10.008. PubMed PMID: 19027330; PMCID: PMC2615553.

89. Dai X, Okon I, Liu Z, Wu Y, Zhu H, Song P, Zou MH. A novel role for myeloid cell-specific neuropilin 1 in mitigating sepsis. FASEB J. 2017;31(7):2881–92. Epub 20170321. doi: 10.1096/fj.201601238R. PubMed PMID: 28325756; PMCID: PMC5471517.

90. Zeng AGX, Nagree MS, Jakobsen NA, Shah S, Murison A, Cheong J-G, Turkalj S, Lim INX, Jin L, Araújo J, Aguilar-Navarro AG, Parris D, McLeod J, Kim H, Lee HS, Zhang L, Boulanger M, Wagenblast E, Flores-Figueroa E, Wang B, Schwartz GW, Shultz LD, Josefowicz SZ, Vyas P, Dick JE, Xie SZ. Identification of a human hematopoietic stem cell subset that retains memory of inflammatory stress. bioRxiv. 2023:2023.09.11.557271. doi: 10.1101/2023.09.11.557271.

