## Supplementary material for "Semaphorin 4A maintains functional diversity of the hematopoietic stem cell pool": Methods

### RESOURCE AVAILABILITY

#### Lead contact

#### Materials availability

This study did not generate new unique reagents.

#### Data and code availability

All data were analyzed with standard programs and packages, as detailed in Methods. Sequencing data from this study are available from ArrayExpress, E-MTAB-11359 and E-MTAB-12890.

### EXPERIMENTAL MODEL AND SUBJECT DETAILS

#### Animals

All animal experiments were approved by the Institutional Animal Care and Use Committee at Fred Hutchinson Cancer Center. Wild-type C57Bl/6J, B6SJL, Mx1-Cre and Mrp8-Cre mice were obtained from Jackson laboratory. PlxnD1 conditional KO mice and PlxnD1-GFP were obtained from Dr Chenghua Gu, Harvard University. Sema4AKO mice were obtained from Dr. A Kumanogoh, University of Osaka [1]. Sema4A conditional KO mice were obtained from Dr T Worzfeld, University of Marburg. Generation of mice carrying a targeted allele of Sema4A has been described previously [2]. These mice were crossed with Flp mice to obtain a floxed Sema4A allele.

To induce conditional deletion via Mx1-Cre, 6-8-week-old mice were interperitoneally (I.P.) injected with 100 μg/mouse high molecular weight polyinosine-polycytidylic acid (Poly(I:C) HMW, InvivoGen) once every other day for a total of three injections and analyzed 4 weeks after the last injection.

For the majority of the experiments, both male and female animals were used. Each cohort of animals was matched for age and sex. For acute LPS-induced inflammation experiments, only male mice were used. Young mice ranged from 8-12 weeks old and aged mice ranged from 74-80 weeks old. For the aging experiments, both male and female cohorts were generated but only the female cohort was analyzed in detail. Both WT and Sema4AKO mice were aged in the same environment.

#### PCR genotyping and excision validation for conditional alleles

Genomic DNA was extracted from peripheral blood samples using the DNeasy Blood & Tissue Kit (Qiagen) according to the manufacturer's instruction or using a fast extraction method involving incubation in ACK Lysing Buffer (Gibco) followed by resuspension in 50mM NaOH, incubation at 95C, and neutralization with 1M tris buffer (pH 7). Genotyping PCR primer sequences are listed in the reagents table.

To confirm Mx1-Cre-mediated excision of PlxnD1 "floxed" allele, up to 50,000 LKS ($lin^- c\text{-}Kit^+ Sca\text{-}1^+$) cells from animals of desired genotypes were sorted into 350µL RLT Plus Buffer (Qiagen). DNA and RNA were extracted with AllPrep DNA/RNA Micro Kit (Qiagen) according to the manufacturer's instructions. Primers listed in the reagent table were used to detect the excised and non-excised alleles by genomic DNA PCR. In addition, RNA was reverse transcribed using SuperScript IV VILO Master Mix (Invitrogen) and expression of PlxnD1 exon 1 relative to GAPDH was determined using PowerTrack™ SYBR Green Master Mix (Applied Biosystems) and qPCR primers listed in the reagent table.

**Sema4A qPCR**

To quantify Sema4A transcript levels in niche cell subsets and HSC, 50,000-80,000 CD45- cells, granulocytes, monocytes and lymphocytes and 2000-5000 HSC, were sorted into 350 µL of RLT Plus Buffer (Qiagen). RNA was extracted with AllPrep DNA/RNA Micro Kit (Qiagen) according to the manufacturer's instructions and was reverse transcribed using SuperScript IV VILO Master Mix (Invitrogen). Expression of Sema4A exon 1 relative to GAPDH was determined using PowerTrack™ SYBR Green Master Mix (Applied Biosystems) and qPCR primers listed in the reagent table.

**Histological analysis**

Dissected tissue samples were fixed in 4% formalin for 24 hours, decalcified in EDTA 2 weeks, and embedded in paraffin. Tissue sections were stained with hematoxylin/eosin.

**Flow cytometry and cell sorting**

Whole bone-marrow mononuclear cells (BMMNC) were collected by crushing tibias, femurs, and pelvis in Ca2+/Mg2+-free phosphate-buffered saline (D-PBS) supplemented with 2% fetal bovine serum (FBS, Fisher Scientific). For mature cells analysis, BMMNC were stained with fluorochrome-conjugated Mac1 (Invitrogen), Gr1 (BioLegend), B220 (BioLegend), and CD3ε (BioLegend) antibodies for 30 minutes. BMMNC were stained with the following fluorochrome-conjugated antibodies for 30 minutes to isolate neutrophils, monocytes and macrophages: CD11c (BioLegend), CD11b (BioLegend), Ly-6C (BioLegend), Siglec-F (BD Biosciences), CD24 (BioLegend), Ly-6G (BioLegend), I-A/I-E (BioLegend), CD3ε (BioLegend), CD19 (BioLegend) and NK1.1 (BioLegend). For hematopoietic stem and progenitor cell (HSPC) analysis, BMMNC were stained with fluorescently conjugated Sca-1 (BD Biosciences), c-Kit (BD Biosciences), Lineage cocktail (BD Biosciences), CD48 (BD Biosciences), CD34 (BD Biosciences), Flk2 (BD Biosciences), and CD150 (BioLegend) antibodies for 90 minutes. For cell sorting, BMMNC were lineage depleted by staining with CD3ε, CD11b, B220, TER-119, Gr-1, CD4 and CD8α biotin-conjugated antibodies (all from BD Biosciences) followed by application of streptavidin microbeads (Miltenyi Biotec) and depletion using magnetic separation columns (Miltenyi Biotec). Lineage-negative fraction was stained with conjugated monoclonal antibodies for HSPC markers as described above, except fluorescently conjugated streptavidin replaced fluorescently conjugated lineage cocktail. Samples were analyzed on BD FACSymphony A5 (BD Biosciences) or sorted on BD Aria II (BD Biosciences). All flow cytometry data were analyzed using FlowJo software. Graphs were made using Prism 9 (GraphPad) software or Microsoft Excel.

For flow cytometry of human bone marrow, cryopreserved cells were stained with fluorescent antibodies against CD45 (BD Biosciences), CD235α (BD Biosciences), CD15 (Caprico Biotechnologies), CD14 (Caprico Biotechnologies), CD16 (Caprico Biotechnologies), CD3 (Caprico Biotechnologies), CD20 (Caprico Biotechnologies), CD31(BD Biosciences), CD271 (BioLegend) and Sema4A (Invitrogen) as indicated and analyzed.

**Cell Cycle analysis**

For cell cycle analysis, we used our published protocol [3]. BMMNC were stained with conjugated monoclonal antibodies for HSPC markers, as described above. The cells were fixed and permeabilized using Cytofix/Cytoperm$^{TM}$ Fixation/Permeabilization kit (BD Biosciences) according to the manufacturer's instruction. Cells were then stained with fluorescently conjugated Ki67 antibody (1:20, BD Biosciences) for 45 minutes followed by DAPI (1:300, Invitrogen) for 10 minutes. Samples were analyzed at 2500-3000 threshold rate.

**BrdU incorporation**

For 5-bromo-2´-deoxyuridine (BrdU) incorporation, mice were first injected intraperitoneally with BrdU (Sigma-Aldrich) and were subsequently put on BrdU drinking water for 4 days prior to analysis, as previously described[4]. Cells were stained with conjugated monoclonal antibodies for HSPC markers, as described above. Cells were then fixed, permeabilized, and stained for BrdU using FITC BrdU Flow Kit (BD Biosciences) according to the manufacturer's instruction. All samples were analyzed using BD FACSymphony A5 (BD Biosciences).

**Transplantation experiments**

For all transplantation experiments, recipients were lethally irradiated 24 hours prior to transplant at 1200 cGy (a split dose of 600 cGy + 600 cGy with a 3 hr interval between doses) and maintained on Baytril-containing water for the first 4 weeks.

For competitive transplantation experiments involving young mice, 200-400 myHSC (lin$^-$c-Kit$^+$Sca-1$^+$CD48$^-$CD34$^-$Flk2$^-$CD150$^{high}$) or 200-400 lyHSC (lin$^-$c-Kit$^+$Sca-1$^+$CD48$^-$CD34$^-$Flk2$^-$CD150$^{low}$) from CD45.2 C57Bl/6J, Sema4AKO, PlxnD1$^{fl/fl}$ Mx1-Cre, and LPS treated models were co-injected with 200-250K CD45.1/2 competitor cells. For competitive transplantation experiments using HSC from aged mice, 2000 myHSC or 2000 lyHSC CD45.2 cells were transplanted with 200K CD45.1/2 competitor cells. Peripheral blood donor chimerism was monitored at 4-week intervals. Flow cytometry gating strategies for the identification of myHSC and lyHSC are shown in Extended Data Fig 1A.

For the non-competitive transplant experiments, the animals were lethally irradiated (as described above) and injected with 1100-1600 WT (CD45.1) myHSC or lyHSC. Engraftment kinetics were monitored by blood counts and flow cytometry every 4 weeks.

**Post-transplant chimerism analysis**

Peripheral blood was collected from each recipient via retro-orbital sinus, and complete blood count (CBC) was performed immediately after blood collection using Element HT5 (HESKA). Red blood cells were lysed in ACK Lysing Buffer (Gibco) and then stained with fluorescently conjugated CD45.1 (BioLegend), CD45.2 (BioLegend), Mac1 (Invitrogen), Gr1 (BioLegend), B220 (BioLegend), and CD3 (BioLegend) antibodies. Samples were analyzed using BD FACSymphony A5 (BD Biosciences).

**Induction of inflammatory response**

To induce chronic inflammation, wild-type C57Bl/6J and Sema4AKO mice were implanted with an intra-abdominal alzet osmotic pump (Braintree Scientific) that was loaded with lipopolysaccharide

(InvivoGen) resuspended in PBS. A dose of 8.4 mg/kg of LPS was administered over a 30-day period.

To induce acute inflammation, mice were injected intraperitoneally with 3.0-4.5 ug/g (depending on the biological activity of a specific batch which was established by titration) of lipopolysaccharide (InvivoGen) resuspended in PBS and euthanized at the point of maximal clinical response, which occurred at 72 hours post-injection. Bone marrow was analyzed on BD FACSymphony (BD Biosciences) or sorted on BD Aria II (BD Biosciences) for transplantation experiments.

**Intravital imaging**

MyHSC and lyHSC were isolated by flow sorting, as described above. Cells were stained with 10 μM DiD (Invitrogen) in D-PBS supplemented with 2% FBS for 20 minutes at 37°C. ~1500 DiD-stained myHSC and lyHSC were suspended in ~100 μl of $Ca^{2+}/Mg^{2+}$-free phosphate-buffered saline (D-PBS) and adoptively transferred via retro-orbital injection into anesthetized young WT and Sema4AKO mice. One day before transplantation, recipient mice were lethally irradiated (1200 cGy) using an x-ray irradiator (X-RAD 320, Precision) with a split dose of 600 cGy + 600 cGy with a 3 hr interval between doses.

For intravital imaging, mice were prepared as previously described [5]. Briefly, 14-15 hours after transplantation, the mice were anaesthetized with an induction dose of 3–4% isoflurane and a maintenance dose of 1.5–2% isoflurane. Mice were deemed anaesthetized by the toe pinch method. The hair on the calvarium was removed with a mechanical trimmer and the skin was cleaned with alcohol. The mice were mounted in a custom-designed heated mouse holder (for z-stack or time-lapse imaging). Next, a calvarial skin flap was created with a U-shaped incision to reveal the underlying calvaria as previously described. A drop of D-PBS was applied to the skull as the immersion fluid. The mice were transferred to the stage of a multiphoton/confocal laser-scanning video-rate microscope and an Olympus 25× 1.05 numerical aperture water-dipping objective was used for the imaging. Regions 3 and 4 of the calvarial bone marrow were imaged for ~3 hours per mouse and DiD cells were located and imaged [5, 6]. Z-stacks were acquired with 2 μm steps and time-lapse images were acquired at 10 min intervals for 90 min in 4-8 fields of view. The excitation wavelength from Insight X3 (Spectra-Physics) and Mai-Tai eHP (Spectra-Physics) lasers were set to 1040 nm for two-photon excitation of DiD (emission collected with a 659-700 nm bandpass filter) and second harmonic generation for bone imaging (emission collected with a 503-538 nm bandpass filter), and 820 nm for two-photon excitation of autofluorescence (emission collected with a 572-608 nm bandpass filter). After imaging was completed, mice were euthanized according to approved animal protocols. The contrast and brightness of images and videos were adjusted for display purposes only.

Intravital images were processed and analyzed using Fiji (ImageJ 1.53k). Representative images of single cells and clusters were created by taking Maximum Intensity Projections (MIPs) of z-stacks and adjusting the image contrast. The number of transplanted cells in the R3/R4 region of the calvaria was quantified and the distance from the transplanted cell's edge to the nearest endosteum surface was determined for each cell. Transplanted cells were classified as single or cluster cells when the nearest cell-to-cell edge distance was >15 μm or < 15 μm, respectively.

**Cell preparation for bulk RNA Sequencing**

For myHSC/lyHSC profiling, 100 cells from each fraction of interest were sorted into the lysis buffer - 10% Triton X-100 (Sigma-Aldrich), SUPERase-In RNase Inhibitor 20U/μl (Ambion) and snap-frozen. cDNA amplification was performed as per Smart-Seq2 protocol. Fragmentation and sample barcoding was performed using the NEBNext® Ultra™ II FS DNA Library Prep Kit for Illumina (New England

Biolabs) according to manufacturer's guidelines. Samples were sequenced on a 200 cycle NovaSeq SP.

**Cell preparation for Single-cell RNA sequencing**

*HSPC.* BMMNC were stained with conjugated monoclonal antibodies for HSPC markers as described above. 10,000 $Lin^{-}kit^{+}$ cells from young or aged WT/Sema4AKO mice were sorted in Ca2+/Mg2+-free phosphate-buffered saline (D-PBS) supplemented with 2% bovine serum albumin (New England Biolabs). Following one round of centrifugation, the cells were processed on 10x Genomics platform according to the manufacturer's protocol.

*MyHSC/lyHSC.* BMMNC were stained with conjugated monoclonal antibodies for HSPC markers as described above except for CD34 in order to minimize staining time and prevent RNA degradation (CD34 staining requires 90 minutes [7]. MyHSC and lyHSC were sorted into 96-well plates with the lysis buffer and stored frozen at -80. cDNA amplification and sequencing were performed as per Smart-Seq2 protocol [8].

**Bulk sequencing analysis**

RNA-Seq reads were aligned and counted using STAR [10] with the GRCm39 mouse genome assembly and Ensembl 106 [11] gene annotations. Differential expression was called using DESeq2 [12]. Genes were considered differentially expressed if their q-values (i.e., FDR adjusted p-value) were below 0.05. Gene set enrichment analysis was performed on the MSigDB mouse hallmark gene set [13]. For this analysis genes were ranked by the Z-test statistic calculated by DESeq2, where more extreme negative or positive values correspond to more significantly downregulated or upregulated genes, respectively. The ranked lists of genes were tested using the GSEAPreranked test [14] accessed through the GSEApy interface [15]. Significant gene sets were those with a q-value below 0.01.

**Single cell RNA-Seq sample processing and data analysis**

**10x Genomics**

**Library preparation**

Libraries from 10,000 Kit+ cells each from WT (n=1) and Sema4A KO (n=2) young and WT (n=2) old mice were prepared using the Single Cell 3′ Reagent Kit v3 (10x Genomics) according to the manufacturer's protocol.

**Quality control and data normalization**

The Cell Ranger 3.1.0 (10x Genomics) analysis pipeline was used to process the 10x single cell RNA-Seq output by aligning reads to mm10-3.0.0 mouse transcriptome (Ensembl). Cell Ranger was also used to aggregate condition replicates prior to processing. For all the analysis steps specified below, we used the SCANPY library [9], unless specified otherwise. As a quality control, first we filtered out cells with fewer than 200 genes detected with >0 reads, and we removed genes present in fewer than 3 cells. We also removed cells with a high percentage of reads mapped to mitochondrial genes (>5%) and a high number of detected genes (>6500). With the tool Scrublet [10], we predicted and removed doublets from the dataset.
After concatenating the data from WT and Sema4AKO mice, we normalized the data for sequencing depth to a target of 1e4 counts per cell. Then, after adding 1 as pseudocount, the count matrix was

log-transformed. Finally, the total counts per cell as well as the percentage of reads mapping to mitochondrial genes were regressed out.

#### Data clustering and visualization

We identified highly variable genes with the scanpy function “sc.pp.highly_variable_genes” (parameters: min_mean=0.0125, max_mean=3, min_disp=0.5) and we computed a neighborhood graph (n_neighbors = 8) on the first 40 principal components. To visualize the data, the graph was embedded in 2 dimensions using Uniform Manifold Approximation and Projection UMAP;[11].

We clustered the neighborhood graph using the Leiden clustering algorithm [12] with resolution 1 and 0.3 for young and old mice respectively. This resulted in 28 clusters in the young mice dataset and 17 clusters in the old mice dataset. The WT and Sema4AKO young dataset was subsetted to exclude small, isolated clusters and re-clustered with resolution 0.2. Cluster marker genes were found with a Wilcoxon rank-sum test. Based on known HSC markers (Ly6a, Procr, Hoxb5), we identified the cluster corresponding to HSC in the dataset from WT and Sema4AKO young mice (cluster 1), which was used for the enrichment analysis. In the WT old mice dataset, we similarly identified HSC and early multipotent progenitors (MPP) using marker genes and by predicted lineage relationships using graph abstraction [13], and, based on them, we built a differentiation trajectory for DPT analysis (see above).

#### Differential gene expression and enrichment analysis

We identified the genes differentially expressed between the WT and Sema4AKO cells in the HSC cluster 1 from young mice using a Wilcoxon test. For the pathway enrichment analysis, the top 500 upregulated and downregulated genes were ranked by statistical significance and the GSEA tool from the Broad Institute was used to perform enrichment analysis as described for bulk sequencing [14].

### Smart-Seq2 data analysis

#### Raw data processing and normalization

We quantified the abundance of the transcripts from 768 cells with salmon (v0.17) [15]. First, we indexed the mouse transcriptome (GRCm38) in quasi-mapping-based mode with --seqBias and --gcBias flags. Then, we aggregated the transcript level abundances into gene level abundances, which in turn was transformed into a gene-cell count matrix. Next, quality control was performed to filter out cells that satisfy any of the following criteria: 1) less than 4000 genes detected (detection threshold: reads-per-million>10); 2) overall mapping less than 50%; 3) fraction of reads mapping to mitochondrial transcripts larger than 0.02; 4) fraction of reads mapping to ERCC spike-ins higher than 0.01. After quality control, we retained 642 cells for downstream analyses (155 WT myHSC, 157 WT lyHSC, 166 KO myHSC, 164 KO lyHSC). The data were normalized using ‘quickcluster’ and ‘computeSumFactors’ functions from the scran package in R [16]. Finally, we added a pseudocount of 1 to the count matrix, followed by natural-log transformation.

#### Batch integration and visualization

Since the data were collected from two separate batches, we performed batch integration before visualizing the data. For this purpose, we used the Seurat batch integration workflow [17]. First, 3000 highly variable genes were selected from each batch. Then, we used the FindIntegrationAnchors function to estimate the anchors to use for the integration with 3000 features (anchor.features =3000) and the first 20 canonical variates (dims =1:20). Finally, the ‘IntegrateData’ function was used with default parameters to integrate the two batches.

To visualize the cells on a low dimensional space, we first built a k-nearest neighbor graph with 'neighbors' function from scanpy [9] with 10 principal components (PCs) and k=30. Then, the 'tl.umap' function was used to calculate a UMAP representation [18] and first two UMAP components were plotted. We applied this procedure separately for myHSC and lyHSC. To verify if myHSC and lyHSC are differentially affected by the absence of Sema4A, we calculated the pairwise Spearman's correlation distance (defined as (1-ρ)2, where ρ is the Spearman's correlation coefficient computed on the top 3000 highly variable genes identified with Seurat) between WT and KO cells for myHSCs and lyHSC separately. Then, we tested the statistical significance of the difference between the two distributions of pairwise distances by using the Wilcoxon rank-sum test.

**Diffusion pseudotime analysis**

For this analysis, we first generated 10x Genomics single cell RNA-Seq profiles of $lin^-c\text{-}Kit^+$ HSPC from 74-weeks old WT animals, i.e. age-matched with WT/Sema4AKO animals for the Smart-Seq2 single-cell RNA-Seq experiment described above. In this 10x dataset, we used previously described markers to map the clusters corresponding to HSC (*Ly6a, Procr, Hlf*) and MPP (*Cd34, Cebpa, Ctsg*) [28] and utilized the transcriptomes of cells within these clusters to estimate a differentiation trajectory, in which higher DPT values correspond to more mature cells (data not shown). For this purpose, a k-nearest neighbor graph was first built with the first 5 PCs and k=15. A diffusion map was then constructed with the 'tl.diffmap' function from scanpy. We defined a diffusion pseudotime (DPT) by selecting the root cell that had the lowest value of the first diffusion component [29]. As expected, analysis of known self-renewing marker genes revealed downregulation of vWF, *Mpl*, *Fdg5*, C*tnnal1*, *Procr*, and upregulation of *Ctsg* and *Cbpa* as cells progressed from HSC to MPP (data not shown). Finally, we used these differentiation trajectories in order to test the difference in the distribution of DPT values for both WT and KO myHSCs, which we estimated with the Wilcoxon rank-sum test.

**Differential gene expression analysis**

We found differentially expressed genes between KO and WT with the DEseq2 package [19] on R. First, we removed genes that were expressed in fewer than 10 cells for each batch, and the counts were rounded to integers. Then, we created a DESeq object from the count matrix with the design ~condition+batch. Fold-changes and p-values were estimated using the DESeq function with default parameters.

**ATAC-seq analysis**

All sequencing reads were trimmed using cutadapt [20], and trimmed reads (>36 bp minimum alignment length) were mapped against the mm10 genome using BWA aligner [21]. We used de-duplicated and uniquely mapped reads for peak calling analysis after excluding high-sensitive black-list regions defined by ENCODE. The candidate peaks were predicted by MACS peak calling software (FDR < 0.05) [22]. After identifying narrow peaks from biological replicates, we created a merged set of consensus peaks and generated a matrix of open chromatin regions (OCRs). Then, we calculated the number of mapped reads located at the center of the peaks (+/- 250bp from mid-point).This OCR matrix was then imported into the R package DESeq2 [23], and we determined differentially accessible regions (DARs) with cutoffs: 1) FC > 1.5, CPM > 1.0, p-value (stringent cutoffs), 2) FC > 1.2, CPM > 1.0, p-value < 0.01 (relaxed cutoffs). We identified 2,379 gain-of-accessible peaks (enriched in MUL) and 2,028 loss of accessible peaks (enriched in PLT) for Nerlov group's ATAC-seq datasets (stringent cutoff). We also identified 215 gain of accessible peaks (enriched in Sema4aKO) and 112 loss of accessible peaks (enriched in WT) from our myHSC ATAC-seq datasets (relaxed cutoffs). Finally, the candidate differential open chromatin regions were submitted to search for potential transcription factor binding sites using HOMER software [24] with all open chromatin peaks as background regions. In this analysis, de novo motif (unbiased way to find motifs using various k-mers) and known motif searches using

HOMER known motif database were performed, and we reported the top denovo motif or known motif results. CREB5 and RUNX3 motifs were identified from both denovo and known motif analysis in Sema4AKO enriched DARs. RUNX3 was second highest motifs from known motif.

**Extended Data Table 1**

Summary of the RNAseq and ATAC-Seq data (List of differentially expressed genes).

**Extended Data Table 2**

Complied list of blood counts.

### METHOD REFERENCES


1. Kumanogoh, A., et al., *Nonredundant roles of Sema4A in the immune system: defective T cell priming and Th1/Th2 regulation in Sema4A-deficient mice.* Immunity, 2005. **22**(3): p. 305-16.
2. Xia, J., et al., *Semaphorin-Plexin Signaling Controls Mitotic Spindle Orientation during Epithelial Morphogenesis and Repair.* Dev Cell, 2015. **33**(3): p. 299-313.
3. Galvin, A., et al., *Cell Cycle Analysis of Hematopoietic Stem and Progenitor Cells by Multicolor Flow Cytometry.* Curr Protoc Cytom, 2019. **87**(1): p. e50.
4. Feng, C.G., et al., *The p47 GTPase Lrg-47 (Irgm1) links host defense and hematopoietic stem cell proliferation.* Cell Stem Cell, 2008. **2**(1): p. 83-9.
5. Christodoulou, C., et al., *Live-animal imaging of native haematopoietic stem and progenitor cells.* Nature, 2020. **578**(7794): p. 278-283.
6. Sipkins, D.A., et al., *In vivo imaging of specialized bone marrow endothelial microdomains for tumour engraftment.* Nature, 2005. **435**(7044): p. 969-73.
7. Okamoto, K., et al., *Self-organization of all-inorganic dodecatungstophosphate nanocrystallites.* J Am Chem Soc, 2007. **129**(23): p. 7378-84.
8. Picelli, S., et al., *Smart-seq2 for sensitive full-length transcriptome profiling in single cells.* Nat Methods, 2013. **10**(11): p. 1096-8.
9. Wolf, F.A., P. Angerer, and F.J. Theis, *SCANPY: large-scale single-cell gene expression data analysis.* Genome Biol, 2018. **19**(1): p. 15.
10. Wolock, S.L., R. Lopez, and A.M. Klein, *Scrublet: Computational Identification of Cell Doublets in Single-Cell Transcriptomic Data.* Cell Syst, 2019. **8**(4): p. 281-291 e9.
11. McInnes, L., J. Healy, and J. Melville, *UMAP: Uniform Manifold Approximation and Projection for Dimension Reduction.* arXiv, 2020. **1802.03426**.
12. Traag, V.A., L. Waltman, and N.J. van Eck, *From Louvain to Leiden: guaranteeing well-connected communities.* Sci Rep, 2019. **9**(1): p. 5233.
13. Wolf, F.A., et al., *PAGA: graph abstraction reconciles clustering with trajectory inference through a topology preserving map of single cells.* Genome Biol, 2019. **20**(1): p. 59.
14. Subramanian, A., et al., *Gene set enrichment analysis: a knowledge-based approach for interpreting genome-wide expression profiles.* Proc Natl Acad Sci U S A, 2005. **102**(43): p. 15545-50.
15. Patro, R., et al., *Salmon provides fast and bias-aware quantification of transcript expression.* Nat Methods, 2017. **14**(4): p. 417-419.
16. Lun, A.T., K. Bach, and J.C. Marioni, *Pooling across cells to normalize single-cell RNA sequencing data with many zero counts.* Genome Biol, 2016. **17**: p. 75.
17. Butler, A., et al., *Integrating single-cell transcriptomic data across different conditions, technologies, and species.* Nat Biotechnol, 2018. **36**(5): p. 411-420.

18. Becht, E., et al., *Dimensionality reduction for visualizing single-cell data using UMAP.* Nat Biotechnol, 2018.
19. Love, M.I., W. Huber, and S. Anders, *Moderated estimation of fold change and dispersion for RNA-seq data with DESeq2.* Genome Biol, 2014. **15**(12): p. 550.
20. Martin M. Cutadapt Removes Adapter Sequences From High-Throughput Sequencing Reads. *EMBnetjournal*. 2011;17. doi: http://dx.doi.org/10.14806/ej.17.1.200
21. Li H, Durbin R. Fast and accurate short read alignment with Burrows-Wheeler transform. *Bioinformatics*. 2009;25:1754-1760. doi: 10.1093/bioinformatics/btp324
22. Zhang Y, Liu T, Meyer CA, Eeckhoute J, Johnson DS, Bernstein BE, Nusbaum C, Myers RM, Brown M, Li W, et al. Model-based analysis of ChIP-Seq (MACS). *Genome Biol*. 2008;9:R137. doi: 10.1186/gb-2008-9-9-r137
23. Love MI, Huber W, Anders S. Moderated estimation of fold change and dispersion for RNA-seq data with DESeq2. *Genome Biol*. 2014;15:550. doi: 10.1186/s13059-014-0550-8
24. Heinz S, Benner C, Spann N, Bertolino E, Lin YC, Laslo P, Cheng JX, Murre C, Singh H, Glass CK. Simple combinations of lineage-determining transcription factors prime cis-regulatory elements required for macrophage and B cell identities. *Mol Cell*. 2010;38:576-589. doi: 10.1016/j.molcel.2010.05.004
