## Supplementary material for "Semaphorin 4A maintains functional diversity of the hematopoietic stem cell pool": Reagent List

| REAGENT or RESOURCE | SOURCE | IDENTIFIER |
| --- | --- | --- |
| Antibodies | | |
| Biotin anti-mouse CD8a | BD Biosciences | Cat# 553029; RRID: AB_394567 |
| Biotin anti-mouse CD3ε | BD Biosciences | Cat# 553060; RRID: AB_394593 |
| Biotin anti-mouse B220 | BD Biosciences | Cat# 553086; RRID: AB_394616 |
| Biotin anti-mouse CD4 | BD Biosciences | Cat# 553728; RRID: AB_395012 |
| Biotin anti-CD11b | BD Biosciences | Cat# 553309; RRID: AB_394773 |
| Biotin anti-mouse TER-119 | BD Biosciences | Cat# 553672; RRID: AB_394985 |
| Biotin anti-mouse Ly-6G and Ly-6C (Gr-1) | BD Biosciences | Cat# 553125; RRID: AB_394641 |
| AF700 anti-mouse CD11b (Mac1) | Invitrogen | Cat# 56-0112-82; RRID: AB_65758 |
| BV570 anti-mouse Gr-1 | BioLegend | Cat# 108431; RRID: AB_10896783 |
| BV785 anti-mouse/human B220 | BioLegend | Cat# 103245; RRID: AB_11218795 |
| FITC anti-mouse B220 | BD Biosciences | Cat# 553087; RRID: AB_394617 |
| APC anti-mouse CD3ε | BioLegend | Cat# 100312; RRID: AB_312677 |
| FITC anti-mouse CD45.2 | BioLegend | Cat# 109806; RRID: AB_313443 |
| BV421 anti-mouse CD45.1 | BioLegend | Cat# 110732; RRID: AB_2562563 |
| PE-CF594 anti-mouse Ly-6A/E (Sca-1) | BD Biosciences | Cat# 562730; RRID: AB_2737751 |
| BUV395 anti-mouse Ly-6A/E (Sca-1) | BD Biosciences | Cat# 566216; RRID: AB_2739606 |
| APC anti-mouse CD117 (c-Kit) | BD Biosciences | Cat# 553356; RRID: AB_398536 |
| PE anti-mouse CD117 (c-Kit) | BD Biosciences | Cat# 561075; RRID: AB_10563204 |
| PerCP-Cy 5.5 mouse Lineage antibody cocktail | BD Biosciences | Cat# 561317; RRID: AB_10612020 |
| APC-Cy7 anti-mouse CD48 | BD Biosciences | Cat# 561242; RRID: AB_10644381 |
| PE/Cyanine7 anti-mouse CD150 (SLAM) | BioLegend | Cat# 115914; RRID: AB_439797 |
| FITC anti-mouse CD34 | BD Biosciences | Cat# 553733; RRID: AB_395017 |
| AF700 anti-mouse CD34 | BD Biosciences | Cat# 560518; RRID: AB_1727471 |
| BV421 anti-mouse CD135 (Flk-2) | BD Biosciences | Cat# 566292; RRID: AB_2739665 |
| PE anti-mouse CD135 (Flk-2) | BD Biosciences | Cat# 553842; RRID: AB_395079 |
| APC-Cy7 anti-mouse CD16/CD32 | BD Biosciences | Cat# 560541 |
| BV510 anti-mouse CD41 | BD Biosciences | Cat# 740136; RRID: AB_2739892 |
| BV786 anti-mouse CD105 | BD Biosciences | Cat# 564746; RRID: AB_2732065 |
| FITC anti-mouse CD127 | BioLegend | Cat# 135007; RRID: AB_1937231 |

| | | |
|---|---|---|
| AF700 mouse anti-Ki-67 | BD Biosciences | Cat# 561277; RRID: AB_10611571 |
| PE anti-mouse CD11c | BioLegend | Cat# 117307; RRID: AB_313776 |
| BV605 anti-mouse CD11b | BioLegend | Cat# 101257; RRID: AB_2565431 |
| BV711 anti-mouse Ly-6C | BioLegend | Cat# 128037; RRID: AB_2562630 |
| AF647 anti-mouse Siglec-F | BD Biosciences | Cat# 562680 ; RRID: AB_2687570 |
| PE/Cyanine7 anti-mouse CD24 | BioLegend | Cat# 101822 ; RRID: AB_756048 |
| BV785 anti-mouse Ly-6G | BioLegend | Cat# 127645 ; RRID: AB_2566317 |
| AF700 anti-mouse I-A/I-E | BioLegend | Cat# 107622 ; RRID: AB_493727 |
| APC/Fire™ 750 anti-mouse CD3ε | BioLegend | Cat# 100362 ; RRID: AB_2629687 |
| APC/Fire™ 750 anti-mouse CD19 | BioLegend | Cat# 115558 ; RRID: AB_2572120 |
| APC/Fire™ 750 anti-mouse NK-1.1 | BioLegend | Cat# 108752 ; RRID: AB_2629764 |
| FITC anti-mouse CD45 | BioLegend | Cat# 103108; RRID: AB_312973 |
| AF700 anti-mouse Plexin B2 | R&D Systems | Cat#; FAB6836N |
| AF700 anti-human CD45 | BD Biosciences | Cat# 560566; RRID: AB_1645452 |
| BV786 anti-human CD235α | BD Biosciences | Cat# 740984; RRID: AB_2740608 |
| PE anti-human CD15 | Caprico Biotechnologies | Cat# 105026 |
| PE/Cyanine7 anti-human CD14 | Caprico Biotechnologies | Cat# 103486 |
| APC-Cy7 anti-human CD16 | Caprico Biotechnologies | Cat# 101496 |
| BV650 anti-human CD3 | Invitrogen | Cat# 416-0038-42; RRID: AB_2921039 |
| APC anti-human CD20 | Caprico Biotechnologies | Cat# 103746 |
| APC-Cy7 anti-human CD31 | BD Biosciences | Cat# 563653; RRID: AB_2738350 |
| PE anti-human CD271 (NGFR) | BioLegend | Cat# 345105; RRID: AB_2282827 |
| PerCP-Cy 5.5 anti-mouse/human Sema4A | Invitrogen | Cat#; 46-975-341 RRID: AB_2573898 |
| Chemicals, peptides, and recombinant proteins | | |
| Cytiva HyClone™ Fetal Bovine Serum (Canada), Characterized | Fisher Scientific | Cat# SH3039603HI |
| BSA, Molecular Biology Grade | New England BioLabs | Cat# B9200 |
| 10 % Triton X-100 | Sigma-Aldrich | Cat# 934433 |
| StemSpan™ SFEM II | Stem Cell Technologies | Cat# 09655 |
| ACK Lysing Buffer | Gibco | Cat# A1049201 |
| Critical commercial assays | | |
| DAPI | Invitrogen | Cat# 62248 |
| PerCP-Cy 5.5 Streptavidin | BD Biosciences | Cat# 551419 |
| Click-iT™ Plus EdU Alexa Fluor™ 488 Flow Cytometry Assay Kit | Invitrogen | Cat# C10632 |
| BD Pharmingen™ APC BrDU Kit | BD Biosciences | Cat# 552598 |
| 5-Bromo-2′-deoxyuridine | Sigma-Aldrich | Cat# B5002-5G |

| | | |
|---|---|---|
| BD Cytofix/Cytoperm Fixation/Permeabilization Solution Kit | BD Biosciences | Cat# 554714 |
| Invitrogen UltraComp eBeads Compensation Beads | Invitrogen | Cat# 50-112-9040 |
| Rainbow Calibration Particles, 8 peaks | Spherotech | Cat# RCP-30-5A |
| Streptavidin MicroBeads | Miltenyi Biotec | Cat# 130-048-101 |
| Poly(I:C) HMW | InvivoGen | Cat# tlrl-pic-5 |
| Lipopolysaccharide from E. coli 0111:B4 | InvivoGen | Cat# tlrl-eblps |
| NEBNext® Ultra™ II FS DNA Library Prep Kit for Illumina | New England BioLabs | Cat# E7805 |
| NEBNext® Multiplex Oligos for Illumina® (Index Primers Set 1) | New England BioLabs | Cat# E7335 |
| SUPERase-In RNase Inhibitor | Ambion | Cat# AM2694 |
| dNTP mix | Thermo Scientific | Cat# 10319879 |
| Maxima H minus Reverse Transcriptase | Thermo Scientific | Cat# EP0752 |
| Terra PCR Direct Polymerase Mix | Takara Bio USA, | Cat# 639270 |
| Agencourt AMPure XP beads | Beckman Coulter | Cat# A63881 |
| EB solution | Qiagen | Cat# 19086 |
| PowerUp SYBR Green Master Mix | Applied Biosystems | Cat# A25742 |
| AllPrep DNA/RNA Micro Kit | Qiagen | Cat# 80284 |
| SuperScript IV VILO Master Mix | Invitrogen | Cat# 11756050 |
| Platinum PCR SuperMix High Fidelity | Invitrogen | Cat# 12532016 |
| LD Columns | Miltenyi Biotec | Cat# 130-042-901 |
| DNeasy Blood & Tissue Kit | Qiagen | Cat# 69506 |
| Deposited data | | |
| NCBI GEO / ArrayExpress | ArrayExpress | Accession: E-MTAB-11359 |
| NCBI GEO / ArrayExpress | ArrayExpress | Accession: E-MTAB-12890 |
| Experimental models: mouse strains | | |
| B6.SJL-Ptprca Pepcb/BoyJ | The Jackson Laboratory | RRID: IMSR_JAX:002014 |
| C57BL/6J | The Jackson Laboratory | RRID: IMSR_JAX:000664 |
| STOCK Tg(Mx1-cre)1Cgn/J | The Jackson Laboratory | RRID: IMSR_JAX:002527 |
| B6.Cg-Tg(S100A8-cre,-EGFP)1Ilw/J | The Jackson Laboratory | RRID: IMSR_JAX:021614 |
| Sema4AKO strain | Dr. A Kumanogoh, University of Osaka | N/A |
| PlxnD1 conditional KO mice | Dr Chenghua Gu, Harvard University | N/A |
| PlxnD1-GFP | Dr Chenghua Gu, Harvard University | N/A |
| Sema4A conditional KO mice | Dr T Worzfeld, University of Marburg | N/A |
| Oligonucleotides | | |
| Forward primer for Sema4AKO allele genotyping 5'- GTTTCCTCAGAACCATCTGGTGACCATCTC -3' | (Kumanogoh et al., 2005) | N/A |
| Reverse primer for Sema4AKO allele genotyping 5'- TCACCATGTTCTTTAGCCTTGGGATTC -3' | (Kumanogoh *et al.*, 2005) | N/A |
| Forward primer for PlxnD1 floxed allele genotyping 5'- ACAGGTGTGTGCTCAAGGCCAC -3' | (Wang et al., 2015) | N/A |
| Reverse primer for PlxnD1 floxed allele genotyping 5'- CAGCCCTATAGTTCTCCACCAAAGA -3' | (Wang *et al.*, 2015) | N/A |
| Forward primer for Mx1-Cre genotyping 5'- GTGAGTTTCGTTTCTGAGCTCC -3' | Jackson Laboratory | N/A |

| | | |
|---|---|---|
| Reverse primer for Mx1-Cre genotyping 5'- CGGTTATTCAACTTGCACCA -3' | Jackson Laboratory | N/A |
| Forward primer for detection of PlxnD1 excised allele 5'- GGCCAGGAACCAGGAAAGAGG -3' | This paper | N/A |
| Alternative forward primer for detection of PlxnD1 floxed allele 5'- GTGTGTGCTCAAGGCCACCTC -3' | This paper | N/A |
| Reverse primer for detection of PlxnD1 excised and floxed alleles 5'- CAGCCCTATAGTTCTCCACCAAAGA -3' | This paper | N/A |
| Forward qPCR primer for GAPDH detection 5'- ATGAATACGGCTACAGCAACAGG -3' | (Kurimoto et al., 2006) | N/A |
| Reverse qPCR primer for GAPDH detection 5'- CTCTTGCTCAGTGTCCTTGCTG -3' | (Kurimoto *et al.*, 2006) | N/A |
| Forward qPCR primer for PlxnD1 exon 1 detection 5'- CGCAACCGTAGCCTAGAAGAC -3' | Massachusetts General Hospital Primer Bank | ID: 153792703c1 |
| Reverse qPCR primer for PlxnD1 exon 1 detection 5'- GGTTAAGGTCGAAGGTGAAGAG -3' | Massachusetts General Hospital Primer Bank | ID: 153792703c1 |
| Forward qPCR primer for Sema4A exon 1 detection 5'- ATGGAGTCTCCTGCGTGTTT -3' | (Sun et al., 2017) | N/A |
| Reverse qPCR primer for Sema4A exon 1 detection 5'- GAAGCAGGTGGCAGTGATG -3' | (Sun *et al.*, 2017) | N/A |
| Template Switching Oligo for bulk RNA-Seq 5′-AAGCAGTGGTATCAACGCAGAGTACATrGrG+G-3′ | (Picelli et al., 2014) | N/A |
| Oligo-dT30VN for bulk RNA-Seq 5′–AAGCAGTGGTATCAACGCAGAGTAC(T30)VN-3′ | (Picelli *et al.*, 2014) | N/A |
| ISPCR oligo for bulk RNA-Seq 5′-AAGCAGTGGTATCAACGCAGAGT-3′ | (Picelli *et al.*, 2014) | N/A |
| Software and algorithms | | |
| R (R-3.2.3 – R-3.6.1) | The R Foundation; https://www.r-project.org | RRID: SCR_001905 |
| FlowJo Software | https://www.flowjo.com/solutions/flowjo | RRID: SCR_008520 |
| GraphPad Prism 9.0 | GraphPad Software | RRID: SCR_002798 |
| Biorender | https://biorender.com | RRID: SCR_018361 |
| Fiji (ImageJ 1.53k) | https://imagej.net/software/fiji/ | RRID: SCR_002285 |
| Cell Ranger 3.1.0 | 10x Genomics | RRID: SCR_017344 |
| Seurat | (Stuart et al., 2019) | N/A |
| FastQC | (Andrews, 2010) | N/A |
| Flexbar | (Dodt et al., 2012) | N/A |
| STAR | (Dobin et al., 2013) | N/A |
| SAMtools | (Li et al., 2009) | N/A |
| Multicov, BEDtools | (Quinlan and Hall, 2010) | N/A |
| DESeq2 | (Love et al., 2014) | N/A |
| ggplot2 | (Wickham, 2016) https://ggplot2.tidyverse.org/ | RRID:SCR_014601 |
| GSEA software | (Subramanian et al., 2005) Broad Institute | RRID:SCR_003199 |
| Cytoscape software | (Shannon et al., 2003) | RRID: SCR_003032 |
| EnrichmentMap | https://www.baderlab.org/Software/EnrichmentMap | RRID: SCR_016052 |
| SCANPY | (Wolf et al., 2018) | N/A |
| Partition-based graph abstraction (PAGA) | (Wolf et al., 2019) | N/A |
| Salmon v0.17 | (Patro et al., 2017) | N/A |
| Scran | (Lun et al., 2016) | RRID: SCR_016944 |
| Cyclone | (Scialdone et al., 2015) | N/A |

| | | |
|---|---|---|
| Scrublet | (Wolock et al., 2019) | N/A |
| Other | | |
| Alzet Osmotic Pump | | Cat# NC0059919 |

**REAGENT REFERENCES**


Andrews, S. (2010). FastQC: A Quality Control Tool for High Throughput Sequence Data [online]. http://www.bioinformatics.babraham.ac.uk/projects/fastqc/

Dobin, A., Davis, C.A., Schlesinger, F., Drenkow, J., Zaleski, C., Jha, S., Batut, P., Chaisson, M., and Gingeras, T.R. (2013). STAR: ultrafast universal RNA-seq aligner. Bioinformatics *29*, 15-21. 10.1093/bioinformatics/bts635.

Dodt, M., Roehr, J.T., Ahmed, R., and Dieterich, C. (2012). FLEXBAR-Flexible Barcode and Adapter Processing for Next-Generation Sequencing Platforms. Biology (Basel) *1*, 895-905. 10.3390/biology1030895.

Guo, P., Poulos, M.G., Palikuqi, B., Badwe, C.R., Lis, R., Kunar, B., Ding, B.S., Rabbany, S.Y., Shido, K., Butler, J.M., and Rafii, S. (2017). Endothelial jagged-2 sustains hematopoietic stem and progenitor reconstitution after myelosuppression. J Clin Invest *127*, 4242-4256. 10.1172/JCI92309.

Kumanogoh, A., Shikina, T., Suzuki, K., Uematsu, S., Yukawa, K., Kashiwamura, S., Tsutsui, H., Yamamoto, M., Takamatsu, H., Ko-Mitamura, E.P., et al. (2005). Nonredundant roles of Sema4A in the immune system: defective T cell priming and Th1/Th2 regulation in Sema4A-deficient mice. Immunity *22*, 305-316. 10.1016/j.immuni.2005.01.014.

Kurimoto, K., Yabuta, Y., Ohinata, Y., Ono, Y., Uno, K.D., Yamada, R.G., Ueda, H.R., and Saitou, M. (2006). An improved single-cell cDNA amplification method for efficient high-density oligonucleotide microarray analysis. Nucleic Acids Res *34*, e42. 10.1093/nar/gkl050.

Li, H., Handsaker, B., Wysoker, A., Fennell, T., Ruan, J., Homer, N., Marth, G., Abecasis, G., Durbin, R., and Genome Project Data Processing, S. (2009). The Sequence Alignment/Map format and SAMtools. Bioinformatics *25*, 2078-2079. 10.1093/bioinformatics/btp352.

Love, M.I., Huber, W., and Anders, S. (2014). Moderated estimation of fold change and dispersion for RNA-seq data with DESeq2. Genome Biol *15*, 550. 10.1186/s13059-014-0550-8.

Lun, A.T., Bach, K., and Marioni, J.C. (2016). Pooling across cells to normalize single-cell RNA sequencing data with many zero counts. Genome Biol *17*, 75. 10.1186/s13059-016-0947-7.

Patro, R., Duggal, G., Love, M.I., Irizarry, R.A., and Kingsford, C. (2017). Salmon provides fast and bias-aware quantification of transcript expression. Nat Methods *14*, 417-419. 10.1038/nmeth.4197.

Picelli, S., Faridani, O.R., Bjorklund, A.K., Winberg, G., Sagasser, S., and Sandberg, R. (2014). Full-length RNA-seq from single cells using Smart-seq2. Nat Protoc *9*, 171-181. 10.1038/nprot.2014.006.

Quinlan, A.R., and Hall, I.M. (2010). BEDTools: a flexible suite of utilities for comparing genomic features. Bioinformatics *26*, 841-842. 10.1093/bioinformatics/btq033.

Scialdone, A., Natarajan, K.N., Saraiva, L.R., Proserpio, V., Teichmann, S.A., Stegle, O., Marioni, J.C., and Buettner, F. (2015). Computational assignment of cell-cycle stage from single-cell transcriptome data. Methods *85*, 54-61. 10.1016/j.ymeth.2015.06.021.

Shannon, P., Markiel, A., Ozier, O., Baliga, N.S., Wang, J.T., Ramage, D., Amin, N., Schwikowski, B., and Ideker, T. (2003). Cytoscape: a software environment for integrated models of biomolecular interaction networks. Genome Res *13*, 2498-2504. 10.1101/gr.1239303.

Stuart, T., Butler, A., Hoffman, P., Hafemeister, C., Papalexi, E., Mauck, W.M., 3rd, Hao, Y., Stoeckius, M., Smibert, P., and Satija, R. (2019). Comprehensive Integration of Single-Cell Data. Cell *177*, 1888-1902 e1821. 10.1016/j.cell.2019.05.031.

Subramanian, A., Tamayo, P., Mootha, V.K., Mukherjee, S., Ebert, B.L., Gillette, M.A., Paulovich, A., Pomeroy, S.L., Golub, T.R., Lander, E.S., and Mesirov, J.P. (2005). Gene set enrichment analysis: a knowledge-based approach for interpreting genome-wide expression profiles. Proc Natl Acad Sci U S A *102*, 15545-15550. 10.1073/pnas.0506580102.

Sun, T., Yang, L., Kaur, H., Pestel, J., Looso, M., Nolte, H., Krasel, C., Heil, D., Krishnan, R.K., Santoni, M.J., et al. (2017). A reverse signaling pathway downstream of Sema4A controls cell migration via Scrib. J Cell Biol *216*, 199-215. 10.1083/jcb.201602002.

Wang, F., Eagleson, K.L., and Levitt, P. (2015). Positive regulation of neocortical synapse formation by the Plexin-D1 receptor. Brain Res *1616*, 157-165. 10.1016/j.brainres.2015.05.005.

Wickham, H. (2016). ggplot2: Elegant Graphics for Data Analysis (Springer-Verlag New York).

Wolf, F.A., Angerer, P., and Theis, F.J. (2018). SCANPY: large-scale single-cell gene expression data analysis. Genome Biol *19*, 15. 10.1186/s13059-017-1382-0.

Wolf, F.A., Hamey, F.K., Plass, M., Solana, J., Dahlin, J.S., Gottgens, B., Rajewsky, N., Simon, L., and Theis, F.J. (2019). PAGA: graph abstraction reconciles clustering with trajectory inference through a topology preserving map of single cells. Genome Biol *20*, 59. 10.1186/s13059-019-1663-x.

Wolock, S.L., Lopez, R., and Klein, A.M. (2019). Scrublet: Computational Identification of Cell Doublets in Single-Cell Transcriptomic Data. Cell Syst *8*, 281-291 e289. 10.1016/j.cels.2018.11.005.
